# Spatially-resolved molecular sex differences at single cell resolution in the adult human ventromedial and arcuate hypothalamus

**DOI:** 10.1101/2024.12.07.627362

**Authors:** Bernard Mulvey, Yi Wang, Heena R. Divecha, Svitlana V. Bach, Kelsey D. Montgomery, Sophia Cinquemani, Atharv Chandra, Yufeng Du, Ryan A. Miller, Joel E. Kleinman, Stephanie C. Page, Thomas M. Hyde, Keri Martinowich, Stephanie C. Hicks, Kasper D. Hansen, Kristen R. Maynard

## Abstract

The hypothalamus contains multiple regions, including the ventromedial hypothalamus (VMH) and arcuate (ARC), which are responsible for sex-differentiated functions such as endocrine signaling, metabolism, and reproductive behaviors. While molecular, anatomic, and sex-differentiated features of rodent hypothalamus are well-established, much less is known about these regions in humans. Here we provide a spatially-resolved single cell atlas of sex-differentially expressed (sex-DE) genes in human VMH and ARC. We identify neuronal populations governing hypothalamus-specific functions, define their spatial distributions, and show increased retinoid pathway gene expression compared to rodents. Within VMH and ARC, we find correlated autosomal expression differences localized to *ESR1/TAC3*-expressing and *CRHR2*-expressing neurons, and extensive sex-DE of genes linked to sex-biased disorders including autism, depression, and schizophrenia. Our molecular mapping of disease associations to hypothalamic cell types with established roles in sex-divergent physiology and behavior provides insights into mechanistic bases of sex bias in neurodevelopmental and neuropsychiatric disorders.

## Introduction

The hypothalamus (HYP) is a specialized brain structure that modulates behavioral and physiological drives essential to survival–ranging from social behavior (*1*) to metabolism (*2*) (**Fig. 1A**). This functional breadth is mediated by different HYP regions (‘nuclei’) with distinct circuitry, neurotransmitters, and neuropeptides. The tuberal portion of HYP contains the ventromedial hypothalamus (VMH) and arcuate (ARC), two nuclei that in model species show sex-differentiated development (*2*) and sex divergence at the levels of behavior (*3–5*), physiology (*6–8*), cell types (*9*, *10*), and gene expression (*8*, *11*).

**Fig. 1.**
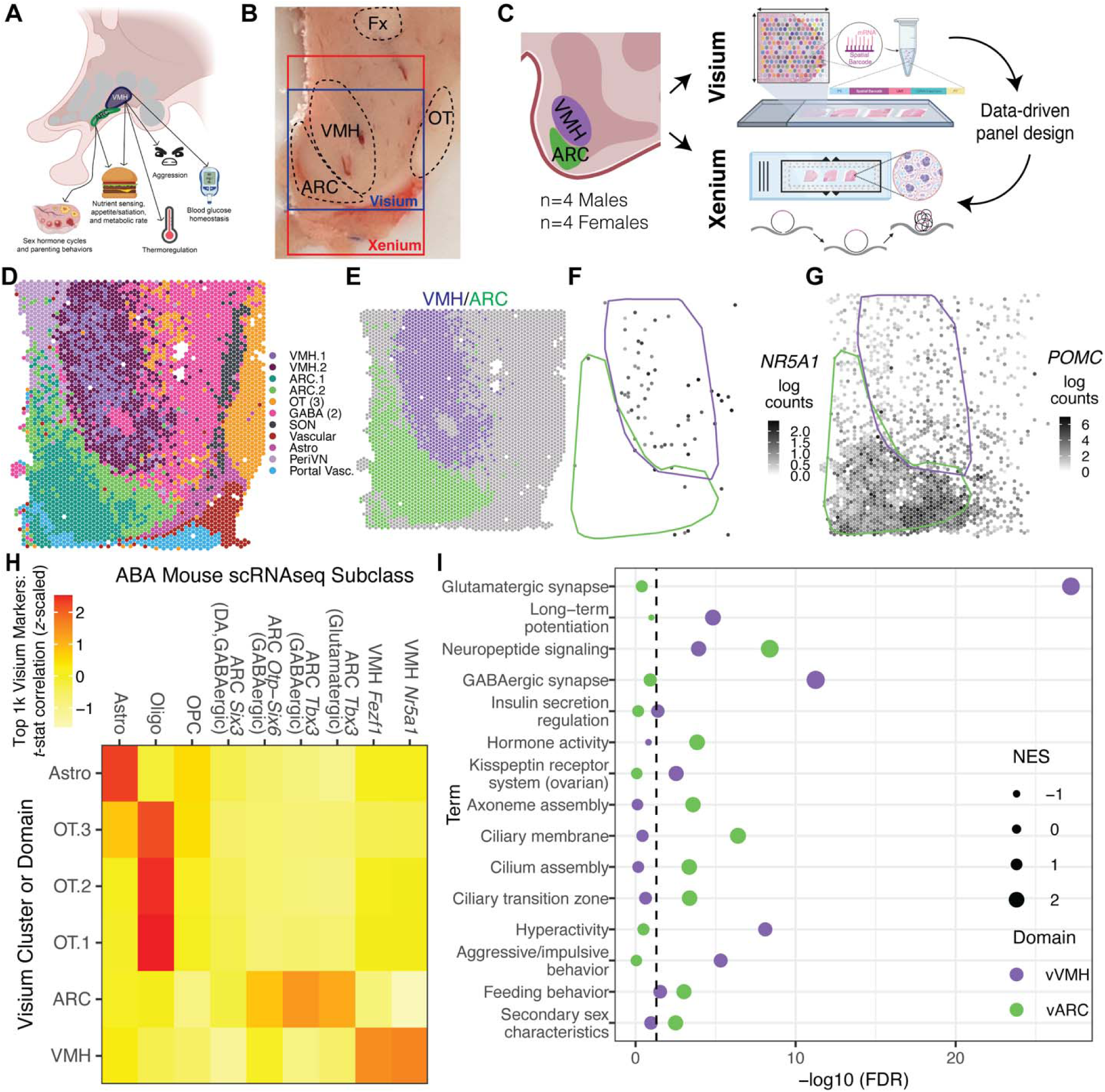
Study overview and identification of VMH and ARC domains. (**A**) Sagittal view of the human HYP highlighting the arcuate (ARC) and ventromedial hypothalamus (VMH) and main functions. (**B**) Tissue block (Br1225) sectioned for Visium and Xenium, indicating the ARC, VMH, and landmark white matter tracts (fornix [Fx] and optic tract [OT]). Boxes denote sampling areas for Visum (blue) and Xenium (red), which have different sized capture areas. (**C**) Study design overview. We first utilized the Visium spatial gene expression platform which uses ∼5,000 expression spots with unique spatial barcodes that correspond to an associated H&E image for a single capture area. We then performed *in situ* sequencing on the Xenium platform to acquire cellular resolution data for 266 pre-defined ‘brain panel’ genes and 100 custom genes based on domain marker, spatial, and sex-differential patterns from Visium. (**D**) *BayesSpace* clustering of Visium data shown for v1225A_M at *k*=15, with 3 optic tract/white matter clusters grouped together and 2 GABAergic neuron clusters grouped together. (**E-G**) Same sample as D, illustrating the Visium (v)VMH and vARC domains, which respectively express known markers *NR5A1* (**F**) and *POMC* (**G**). Approximate boundaries of vVMH and vARC domains labeled in same colors as **E**. (**H**) Correlation between marker statistics for vVMH/vARC and mouse cell subclasses assigned to these domains in a recent whole-brain mouse atlasing dataset (*85*). (**I**) GSEA analysis of vVMH- and vARC-enriched genes for ontology/function terms aggregated in MSigDB. NES: normalized enrichment score (a GSEA output metric). Sample identifiers are given as platform (v or x), donor number (4 digits), replicate tissue section (A-D), followed by an underscore and donor sex.

The rodent VMH has molecularly distinct anatomic subdivisions (*3*, *12*) linked to regulation of metabolism, appetite, and parenting behavior. VMH modulates both adrenal (*13*) and peripheral nervous catecholamine release (*14*, *15*), and contains a neuronal population expressing corticotropin-releasing hormone receptor 2 (*Crhr2*) (*16*), suggesting critical roles in regulating the autonomic nervous system and stress-related behaviors (*17*, *18*). VMH roles in metabolism (*19*) include glucose-mediated modulation of brain-derived neurotrophic factor (BDNF) expression (*7*), potentially in a sex-differential fashion (*20*). VMH is necessary for several sex-differentiated mouse behaviors, including mating, maternal aggression, and male fighting (*1*, *21–23*). Both of these functions are partially mediated by ventral lateral subdivision (VMHvl) neurons coexpressing estrogen receptors (*Esr1*) and progesterone receptors (*Pgr*) (*3*, *21*, *22*) and sexually differentiated during developmental androgen exposure (*10*).

The ARC is adjacent to the VMH and serves two primary functions, the first of which is metabolic regulation. ARC neurons detect circulating nutrients and nutritional status hormones (e.g., leptin)—in part via primary cilia (*24*)—and release neuropeptides including ghrelin, growth hormone releasing hormone (GHRH) (*25*), neuropeptide Y (NPY), and agouti-related peptide (AgRP) (*26*) throughout the HYP and pituitary to modulate metabolism and behavior. Second, *Kiss1* (kisspeptin)-*Esr1* coexpressing ARC neurons regulate cyclic release of gonadotropin releasing hormone (GnRH) from HYP, resulting in pulsatile pituitary release of luteinizing hormone (LH) and follicle-stimulating hormone (FSH) (*9*, *27*, *28*)*. Kiss1*- and metabolism-regulating populations of ARC bidirectionally influence one another to alter metabolism/growth according to reproductive status in a sex-differentiated manner (*6*, *29*).

While the ARC and VMH have been extensively studied in rodents, little is known about their function in the human brain despite their fundamental importance to survival and their potential roles in neuropsychiatric disorders. At the molecular scale, *in vitro* studies of human ARC-like neurons have identified enrichment in genomic regulators associated with major depressive disorder (MDD) or bipolar disorder (*30*). Furthermore, diagnostic features of several neuropsychiatric disorders implicate pathways in which VMH/ARC participate: examples include eating disorders, which by definition involve appetitive/metabolic (dys)regulation; major depressive disorder, which can involve increased or decreased appetite; and anxiety/posttraumatic stress disorders, which may stem from deficits in fear extinction (in part a VMH-regulated process (*31*)). Notably, all of these disorders have a sex-differential prevalence (*32–36*). Likewise, sex differences in molecular and behavioral phenotypes relevant to mental health are documented in many rodent HYP studies (*1*, *3*, *4*, *21*, *22*, *31*, *37*). In sum, observations ranging from animal models to clinical phenotypes suggest that the HYP may mediate sex differences in molecular risk for—and presenting features of—neuropsychiatric/neurodevelopmental disorders, highlighting the importance of investigating molecular sex differences in human VMH and ARC.

In-depth molecular profiling of individual subregions of the human HYP has been limited. Previous analyses of bulk RNA-sequencing (RNA-seq) data from adult human tissue by UKBEC (*38*) and GTEX (*39*, *40*) revealed substantial sex differences in HYP, though these could not be attributed to specific regions (*41*, *42*). To date, single-cell (*43*) and spatial transcriptomic (*44*) atlases from adult human HYP tissue have included only male samples or were underpowered for sex-differential analyses, respectively. To our knowledge, transcriptional sex differences within molecularly-defined HYP cell types have been the focus of only one report, with limited information on tissue anatomy and sample handling (*45*). Thus, significant questions remain regarding the regional enrichment and cell type-specificity of molecular sex differences in the human VMH and ARC.

Here, we utilized the 10x Genomics Visium and Xenium platforms to profile spatial gene expression in human postmortem VMH and ARC from 8 adult control donors without psychiatric diagnoses, evenly split by sex. A preliminary analysis of transcriptome-wide Visium data was used to prioritize genes for follow up at single cell resolution with a custom gene panel using Xenium. Using Visium, we illustrate that transcriptome-wide sex-differential expression (sex-DE) patterns in VMH and ARC are well-correlated. Subsequent single cell resolution measurements with Xenium confirmed regional sex DE observations, and pinpointed cell types responsible for observed sex-DE in each region. Further, we identified potential transcription factors (TFs), including gonadal hormone- and retinoid-binding TFs and TFs associated with neuropsychiatric disorders, whose putative regulatory targets are enriched for sex-DE genes. Across cell types residing in VMH or ARC, we identify 122 genes that are both sex-DE and linked to neuropsychiatric disorders with sex differences in prevalence or clinical features. Finally, we provide evidence that VMH and ARC cell types are enriched for sex-DE genes associated with disorders that show sex differences in prevalence rates, including autism, depression, and schizophrenia. We have made this state-of-the-art multisample, multimodal spatial atlas of molecular features and sex differences in the adult human VMH and ARC via interactive web resources at https://research.libd.org/spatial_HYP.

## Materials and Methods

### 3.1 | Experimental design: postmortem human tissue samples

Postmortem human brain tissue from 8 adult neurotypical donors of European ancestry was obtained at the time of autopsy following informed consent from legal next of kin, Office of the Chief Medical Examiner of the State of Maryland, under the Maryland Department of Health IRB protocol #12-24, from the Western Michigan University Homer Stryker MD School of Medicine, Department of Pathology under the WCG protocol #20111080, and from the NIMH Human Brain Collection Core, protocol #90-M-0142. Using a standardized strategy, all donors were subjected to clinical characterization and diagnosis. Macro- and microscopic neuropathological examinations were performed, and subjects with evidence of neuritic pathology or other neuropathological abnormalities were excluded. Details regarding tissue acquisition, processing, dissection, clinical characterization, diagnosis, neuropathological examination, RNA extraction and quality control (QC) measures have been previously published (*46*). Demographics for the 8 neurotypical control donors included in the study are listed in **Table S1**. All subjects had BMIs between normal range and class I obesity (22.3-31.6; 7 of 8 donors non-obese with BMI < 29) and no history of neuroendocrine disorders. Female subjects were premenopausal and not pregnant at time of death. Information on donor use of oral contraceptives was not systematically collected. Briefly, fresh frozen coronal brain slabs with clearly visible thalamus, hypothalamus, putamen, globus pallidus (internal and external segments), optic tract, anterior commissure, fornix, and posterior amygdala (MNI: -7.04) were selected for VMH and ARC dissection. Using a hand-held dental drill, tissue blocks of approximately 10 X 20 mm were dissected, encompassing the hypothalamus, optic tract, anterior commissure and fornix. White matter (WM) tracts were used as landmarks to identify the approximate location of VMH and ARC for microdissection. Tissue blocks were stored in sealed cryogenic bags at -80°C until cryosectioning.

### 3.2 | Tissue processing and quality control

At the time of cryosectioning, tissue blocks were acclimated to the cryostat (Leica CM3050s) at - 14°C and mounted onto a chuck with OCT (TissueTek Sakura). Approximately 50 µm of tissue was trimmed from the block to achieve a flat surface, and several 10 µm sections were mounted onto pre-chilled glass slides for anatomical validation quality control (QC) experiments using single molecule fluorescent in situ hybridization (smFISH) with RNAScope technology (Advanced Cell Diagnostics [ACD]). RNAscope was performed as described below to verify the presence of neuroanatomical landmarks with probes specific for the VMH (Hs-NR5A1-C1, catalog no. 553151), paraventricular nucleus/supraoptic nucleus ([PVN/SON]; Hs-OXT-C2, catalog no. 538811), ARC (Hs-NPY-C3, catalog no. 416671), and WM tracts (Hs-MBP-C4, catalog no. 411051). Successive rounds of RNAscope were completed until the anatomical plane containing VMH and ARC was reached. This location approximately corresponds to the area between the optic tract (as the ventro-lateral landmark) and the fornix (as the dorsal landmark). Blocks were then scored with a razor blade to isolate the VMH and ARC nuclei in ∼6.5 X 6.5 mm squares for the Visium assay. Adjacent 10 µm tissue sections were mounted onto pre-chilled Visium Spatial Gene Expression slides (part number 2000233, 10x Genomics) (**Fig. S1**) and onto Superfrost plus glass slides, which were banked for additional RNAscope validation experiments. In some cases, tissue sections from the same block were collected onto multiple Visium arrays to ensure inclusion of target structures. In these cases, distance along the anterior-posterior axis between experiments did not exceed 100μm. Following Visium analysis, tissue blocks were re-mounted for cryosectioning, approximately 50 µm of tissue was trimmed off to achieve a flat surface, and several 10 µm sections were mounted onto pre-chilled glass slides and banked for validation studies. Blocks were then re-scored with a razor blade to isolate the VMH and ARC nuclei in ∼12 X 6.5 mm strips and mounted onto Xenium slides (Part Number 1000460, 10x Genomics). 3-4 tissue strips were fitted onto each Xenium array.

### 3.3 | Visium data generation

Visium Spatial Gene Expression slides were processed according to the manufacturer’s protocol (protocol number CG000239, Rev G, 10X Genomics) as previously described (*47*, *48*). The tissue permeabilization step was performed for 18 minutes based on tissue optimization experiments in other brain regions (*48*, *49*). For each Visium slide, H&E staining was performed and the slide was coverslipped. Tissue sectioning and mounting to slide, sequencing library preparation, sequencing, H&E staining, and microscopy were performed across two sites (listed for each step per sample in **Table S1**). Briefly, microscopy images were acquired either at 40x on a Leica CS2 slide scanner equipped with a 20x/0.75NA objective and a 2x doubler, or on an Olympus VS200 slide scanner equipped with a 10x/0.4NA objective. Following removal of the coverslip, tissue was permeabilized, cDNA synthesized, and sequencing libraries generated. Libraries were loaded at 300 pM on a NovaSeq6000 System (Illumina) at the sites listed in **Table S1**. Sequencing was performed according to manufacturer’s instructions, targeting ∼60K reads per spot.

### 3.4 | Xenium Data Generation

Xenium slides were fixed and permeabilized according to 10X Genomics’ *Xenium In Situ for Fresh Frozen Tissues – Fixation & Permeabilization Protocol* (CG000581, Rev C). Briefly, slides were warmed for 1 minute prior to submersion in 3.7% formaldehyde for 30 minutes. Slides were then permeabilized with 1% sodium dodecyl sulfate solution (SDS) for 2 minutes and rinsed with PBS. This was followed by a 1 hour incubation in 70% methanol on ice. Following PBS washes, slides were placed in their respective cassettes and incubated with 0.05% PBS-T. For probe hybridization, ligation, and amplification, the 10X Genomics protocol: Xenium In Situ Gene Expression was followed (CG000582, Rev E). The probe hybridization solution was then prepared. We used a combination of the off-the shelf “human brain,” probe panel designed by 10X Genomics (Xenium Human Brain Gene Expression Panel, Part No.1000599) and our custom probe panel (10X Genomics; Xenium Custom Gene Expression Panel, Part No. 1000561). Details of the custom probe panel design are described in section 3.8.1. Probe mix was applied to each slide, a cassette lid was applied to each slide cassette and the slides were incubated at 50°C overnight.

The next day, the slides were washed twice in 1X PBS-T followed by an incubated wash in post hybridization wash buffer (Xenium Post Hybridization Wash Buffer, Part No. 2000395, 10X Genomics). Three washes of 1X PBS-T were completed before addition of ligation mix to each slide.Slides were then incubated at 37°C for 2 hours. After ligation, the slides were washed three times in 1X PBS-T, and then incubated in an amplification master mix at 30°C for 2 hours. After amplification, slides were washed three times in TE buffer followed by three washes in 1X PBS and then incubated in diluted reducing reagent B for 10 minutes at room temperature. Slides were then washed in 70% ethanol followed by two washes of 100% ethanol. Autofluorescence solution was then added to each slide and incubated in the dark for 10 minutes at room temperature. Slides were then rinsed three times with 100% ethanol before drying on a 37°C preheated thermocycler for 5 minutes. Additional washes were performed before adding Xenium Nuclei Staining Buffer, which incubated for 1 minute in the dark at room temperature. Slides were then washed three times in 1X PBS-T and stored at 4°C prior to being loaded on the Xenium analyzer at the Johns Hopkins Single Cell & Transcriptomics Core (SCTC). The solutions needed to run the Xenium analyzer were prepared according to the manufacturer’s instructions. Region selection was guided by our team with instrument operation performed by the SCTC Core.

Upon completion of the Xenium analyzer run, post-xenium H&E was completed following 10X Genomics Xenium In Situ Gene Expression - Post-Xenium Analyzer H&E Staining protocol (CG000613, Rev A) with the following modifications: reduced staining time in hematoxylin, from 20 minutes to 17 minutes, and reduced eosin staining time, from 5 minutes to 2 minutes. Briefly, slides were quenched in a 10mM sodium hydrosulfite solution for 10 minutes, slides were then washed four times in milli-Q water before being stained with Mayer’s Hematoxylin for 17 minutes (Sodium hydrosulfite, Cat No. 157953-5G, Sigma Aldrich; Hematoxylin Solution, Mayer’s, Cat No. MHS16, Sigma Aldrich). After nuclear staining was completed, slides were rinsed three times in milli-Q water before submerging into bluing buffer (Bluing Solution, Cat No. CS702, Dako). Slides were washed once in milli-Q water, once in 70% ethanol, and once in 95% ethanol before being introduced to alcoholic eosin (Eosin Y Solution, Alcoholic, Cat No. 3801615, Leica). After cytoplasmic staining was completed, the slides were rinsed twice in 95% ethanol and twice in 100% ethanol before being cleared in two changes of xylene (Xylene, Reagent Grade, Cat. No. 214736, Millipore Sigma). Slides were then coverslipped using Vectamount mounting media and allowed to dry overnight before imaging (VectaMount Permanent Mounting Medium, Cat No. H-5000, Vector Laboratories Inc.). Imaging was completed at 20X and 40X using a Leica Aperio CS2 digital pathology slide scanner. While H&E images were not utilized in the current study, they have been made available with other supplemental data through a Globus endpoint (see *Data Availability*).

### 3.5 | Single molecule fluorescence in situ hybridization (smFISH)

smFISH assays were performed with RNAscope technology utilizing the RNAscope^TM^ Multiplex Fluorescent Reagent Kit v2 (catalog no. 323100; Advanced Cell Diagnostics [ACD]) in reference to the manufacturer’s protocol (document no. UM323100, rev B; ACD) with the following RNAscope probe sets: Anatomical quality control: Hs-NR5A1 (assigned Opal dye 570), Hs-OXT (assigned Opal dye 620), Hs-NPY (assigned Opal dye 690), Hs-MBP (assigned Opal dye 520) (catalog no. 553151, 538811-C2, 416671-C3, 411051-C4, respectively; ACD); Experiment 1: Hs-NR5A1 (assigned Opal dye 570), Hs-LAMP5 (assigned Opal dye 620), Hs-GAD1-01 *or* Hs-GAD2-ver2 (assigned Opal dye 690), Hs-SLC17A6 (assigned Opal dye 520) (catalog no. 553151, 487691-C2, 573061-C3 or 415691-C3, 415671-C4, respectively; ACD); Experiment 2: Hs-KISS1 (assigned Opal dye 520), Hs-ESR1 (assigned Opal dye 570), Hs-TAC3 (assigned Opal dye 690) (catalog no. 507981, 310301-C2, 507301-C3, respectively; ACD). Specifically, 10-µm tissue sections were fixed in 10% neutral buffered formalin for 30 minutes at room temperature (RT). After 2 PBS washes, the samples were sequentially dehydrated in 50%, 70%, 100%, and 100% ethanol solutions each for 5 minutes at RT, treated with hydrogen peroxide for 10 minutes at RT, washed 2 times again with PBS, and treated with protease IV for 30 minutes at RT. After another 2 PBS washes, sections were incubated in a solution of the four target probes for 2 hours at 40°C, washed 2 times in wash buffer (catalog no. 310091; ACD), and stored in SSC for up to 48 hours at 4°C. Afterward, the sections underwent amplification with AMP1 through AMP3 for 30, 30, and 15 minutes, respectively, at 40°C with 2 washes with wash buffer performed before each new reagent was added. In sequence, the sections underwent 4 cycles of 15-minute incubation with HRP-Cx, 30-minute incubation with a fluorescent Opal dye solution, and 15-minute incubation with HRP blocker, all at 40°C and with 2 washes with wash buffer in between. Each cycle used a different fluorescent Opal dye (520, 570, 620, or 690; catalog no. FP1487001KT, FP1488001KT, FP1495001KT, and FP1497001KT, respectively; Akoya Biosciences) diluted 1:500 in TSA buffer. Finally, the tissue sections were treated with DAPI for 20 seconds at RT, decanted, and then coverslipped with Fluoromount G mounting medium (catalog no. 0100-01; Southern Biotechnology). Coverslipped slides were stored in the dark for at least 24 hours before imaging. Slides were imaged as z-stacks (at least 10 per image) using a Nikon AXR confocal microscope system powered by the NIS-Elements imaging software with a Nikon APO lambda D 20x / 0.80 objective and/or Nikon APO lambda S 40x / 1.25 objective. For anatomical quality control, slides were imaged in a single plane with a Nikon Plan APO lambda D 2x / 0.1 objective. All confocal images at the resolutions presented/analyzed are available with other supplemental data files through a Globus endpoint (see *Data Availability*).

#### 3.5.1 | Puncta & Colocalization Quantification

For Experiment 2 HALO (Indica Labs) was used to segment and quantify fluorescent signals for each probe in single cells. Nikon .nd2 files were imported into HALO and viewed at the manufacturer’s recommended magnification (40x) for punctate probe signal quantification. The FISH-IF module was used to quantify RNA transcripts (copy counts) within each detected object (i.e. a nucleus with radially defined “membrane” at 3 um to estimate a cell) in accordance with the manufacturer’s guidelines: HALO 3.3 FISH-IF Step-by-Step guide (Indica labs, v2.1.4 July 2021). For each image, fluorescent detection parameters were optimized for DAPI and probe signal to ensure image faithfulness in accordance with the HALO 3.6 User Guide (Indica labs, February 2023). To ensure accurate counting of RNA transcript copies assigned to each cell for each of the 3 RNAScope probe channels, the median probe signal intensity value of positive cells per signal was input into HALO as the representative intensity value parameter for that channel’s copy intensity parameter consistent with previously published work (*50*).

HALO .csv outputs were imported into R. As previously described (*51*), *k*-means clustering was performed on HALO variables Signal Area and Copy Count with a k = 3 to cluster “high,” “medium,” and “low/no” expressing cells) for each of the 3 RNAScope probe channels. High (signal area: *KISS1*–41-85um^2^; *ESR1*–17-62um^2^; *TAC3*–56-127um^2^; and copy count: KISS1–27-82; *ESR1*–6-33; *TAC3*–12-68) and medium (signal area: *KISS1*–13-48um^2^; *ESR1*–4-19um^2^; *TAC3*–20-61um^2^; and copy count: *KISS1*–2-29; *ESR1*–1-12; *TAC3*–3-32) clusters were deemed positive. All cells were plotted as circles with x and y coordinates as determined by HALO, and colored by positivity for each channel. The ARC region of interest (ROI) was selected by drawing a polygon around the area expressing *TAC3* and *KISS1* for each donor. A data frame was then produced containing all information for each segmented cell in the ROI, including cell phenotypes (channel(s) meeting the positivity definition above). Phenotype occurrences were then tabulated to quantify cell phenotype counts and proportions. All HALO settings files are provided on GitHub.

### 3.6 | Visium H&E image segmentation and raw data processing

Multi-sample slide images acquired on the Leica CS2 were first processed using VistoSeg software (*52*), which divides the whole-slide Visium images into individual capture area images using the *splitSlide* function. The Olympus images were of single capture areas and did not require such processing. Images for each capture area were aligned in Loupe Browser (10x Genomics) with the gene expression spots captured. Slides were then processed using the Spaceranger v. 2.1.0 software from 10X Genomics, yielding feature counts for each spatial location (“spot”) per sample using the .json output from Loupe browser, the sample image, and FASTQ sequencing files.

### 3.7 Visium Data Analysis

All analyses were performed using R 4.4.1 unless otherwise noted. Analysis was performed within the SpatialExperiment R framework (53), coercing outside datasets in other formats to this framework where necessary using zellkonverter (54). Spatial expression and clustering plots were generated using SpatialLIBD (55) or escheR (56). ggplot2 was used to generate heatmaps and scatter/volcano plots unless noted. Extended color palettes were generated using the package Polychrome (57).

#### 3.7.1 | QC and Normalization

Fraction of mitochondrial reads and number of unique genes identified were calculated for each spot. Plots of these values were manually examined for spatial patterning in each sample to ensure selection of filter thresholds that would not remove putative biological information (for example, WM tracts would be expected to have low gene diversity in a contiguous area, a matter of biological rather than technical variance). Using *SpotSweeper* (*58*), we first detected and removed high-end outliers relative to surrounding spots in terms of mitochondrial read percentage, low-end outliers for number of UMIs, or low-end outliers for number of unique genes. Subsequently, we filtered remaining spots for high mitochondrial percentage, low UMI count, and low unique gene count. We noted during exploratory analyses that the highest mitochondrial read fractions were largely in spots later identified as VMH—containing the only excitatory neurons of note in the samples—and spots along its lateral edge. We thus set a lenient threshold of 50% for mitochondrial read content in the final analysis. We plotted histograms of UMI and gene count distributions across all remaining spots and identified a clear drop-off in spot frequencies within the histograms at approximately ≤210 UMIs and ≤126 unique genes, suggesting the boundary between technical artifact and biologically low content. Indeed, such spots were almost exclusively localized to sample edges, and these thresholds were therefore used to remove low-content spots. For the retained spots, library sizes (number of reads) per spot were calculated, normalized, and log2 transformed (*59*) across all spots in all samples to get expression values per gene per spot.

Of 4,992 Visium spots possible in a sample, tissue occupied 4,621±341 (mean ± standard deviation) spots. Spots with high mitochondrial read proportion (≥50%) were removed to preserve a potentially biological pattern of high mitochondrial counts adjacent to VMH (**Fig. S2C**). Subsequent filtering for low unique molecular identifiers (UMIs) and/or gene diversity (≤210 UMIs, ≤126 unique genes) resulted in 4,508±363 spots for downstream analysis. The filtered data included 6,059±5057 UMIs/spot (mean±SD) spanning 2,536±1470 unique genes/spot, covering 45,074 spots and 30,361 unique genes altogether. Additional pre- and post-filtering metrics are included in **Table S1**.

#### 3.7.2 | Feature Selection

We used the filtered data to identify highly-variable genes (HVGs, spatially-naive) and spatially variable genes (SVGs) using the packages *scran* (*60*) and *nnSVG* (*61*), respectively. For HVGs, we used sample as the blocking factor and collected the top 10%ile and 20%ile of resultant genes using *scran::getTopHVGs(…, prop=0.1 or 0.2)*(*60*) (**Table S2**).

In the nnSVG approach, each sample is analyzed independently and genes are pre-filtered. Here, we filtered to genes with at least 3 counts in at least 0.5% of spots in a sample. We subsequently recalculated the logcounts for use within nnSVG to account for dropped genes. We extracted and concatenated the nominally significant SVGs from each sample (**Table S2**) and determined the experiment-wide SVGs by averaging their ranks across samples where the gene achieved nominal significance. We then filtered the union of samplewise SVGs to those that achieved nominal significance in three or more samples. These features were sorted in ascending order of mean rank (among samples where the feature achieved nominal significance for spatial variation) to collect the same number of features as contained in 10%ile and 20%ile HVG sets.

#### 3.7.3 | Selecting the Number of Clusters

In a preliminary subset of samples, we used each of the four feature sets described above for performing dimensionality reduction concurrent with batch correction using *Harmony* with sample ID as the grouping variable (*62*). The default output of 50 reduced dimensions were appended to the *SpatialExperiment* object for downstream use separately for each feature set.

To determine an expected number of spatial domains, we turned to the Allen Brain Atlas for adult human (*63*) and tabulated the number of unique anatomic areas denoted along the entire anterior-posterior span of the VMH, yielding 14 potential anatomically-defined domains. We assumed WM would comprise a single, homogeneous domain and added this to get a grand total of 15 potential anatomic areas: dorsal, central, and ventral VMH; ARC; periventricular nucleus, tuberal part; lateral hypothalamic area, tuberal part; posterior hypothalamic nucleus; dorsomedial nucleus; tuberomamillary nucleus; lateral tuberal nuclei; paraventricular nucleus; magnocellular nucleus of the lateral hypothalamic area; supraoptic nucleus; accessory secretory cells of lateral hypothalamus; and generic WM.

In this same preliminary subset of data, we also performed spatially-agnostic, weighted nearest-neighbors clustering using *buildSNNgraph* from *scran*. We utilized *Harmony* batch-corrected principal components calculated on the 10%ile or 20%ile highly variable gene sets, and constructed two SNN graphs for each feature set, using *k*=5 or *k*=10 nearest neighbors, for a total of four initial neighbor graph spaces. From each of these four neighbor graphs, we divided the tree into 4, 5…47 clusters using *igraph*::*cut_at* and aggregated these assignments. We determined the optimal, spatially-agnostic clustering resolutions for maximal within-cluster concordance relative to between-cluster concordance using the *fasthplus* method (*48*, *64*). The values returned as *seg_psi_est*, *seg2_psi1_est*, and *seg2_psi2_est* represent the three estimated clustering values most optimal for a given graph. These 12 estimates included two values rounding to 9, three in the range 14.6-15.6, three ranging 17.0-19.8, and a maximum value of 30.8 leading us to select the recurrent minimum (9), the value consistent with anatomic expectations (15), the upper end of this range (20), and the highest value (31) as the number of clusters to generate in the final data.

We then utilized *BayesSpace* (*65*) with parameters of *k*=15 and 20,000 iterations for spatially-aware clustering of the *Harmony* reduced dimensions for each of the four variable-feature sets. The resulting domain assignments from each feature set were comparable when plotted onto the spatial data and plots of the top principal component; however, UMAP plots labeled by domain assignment revealed that the majority of domains separated into multiple sectors of UMAP space, except when using the “10 %ile” nnSVG set. We therefore used this feature set and its derived spatial domains for all subsequent analysis.

#### 3.7.4 | Dimensionality Reduction and Clustering

For the analyses of the full dataset presented here, we only utilized an equivalently-derived “10 %ile” nnSVG feature set for dimensionality reduction by principal components analysis (PCA), UMAP, and sample-level batch correction using *Harmony* (**Data S1**) (see *Code Availability*) with initialization of the lambda parameter—which alters the strength of correction—from the data itself, rather than requiring a user-defined initial value (an approach which we term “lambda null” or “lambda NA” throughout the methods and in certain Supplemental files). We used the Harmony batch correction as input dimensions for clustering in *BayesSpace* at resolutions of *k*=15, 20, and 31. All results presented reflect *Harmony BayesSpace* clusters computed with *k*=15 and 60,000 iterations. We produced additional clustering assignments with the nnSVG “10th %ile” feature set using reduced dimensions from MNN batch correction (*66*) or *Harmony* with its default lambda value of 1 as inputs to 60,000-iteration BayesSpace runs at *k*=15, 20, and 31 (see *Code Availability*). Marker, sex-DE, and GSEA results are provided for clusters defined after using the development version of *Harmony* (“lambda null”-enabled) and BayesSpace *k*=15 (individual clusters), *k*=15 (VMH/ARC clusters joined into single clusters), *k*=20 and *k*=31 (see *Data Availability* and *Supplemental Materials*). Since spots can contain multiple cells, we emphasize that we subsequently call these clusters “domains’’ to emphasize that they do *not* necessarily correspond to single cell types or subtypes as would be identified in single-cell or - nucleus approaches.

#### 3.7.5 | Marker Gene Identification

Marker genes were comprehensively determined using several complementary statistical approaches. We primarily utilized the *registration_wrapper* function in *spatialLIBD* (*55*) to calculate *t*-test statistics comparing log counts in each domain vs all others (this method being most analogous to the logFC comparisons assessed using Cohen’s *d* in *scran*). We performed select analyses using the *scoreMarkers* function from *scran* (*60*) to identify marker genes for select pairs of domains (e.g., to compare VMH subdomains) using multiple statistics: a Cohen’s *d* statistic for mean log fold-change (logFC) between compared groups; an AUC statistic, indexing the probability that a gene in a random spot drawn from domain/group 1 is greater than in a random spot drawn from domain/group 2; and the logFCs calculated when only considering spots from each domain/group where a given gene was detected. We focus on *SpatialLIBD t*-test statistics in the main text for both global marker identification for comparisons between VMH or ARC subdomains. Putative VMH/ARC subdomains from the additional clustering approaches were identified based on marker genes and referred to throughout the supplement as (*VMHorARC)(k).(digit)*, where each subdomain is assigned a sequential number arbitrarily (e.g., “VMH20.1” and “VMH20.2” are two subdomains of VMH identified from *k*=20 BayesSpace analysis). To discern genes differentiating among VMH or ARC clusters from Visium, we additionally performed ‘one-vs-all-other’ analyses, wherein each cluster from a domain was separately compared to other clusters *only* from that domain by subsetting the data only to spots with VMH or ARC cluster labels before performing marker analysis using *spatialLIBD* as above. For *k*=15 or 20, this amounted to a single pairwise comparison for each domain, as VMH and ARC each spanned two clusters at both of these clustering resolutions.

#### 3.7.6 | Spatially Variable Gene Identification within VMH and ARC

To further explore whether we could identify spatial patterns of gene expression within the VMH and ARC specifically, we subset the data to those spots assigned a VMH or ARC domain as above and re-ran nnSVG on individual samples, only considering spots within ARC or within VMH. One sample was excluded from this analysis for VMH due to an impractically small number of VMH spots (sample v5459A_M, 330 spots) for SVG analysis. One sample was likewise excluded from ARC analysis for the same reason (sample v8667A_F, 207 spots). The threshold for gene inclusion for SVG analysis in a given sample was ≥1 count in at least 5% of spots within the domain being analyzed. The ranks of nominally significant genes per sample were tabulated and aggregated to their means across those samples where the gene achieved nominal significance, as was done for the feature selection steps described above. As the collective structure of VMH and ARC domains was similar at all clustering resolutions, we only performed these SVG analyses on the domains as defined by the *k*=15 BayesSpace results.

#### 3.7.7 | Pseudobulk Sex-Differential Expression Analyses

We set out to test each Visium cluster/domain for sex differences in expression. To achieve this, spots with a given cluster/domain level were pseudobulk aggregated within-sample using *dreamlet* 1.3.1. We then subsetted to clusters found in at least 5 samples (at least 2 per sex). Pseudobulk counts were preliminarily normalized to total counts using TMM normalization. We then programmatically set a low-expression filter by detecting the smaller sex group (in terms of sample number) among the retained samples for a given cluster, detecting the second minimum in a histogram of gene-wise mean log counts. This value was used to perform low-expression filtering such that retained genes which adhered to an approximately normal distribution. TMM factors were then recalculated for the remaining genes, followed by DE analysis using *voomLmFit*, with donor as a blocking factor (which is internally passed to *DuplicateCorrelation*), *adaptiveSpan* set to true to identify the number genes to use for smoothing the mean-variance trend, *sample.weights* set to true to perform an analysis analogous to the *voomLmFit* predecessor *voomWithQualityWeights*, and fitting a model of *∼0+Sex* to test for sex effects. This was run for all batch correction and clustering resolutions described above, with results available in the Supplement.

#### 3.7.8 | Gene Set Enrichment Testing of Markers and Sex-DEGs in GSEA-MSigDB and for Transcription Factor Target Genes

We performed comprehensive, threshold-free analyses of genes marking spatial domains to confirm expected functional roles and identify potentially unexpected ones. Using the *t*-statistics of all genes tested in marker analysis from *spatialLIBD* (*55*), we applied Gene Set Enrichment Analysis (*67*, *68*) as implemented in the package *fGSEA* (*69*) to analyze marker enrichment in gene sets from MSigDB (*67*, *70*, *71*). We included “hallmark” sets, along with sets representing chromosomal regions; curated sets from publications, including pathway databases like KEGG; encompassing predicted microRNA (miRNA) / transcription factor (TF) targets; defined in ontologies; and reported human cell type markers. (These correspond to genesets “h.all”, “c1.all”, “c2.all”, “c3.all”, “c5.all”, and “c8.all”, respectively; all versioned “2022.1”.) We also included the MSigDB “oncogenic signature”/”c6.all” dataset as a null comparator and to screen for transcriptomic changes suggestive of HPA tumors, which are relatively common and whose presence would run counter to our present goal of profiling healthy neurotypical hypothalamus.

We additionally aggregated several databases of tissue-*non*specific putative target genes of transcription factors (TFs) ascertained from model organism and human cell/tissue types. We collated TF-target sets from ChEA 2022 (*72*), TRRUST (*73*), those mined from publication supplements by Rummagene (*74*), and those curated in the Enrichr (*75*) tool (Gene Expression Omnibus-mined results of TF perturbation and knockout experiments and TF protein-protein interactions). We subsequently filtered the results for each domain to those TFs expressed in that domain. Given that TFs are often lowly expressed at the mRNA level (*76*), we used lenient filters to define TFs as expressed: for marker gene enrichment, we required the at least 1% of spots to have log counts >0 for the TF in ≥5 (majority) samples and a mean [% of sample’s domain spots with nonzero counts of the TF] >5%. For sex-DE enrichment testing, we considered a TF expressed if ≥3 (smallest group size, when considering limited ARC or VMH spots in a Visium sample from each sex—see *Methods 3.7.6*) samples had at least 0.25% of spots in a domain with log counts >0 and a dataset mean of at least 1% of spots in the cluster with log counts >0. Our rationale for this difference in parameters was that our marker-based statistics were for broad domains, such that regulatory signatures defining the domain should be relatively abundant, whereas for sex differences, the expectation is not necessarily that all (below-detectable resolution) subdomains or cell types/subtypes in the domain—ie., a much smaller number of spots than define the domain itself—drive the detected DE events.

Gene sets analyzed were limited to those containing 15-500 genes for MSigDb and TF-target analyses. We utilized signed, transcriptome-wide one-vs-all marker or male-vs-female DE *t*-statistics as our ranked list for the respective analyses. To retrieve enrichment statistics from every gene set for both male-upregulated and female-upregulated expression patterns, we explicitly ran analyses for the top (male-upregulated / positive *t* statistic) and bottom (female-upregulated / negative *t* statistic) of the rank lists, as fGSEA testing on both directions simultaneously will otherwise return data for the effect direction with greater enrichment, even if both directions are significant. Additional sex-DE GSEA analyses were performed using the absolute value of DE *t*-statistics to identify gene sets enriched for sex-differential expression as a general phenomenon. To ensure the ability to resolve extremely small *p*-values, we set the fGSEA parameters *eps* to 0.0 to specify that p-values should be estimated for all enrichments, using 50,000 permutations for initial *p*-value estimations. To determine the most precise, independent enrichments for some of these analyses, we subsequently used the *collapsePathways* function in fGSEA. For marker gene analysis, we only examined “positive” enrichment, i.e., enrichments corresponding to genes more specific to a given spatial domain relative to others. For sex-DEGs, we used the same parameters and gene sets as above, but examined both directions of enrichment for each set of spatial domain’s DEGs (as the test statistic’s sign indicates male or female upregulation).

#### 3.7.9 | GWAS-Derived Gene Set Enrichment Testing

To comprehensively examine whether neuropsychiatric disorder risk genes were enriched among markers or sex-DE genes, we collected genes reported in common-variant (genome-wide association study (GWAS)-based) and rare-variant studies across several modalities. GWAS-derived gene sets comprised those reported in GWAS studies, explicitly *excluding* genes assigned to loci on the sole basis of being the most proximal to a phenotype-associated variant, while retaining e.g. genes reported in the GWAS publications resulting from “post-GWAS” analyses such as MAGMA, quantitative trait locus (QTL) colocalization, etc. For schizophrenia and autism spectrum disorders, we also collected genes identified in, e.g., whole exome sequencing cohorts of patients and healthy sibling probands. We additionally collected transcriptome-wide association study (TWAS) analyses performed on GWAS summary statistics for the same traits (generally from the same GWAS studies); largely collected from webTWAS (*77*),(*^78^*),(*^79^*),(*^80^*) and TWASatlas (*81*), as well as individual publications that performed TWAS. Finally, we gathered literature-mined, curated gene-disease pairs from DisGeNET (*82*). For major depressive disorder, these included several genes of historical interest that have been explicitly demonstrated *not* to play a role in heritable depression risk (*83*); we thus removed the genes covered by (*83*) from this set specifically. The sets from GWAS, TWAS, and DisGeNet and corresponding sources are provided in *Supplemental Materials*. For the analyses presented, we analyzed enrichments solely in the union of genes collected for each diagnosis across these three source types. As some of these gene sets fell outside the above bounds of 15-500 genes, the largest gene set size was elevated to 8200 to exceed the largest disease-gene list (BMI, union of all gene-associated modalities, >8100 genes).

#### 3.7.10 | Spatial Registration

We opted to map our Visium spatial domains to Allen Brain Atlas *in situ* hybridization (ISH) data from mouse (*84*) using *scCoco/cocoframer*, as ISH signal within atlas regions should reflect mixed neuronal and glial content akin to that measured with Visium. We provided the top 150 markers per Visium spatial domain based on one-vs-all logFC, and set a minimum of 50 genes with adult mouse ISH data for a given domain’s comparison to proceed. *scCoco* then was used to score the expression of query markers within each Allen Atlas Common Coordinate Framework (CCF) area of the hypothalamus. Scoring proceeds for one query marker set at a time (i.e., 50-150 genes) by ranking the ISH expression of each query gene in 10 voxels from every CCF hypothalamic area, then converting the median rank of each marker’s expression to a value ranging 0-1 by taking 1 - (median rank / number of voxels queried). Finally, these converted values are averaged across all genes for each CCF region to get an enrichment score ranging from 0 (least overlap) to 1 (most overlap). Results using development *Harmony* with *BayesSpace* at *k*=15, 20, and 31 are provided in the Supplement.

We additionally utilized a recent mouse brain atlasing effort that included single-cell RNA-sequencing results from 45 sequencing libraries collected across over 25 donor mouse hypothalami from both sexes (*85*). The *registration_wrapper command* in spatialLIBD to identify marker genes of each cell “subclass” (a relatively coarse level of hierarchical clusters reported in the Yao data). We then tested the correlation of marker *t* statistics from our spatial domains (k=15, 20, 31, and k=15 with collapsed VMH and ARC domains) with each mouse subclass’ *t*-scores for the same genes. We tested correlations using the top 25, 50, 75, 100, 250, 500, 750, and 1000 spatial domain marker genes (ranked by *t*-statistic), as well as using all genes with one-to-one mouse orthologs. We additionally repeated the analyses with all mouse VMH subclasses re-labeled as “VMH” and all ARC subclasses relabeled “ARC” to test domain correlation to collective VMH/ARC mouse types. We limited these analyses to a total of 47 subclasses—those represented by at least 15 cells per sample in 22 or more of the samples. Pseudobulk expression matrices from these analyses were also created and utilized for comparison of retinoid gene expression described below.

### 3.8 | Xenium Data Analysis

#### 3.8.1 | Gene Selection for Xenium Custom Panel

Genes were selected for measurement on the Xenium platform in an initial subset of 7 Visium samples, all from donors in the final dataset. (One sample from the preliminary dataset was not included in the final data due to low read depth compared to all other samples; a second, adjacent tissue section sampled from the same donor was processed using Visium to collect higher-depth sequencing data and is included in the final dataset). This data subset was filtered using traditional QC approaches, removing spots with over 35% mitochondrial reads, under 600 UMIs, and/or under 450 unique genes detected. This data was integrated across samples with Harmony, clustered using BayesSpace with k=15, and analyzed for marker genes, VMH/ARC-local SVGs, and sex-DE, as described above, with the difference that dreamlet 1.0.0 was used for pseudobulk sex-DE analysis and a random effect of donor. Top marker genes of one or both VMH or ARC domains were visually inspected in sample-wise fashion for consistent patterns of specific or enhanced expression in that domain across samples. For SVGs, nnSVG results were aggregated across samples by taking the mean of ordinal gene ranks in samples nominally significant (p<0.05) for spatial variability in at least 3 samples. The top 100 mean-ranked genes each for ARC and VMH were visualized in all samples to determine whether there was a consistent spatial pattern of expression within the ARC or VMH across donors, and visually robust genes were selected for inclusion. For sex-DE genes, the preliminary analysis with dreamlet 1.0.0 returned a much higher (compared to the final dataset) number of significant DEGs at FDR<0.05 due to algorithmic differences at that time, with approximately 150 in VMH and 400 in ARC. All genes meeting this significance threshold were plotted and visually examined in spotplots to determine whether the male/female pattern of expression difference was consistent. Further, visual inspection was used to rule out DE events that occurred due to genes predominantly detected at the edges and outside of VMH/ARC, which would indicate confounding due to differences in VMH/ARC boundaries relative to the neighboring domains primarily expressing the gene. Additional genes were selected on the basis of genes identified in model organism literature as marking VMH or ARC or their general relevance to sex differences in gene expression (e.g., androgen and estrogen receptors). We provided 10x Genomics with expression reference data for the panel design (balancing probing depth for each gene based on its expression) in the form of 1 male (v6197A_M) and 1 female (v6588B_F) Visium sample, with k=15 BayesSpace cluster labels from our preliminary analysis. A table of the final panel of genes and the rationale(s) for inclusion in the study is provided in the Supplement.

#### 3.8.2 | Data Acquisition

Data was obtained from two slides (see *Methods*, section 3.4) each in two separate runs. The first two slides of Run 1 were acquired and post-processed by the Johns Hopkins SCTC using instrument firmware/xeniumranger version 1.5. The second two slides of Run 2 assayed at SCTC were acquired and post-processed with instrument firmware/xeniumranger 1.9. Due to changes in cell segmentation algorithms between these two Xenium versions, we uniformly re-postprocessed all data by using *xenium-resegment --with-nuclei=TRUE* in xeniumranger 1.7.

#### 3.8.3 | Data QC and Preprocessing

Xenium data was initially assembled into an R *SpatialFeatureExperiment* (SFE)(*86*) object by importing xeniumranger outputs: cell boundaries, nucleus boundaries, and per-cell count data for transcript detections above quality score (QV) > 20. Cells and nuclei with segmentations containing holes as detected by the *nngeo* package were removed. Per-cell proportions of each type of negative control elements (negative control probes, negative control codewords, and unassigned/formerly BLANK codewords) were tabulated for each cell and added to the SFE. As the Xenium documentation does not address ‘Deprecated’ codewords or what they correspond to, this category of negative element was not considered further. Samples were then rotated or mirrored in space so that, for visualization, all tissue sections were oriented with the ARC in the bottom left-hand corner.

We then performed general quality control of the data as outlined in the documentation for the *Voyager* R package. Briefly, we first screened the data for nuclei that had either zero area or had an area ≥ 95% of their parenting cell’s segmentation (none were found). We then visualized the proportion of counts coming from each type of negative control element or of all elements considered jointly, confirming there were not marked spatial differences in negative element detection. We also examined the distribution of these proportions, finding that the vast majority of cells had 0 negative element reads and a small number of cells with ≥ 10% such reads. Following the *Voyager* approach, we used *scuttle::detectOutliers* to remove cells for which the proportion of reads contributed by any negative element (or their union) was > 3 median absolute deviations above the sample’s median. Cells deemed outliers in this manner also did not show any clear spatial pattern, though they were more visually dense in more cell-dense areas. Finally, we removed nuclei and corresponding cells wherein the nuclear area was under 400 pixels^2^, resulting in an initial dataset of 870,870 cells/nuclei.

Cell counts were normalized using both TMM normalization into log counts based on the number of total counts in a cell, and normalized without log transformation using *scuttle::normalizeCounts(log=FALSE)*. PCA and UMAP dimensionality reduction was performed to visually inspect for batch effects of donor and Xenium run; remarkably, the non-log-normalized data (the data used for clustering with Banksy, as advised by its authors (*87*)) occupied a UMAP space shared across these potential batch covariates (**Fig. S3**), and thus did not require batch integration for further analysis.

#### 3.8.4 | Xenium Clustering with Banksy

We subsequently set out to cluster the segmented Xenium cells using the *Banksy* package in R (*87*). Per their documentation for multi-sample datasets, we placed an artificial buffer zone between the spatial coordinates of each tissue section, such that each section had unique x-coordinates and was “spaced” at least 1.5 tissue section widths apart from every other section. We then performed *non*-spatial clustering of the Xenium data with *Banksy*, using Seurat’s non-logarithmic “relative counts” normalization as suggested in the *Banksy* documentation. We then performed *Banksy* dimensionality reduction of the relative counts with parameters *kgeom*=6 (setting the physical 6 nearest cells to be defined as a cell’s neighborhood) and the azimuthal Gabor filter setting and *lambda* both set as 0 for spatially-unaware dimensionality reduction, followed by Leiden clustering with a resolution parameter of 2. We elected to forego spatially-aware clustering in the Xenium data in order to examine *whether* genuinely distinct spatial clusters were present without biasing our analyses toward producing spatially-arranged clusters *a priori*.

#### 3.8.5 | Cell Type Annotation and VMH/ARC Domain Smoothing

We subsequently removed 8 sample/donor-specific clusters (one of which was a VMH cluster) from the SFE object to perform pseudobulk one-vs-all marker detection for each of the 33 clusters found across the dataset using *spatialLIBD* as described above for Visium. Based on marker genes, we were able to assign each cluster a putative identity, including 3 main VMH clusters and 5 ARC clusters. One ARC cluster, marked by *GAL* expression, was present both in the ARC and along the entire medial longitudinal axis of the VMH; as this cluster was not restricted to the traditional atlas boundaries of ARC (ventral to VMH and extending only along the ventromedial portion of VMH), we did not include this cluster in the smoothing procedure to define the ARC domain. For smoothing of the VMH, we excluded a fourth, rare cluster (7.3k cells total) tracing the approximate edges of VMH. We note that this cluster is annotated as “VMH boundary” in recognition of the spatial pattern, but was not marked by any canonical VMH genes (*SLC17A6*, *FEZF1*, *NR5A1*).

In order to define the spatial boundaries of VMH and ARC, we then performed a two-step spatial *k*-nearest neighbors (kNN) approach. In brief, this procedure first quantified the proportion of each cell’s 10 (VMH) or 50 (ARC) spatially nearest neighbor cells (*including* those in sample-specific clusters) that were one of the three main VMH clusters or 4 main ARC clusters. A second round of kNN, obtaining the proportion of the 200 (VMH) or 500 (ARC) spatial neighbors labeled as VMH or ARC was performed, with intermediate VMH and ARC labels assigned based on thresholding the first kNN value for each region at 0.1, 0.2…1.0 (100 VMH/ARC combinations). We collected kNN values for the second step using each first-round threshold pair, and subsequently tested final definitions of ARC and VMH with thresholds of 0.1, 0.2…1.0 as applied to the second-round kNN smoothing (total of 10,000 combinations of VMH-ARC threshold pairs).

For each of these pairs, pseudobulk DE testing was done for marker genes between all cells passing the given final VMH or ARC threshold to quantify VMH/ARC marker gene enrichment in the respectively assigned domains. DE analysis was performed using *registration_wrapper* in *spatialLIBD* as we did for other gene marker analyses. Cells from any of the 33 retained clusters falling within the boundaries of the VMH or ARC domain were given that domains label as their ‘cluster’, while any cells from a VMH or ARC cluster but found outside the two domains were removed from analysis. One-vs-all analysis was then performed with *spatialLIBD* using these modified labels such that VMH and ARC domains were treated as single clusters alongside the unmodified (non-VMH, non-ARC neuron) clusters; enrichment statistics for the VMH and ARC domains were then retrieved. First-second round combination pairs with robust enrichment of VMH markers in VMH and of ARC markers in ARC were visualized to determine whether the parameters also resulted in contiguous and directly adjacent VMH and ARC domains. For smoothing parameter combinations where VMH and ARC boundaries were adjacent, we further performed pseudobulk DE analysis to determine whether expected VMH and ARC markers were enriched on their respective sides of the boundary zone. To achieve this, we used the *sf* package to define the convex and concave hulls of smoothed VMH/ARC domains, then extended these boundaries by 5% of the median distance between hull vertices using *st_buffer* in *sf*. We performed a second round of VMH/ARC pseudobulk DE testing only comparing intra-VMH and intra-ARC cells within the overlap region.

Based on the magnitudes of enrichment for known marker genes, domain continuity, and domain adjacency, we selected first-round KNN thresholds of 0.2 (VMH) and 0.1 (ARC) followed by second-round KNN thresholds of 0.2 (VMH) and 0.5 (ARC). Cells that passed both ARC and VMH thresholds in the first or second step were deconflicted by assigning the higher KNN score (e.g., if the KNN round 1 proportion of ARC neighbors was 0.25 and of VMH neighbors was 0.3, then the cell would be labeled VMH going into the second round of KNN analysis). The labels of VMH or ARC were then assigned to all cells, regardless of cluster, to demarcate the boundaries of the VMH and ARC. Refined labels were determined by performing a one-vs-others pseudobulk enrichment approach similar to that described for Visium VMH/ARC clusters in section 3.7.5: that is, we tested pseudobulk expression on data subsetted only to ARC clusters or only to VMH clusters. This again was performed using tools from *spatialLIBD*.

Domain labels were then cleaned by removing cells outside of the main domain area that received the label; in the case of one donor, Br8741, both Xenium samples were instead processed with a second round kNN threshold of 0.15 for VMH due to holes in the domain assignments at higher thresholds. This donor contained the donor-specific VMH cluster, which was excluded from the smoothing procedure, resulting in fewer remaining VMH neurons in the sample and consequently in lower KNN proportions for VMH neuron type-neighbors.

#### 3.8.6 | Sex-Differential Expression Analyses

We set out to test each cell cluster for sex differences in expression local to the VMH or ARC. To achieve this, we first extracted all cells within either the VMH domain or the ARC domain as defined above, then performed pseudobulk aggregation within samples for each cell cluster using *dreamlet* 1.3.1. We then subsetted to clusters for which there were at least 6 samples with at least 10,000 total transcript detections for that cluster within the domain of interest. Pseudobulk counts were preliminarily normalized to total counts using TMM normalization. We then programmatically set a low-expression filter by detecting the smaller sex group (in terms of sample number) among the retained samples for a given cluster in a given domain by detecting the second minimum in a histogram of gene-wise mean log counts per million to retain only those genes which adhered to an approximately normal distribution. Genes falling below this second minimum were discarded and TMM factors recalculated for the remaining genes. DE analysis was then performed with voomLmFit, using donor as a blocking factor (which is internally passed to *DuplicateCorrelation*), *adaptiveSpan* set to true to identify the number genes to use for smoothing the mean-variance trend, *sample.weights* set to true to perform an analysis analogous to the *voomLmFit* predecessor *voomWithQualityWeights*, and fitting a model of *∼0+Sex+Xenium Run* to test for sex effects and control for potential variation between the two Xenium runs. The filtered pseudobulk object and full DE analysis output for each cluster within each domain were returned for visualization and downstream analyses. This same approach was also taken to conduct Visium-analogous, domain-level sex-DE analyses, treating cells with any of the 33 retained cluster labels and inside the VMH or ARC domain as “VMH” or “ARC”, respectively.

In order to validate our sex-DE findings within VMH and ARC domains, we performed two additional high-confidence analyses. In the first, we utilized the *sf* package in R to define the convex and concave hulls for VMH and ARC, respectively, and trimmed these boundaries back by 5% of their respective domain areas to obtain “high-confidence” domains. Cells falling within these interior domain boundaries were then analyzed as above for sex-DE. In a second analysis, we sought to rule out cell segmentation effects that could have resulted in transcripts being incorrectly attributed to a given cell type (e.g., astrocyte endfoot processes could potentially end up within neuronal segmentation boundaries). We therefore also performed sex-DE only considering transcripts overlapping nucleus segmentations by assigning a given nucleus the same cluster label as its parent cell segmentation. The nuclear count data was filtered to quality scores > 20 and totaled per gene per nucleus using the *MoleculeExperiment* package in R (*88*), followed by the same pseudobulk per-cluster-per-domain procedure as described above.

## Results

### Molecular identification of human VMH and ARC using transcriptome-wide spatial transcriptomics

To understand sex-differentiated gene expression at cellular and spatial resolution in the human VMH and ARC, we performed Visium spatial gene expression profiling followed by Xenium *in situ* sequencing (10x Genomics) on adjacent sections of postmortem human HYP tissue from male and female adult neurotypical donors (*N*=4 per sex, BMI range for males 23.1-28.7, 24-31.6 for females; **Fig. 1A-C**). We used Visium to measure transcriptome-wide gene expression across the VMH and ARC, facilitating data-driven selection of genes for expression profiling at cellular resolution using Xenium (**Fig. 1C**). To locate the VMH and ARC, we performed single-molecule fluorescence *in situ* hybridization (smFISH) for *NR5A1* and *NPY,* marker genes for VMH and ARC, respectively. When the VMH and ARC were reached, tissue sections were first collected for Visium (10 total samples, 1-2 sections per donor) (**Table S1**); sections from these same tissue blocks were later obtained for Xenium experiments. After quality control (**Fig. S2**, **Fig. S4**), the filtered Visium dataset included ∼45K spots and ∼30k genes.

To identify data-driven spatial domains representing VMH and ARC from Visium, we performed unsupervised spatial clustering. Specifically, we used 1,816 spatially variable genes (SVGs, **Table S2**)(*61*) as features for dimensionality reduction and batch correction with Harmony (*62*) (**Data S1**), followed by unsupervised clustering at *k*=15, 20 or 31 (*Methods*). We present results using *k*=15 (**Fig. 1D**) as this value is consistent with the number of anatomical regions annotated in the Allen Brain Atlas in our samples. At *k*=15, we discerned 2 oval-shaped VMH clusters and two ARC clusters ventral to them (**Fig. 1E**). VMH clusters uniquely expressed the glutamate transporter *SLC17A6* and signature VMH marker genes *FEZF1* and *NR5A1* (**Fig. 1F**; **Fig. S1**, **Fig. S5**). ARC clusters were enriched in *KISS1*—a regulator of gonadotropin releasing hormone (GnRH)—as well as neuropeptide-encoding genes *GHRH*, *AGRP*, *NPY*, and *POMC* (**Fig. 1G**, **Fig. S5**, **Table S3A-C**). We henceforth refer to these Visium (v) clusters as vVMH.1, vVMH.2, vARC.1 and vARC.2, and when considered together as ‘vVMH’ and ‘vARC’.

We examined our two vVMH clusters to determine whether they contained spatially-restricted cell types. Pairwise differential expression between vVMH clusters demonstrated that vVMH.1 was more enriched for canonical marker genes, while vVMH.2 was more enriched for glial markers (*e.g.*, *GFAP*, *MBP*; 0 < logFCs < 0.75; **Fig. S5**, **Table S4**). This was also the case for vARC, suggesting that vVMH and vARC clusters reflect local differences in glial content rather than spatially exclusive neuronal populations *per se*. In neither vVMH nor vARC did higher *k* values yield further molecularly distinct (neuronal) clusters inside ARC or VMH domains (**Fig. S5**, **Fig. S6**, **Fig. S7**). Therefore, we joined vVMH.1 and vVMH.2 into a single vVMH cluster and did the same for vARC for downstream analyses.

To examine how well vVMH and vARC domain marker genes corresponded to those of the adult mouse VMH and ARC, we compared our human data to a recently published single cell atlas of the adult mouse brain (*85*). vVMH, vARC, and optic tract (OT) clusters shared gene marker signatures with mouse VMH, ARC, and white matter cell types, respectively (**Fig. 1**, **Table S5E**). We obtained similar results when comparing marker gene expression against adult mouse *in situ* hybridization data from the Allen Brain Atlas (*84*) (**Table S5A-D**).

We next performed gene set enrichment analysis (GSEA) for vVMH and vARC using MSigDB-indexed gene sets (*Methods*) (**Fig. 1I**; full results of this analysis at *k*=15 (with vVMH and vARC as well as their individual clusters), *k*=20, and *k*=31 are available in **Data S2**). As expected, vARC was enriched for inhibitory neuron terms, vVMH for excitatory neuron terms, and both for terms related to neuropeptides. vVMH-enriched genes were overrepresented for kisspeptin signaling pathways, which is consistent with previous literature demonstrating VMH responses to kisspeptinergic neuron activity (*89*), and with our observed expression of kisspeptin receptor (*KISS1R)* in vVMH (**Table S3A-C**). Further, “regulation of insulin secretion” genes were enriched among highly expressed genes in vVMH, consistent with its glucoregulatory function in rodents (*90*),(*91, 92*). Interestingly, vARC-enriched genes were overrepresented for terms relating to primary cilia and their organizing body, the axoneme. We confirmed this by performing GSEA with a curated set of human primary cilia genes (*93*), and found ARC-specific enrichment for primary-cilium-localized genes (FDR<5x10^-7^; **Table S6I**). Taken together, these results show that we successfully identified ARC and VMH in our human Visium samples, and that the signals, circuits, and cellular biology in these regions are broadly consistent with expectations from the rodent literature.

### Correlated molecular sex differences in VMH and ARC domains

To assess transcriptome-wide sex-differential expression in vARC and vVMH and identify genes of interest for cell type-localization with Xenium, we performed pseudobulk DE analyses, which revealed 11 significant vVMH (**Fig. 2A**) sex-DEGs, all of which were allosomal. Orthologs to sex-DEGs previously identified in mouse VMH (*Cckar*, *Rprm*, *Pdyn*)(*8*) did not achieve nominal significance in human vVMH, though we did note nominally significant male-biased expression of melanocortin receptor *MC4R* (**Fig. 2B-C**), which has been previously described to play roles in VMH-mediated metabolic regulation (*94*, *95*). Meanwhile, we identified 26 significant vARC (**Fig. 2**) sex-DEGs (FDR<0.05, **Table S7**), 13 of which were autosomal. These included female-biased expression of *RDH10*, which encodes an enzyme responsible for converting retinoids into active ligands for retinoid receptors (**Fig. 2**). The distribution of p-values for both analyses showed an overrepresentation of p-values close to zero, suggesting a sizable number of ‘true’ differentially expressed genes in both domains, higher than the 11 or 26 we report here (**Fig. S8A-D**).

**Fig. 2.**
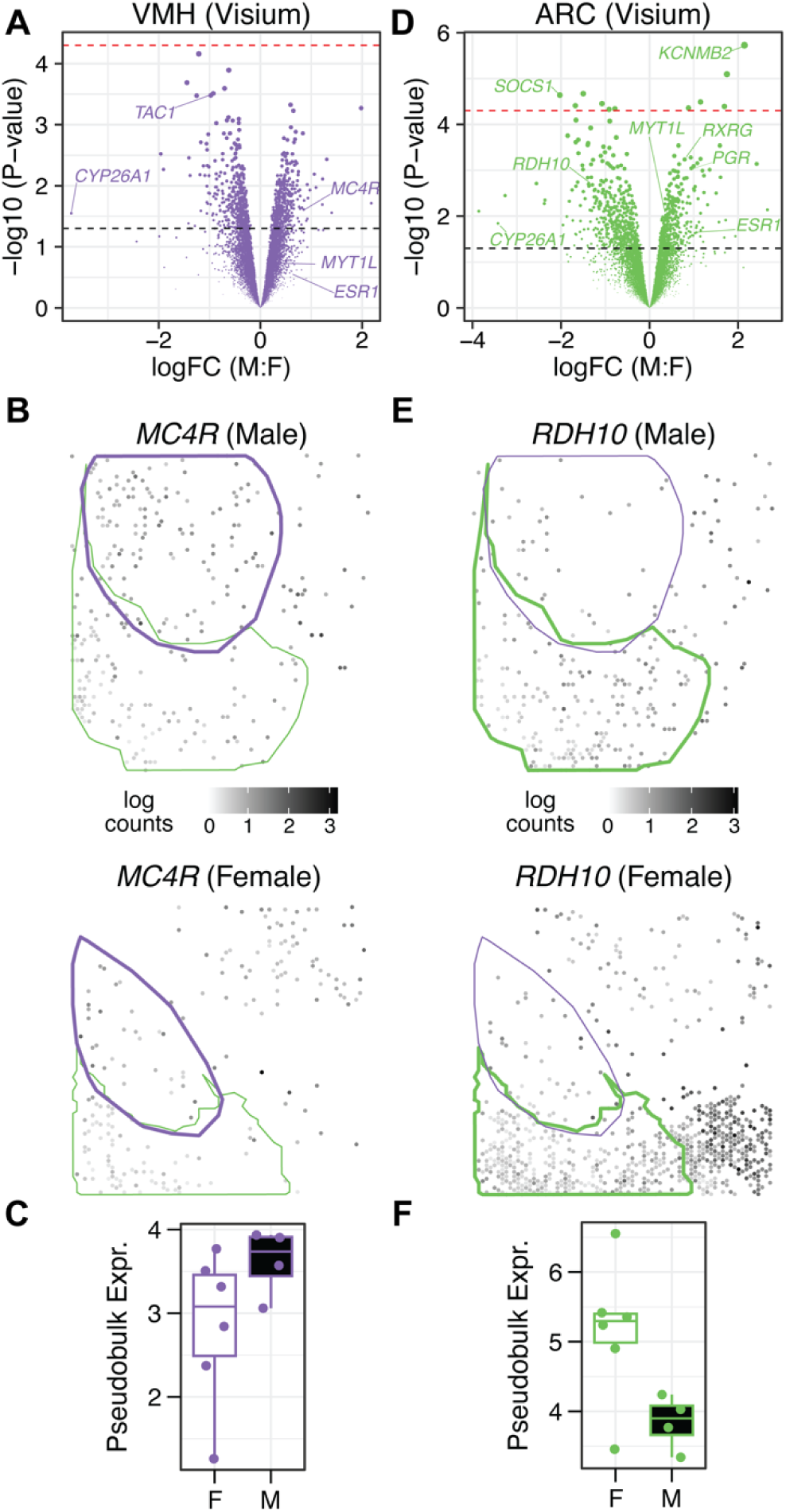
Sex-differential expression (sex-DE) at the domain level in the human VMH and ARC. (**A**) Volcano plot illustrating autosomal sex-DE within vVMH. The black line indicates nominal *p*<0.05; the red line indicates FDR<0.05. (**B**) Spotplots from one male (upper; v6197A_M) and one female (lower; and v6588A_F) Visium sample illustrating the nominally significant male-upregulation of *MC4R* in vVMH (thicker domain border colored in purple). (**C**) Pseudobulk domain-level expression (log2 counts per million (CPM)) of *MC4R* in all vVMH samples, stratified by sex. (**D**) Volcano plot illustrating autosomal sex-DE within vARC. Lines as described for panel A. (**E**) Spotplots from the same male (upper) and female (lower) Visium samples as in B, illustrating female-preferential expression of *RDH10* in ARC (thicker domain border colored in green). (**F**) Pseudobulk domain-level expression (log2 counts per million (CPM)) of *RDH10* in all vARC samples, stratified by sex. Sample identifiers are given as platform (v or x), donor number (4 digits), replicate tissue section (A-D), followed by an underscore and donor sex.

We observed a strong concordance between autosomal sex-differential expression in vARC and vVMH, with greater fold changes in vARC. Effects of sex were correlated (*t*-statistic Spearman rho=0.542; logFC Spearman rho=0.585; **Fig. S8A-B**), and the correlation increased when we restricted to genes nominally DE in either of the two domains (t-statistic rho=0.586; logFC rho=0.683). Among nominal autosomal sex-DEGs in Visium, 211 were found in both domains, of which 210 had concordant effects. The one exception was histamine H1 receptor (*HRH1*), which showed greater vARC expression in females (logFC 0.77) and greater vVMH expression in males (logFC=0.49). Analysis of the p-value distribution of vVMH conditional on vARC (and vice versa) likewise suggested a substantial overlap among the sex-DEGs in the two domains (**Fig. S8E-F**).

### Cell-level expression of sex-DE genes in VMH and ARC with imaging-based spatial transcriptomics

Following an analysis of preliminary Visium data, a unique panel of 100 genes was designed for the Xenium platform (**Table S8**). This panel included vVMH and vARC marker genes, 36 putative sex–DEGs identified with Visium, sex hormone receptors (the androgen receptor, *AR*; nuclear estrogen receptor *ESR1*; and a G-protein coupled estrogen receptor, *GPER1*), and genes with spatial or sex patterns from rodent HYP literature (*Methods*). We additionally performed spatially-variable gene (SVG) analyses of vVMH or vARC in isolation to identify expression patterns suggestive of fine-grained cellular organization in these domains. We identified 277 SVGs in vVMH and 1279 SVGs in vARC (**Table S9**); among genes in the latter set were 11 markers of 6 ARC clusters previously reported in developing human HYP, including *TAC3, GHRH,* and *DIRAS3* (*96*). All 11 of these ARC cluster markers were significant SVGs in ≥6 tissue sections, with mean ranks skewed toward the top (Wilcoxon test, p<1x10^-7^), and 8/11 were in the top 150 vARC SVGs overall. We thus concluded that genes with strong SVG signals and consistent patterns within a single Visium domain were likely indicative of discrete neuron types not resolvable as distinct clusters using Visium; therefore, several of these SVGs were included in the Xenium panel (**Fig. S9-Fig. S17**). Our 100 custom-selected genes were supplemented by a commercially available “human brain” panel of 266 genes spanning markers for broad cell types largely derived from human cortex data. Since our goal was to study sex differences, we designed our experiment such that each processing batch contained an equal number of males and females. This ensures that any unwanted variation associated with the processing batch is independent of sex differences.

From all 8 donors, tissue sections adjacent to those profiled with Visium were collected for Xenium (*N*=7 male and *N*=6 female sections). We balanced each Xenium slide to contain sections from both sexes, and analyzed two slides per Xenium run. This ensures that any unwanted variation associated with slide (and processing batch) is independent of sex differences. After quality control (**Fig. S18**), data were clustered (**Fig. 3A**, **Fig. S3**), yielding 33 clusters total, including 5 Xenium ARC (xARC) clusters and 3 Xenium VMH (xVMH) clusters identifiable by marker genes and spatial distributions (**Fig. S3**, **Fig. S21A-B**, **Table S3G**). All three xVMHclusters shared enrichment for a number of VMH marker genes (*SLC17A6*, *NR5A1*, *FEZF1*, *NRGN*, *ANKRD34B*, *ADCYAP1*, *HS3ST4*; **Fig. S21B**). Comparison across these three clusters revealed that xVMH.1 was highly enriched for *CRHR2*, known to be expressed in rodent VMH (*97*, *98*). Interestingly, xVMH.2 was comparatively enriched for the oxytocin receptor (*OXTR),* but also for astrocyte (*GJA1*, *SOX9*) and microglia (*P2RY13*) markers (**Fig. S21C**, **Table S3I**); as such, we refer to xVMH.2 as “VMH-glia mixed.” xVMH.3 was enriched for *LAMP5*; as *LAMP5* is associated with inhibitory neurons in other brain regions (in contrast to the VMH, which is composed of excitatory neurons), we performed smFISH to confirm *LAMP5* colocalization with excitatory neuron marker *SLC17A6* and VMH marker *NR5A1* (**Fig. S19**, **Fig. S20**). A fourth, *HCRT*-expressing cluster showed striking restriction to the VMH perimeter; we grouped this “lateral border” population as a fourth xVMH cluster in downstream analyses based on its surprising spatial pattern (**Fig. S22**).

**Fig. 3.**
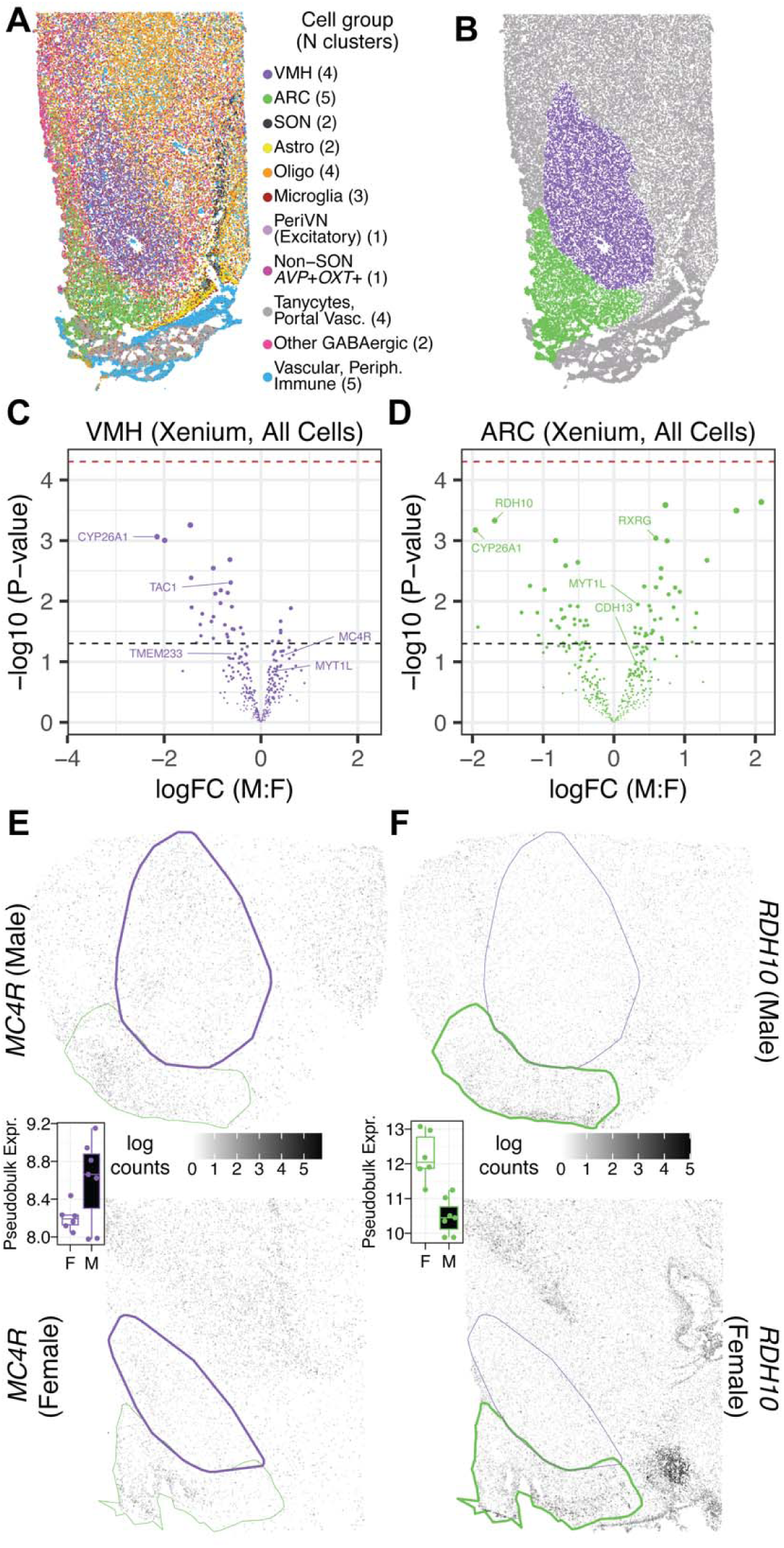
Xenium clustering, domain identification, and sex DE analyses at domain level. (**A**) Xenium cell type clustering from the same donor (sample x1225B_M) as shown in Figure 1D, with clusters combined into Visium-analogous groups for visualization. Legend notes the number of cell type clusters in the visualized group. (**B**) Xenium (x)VMH and xARC anatomical boundaries as inferred using *k*-nearest neighbors analyses to quantify the number of spatial neighbor cells from xVMH or xARC clusters. (**C**) Volcano plot illustrating autosomal sex-DE within xVMH (all cells from all clusters and within xVMH). The black line indicates nominal *p*<0.05; the red line indicates FDR<0.05. (**D**) Volcano plot illustrating autosomal sex-DE within xARC (all cells from all clusters and within xARC). Lines as in panel C. (**E**) Expression spotplots of example male (above) and female (below) Xenium samples x6197B_M and x6588C_F, illustrating male-preferential expression of *MC4R* in VMH (thicker domain border colored in purple). Boxplot inset shows pseudobulk domain-level expression (log2 counts per million (CPM)) of *MC4R* from each xVMH sample stratified by sex. (**F**) Xenium samples from the same representative donors as in E, illustrating female-preferential expression of *RDH10* in ARC (thicker domain border colored in green). Boxplot inset show pseudobulk domain-level expression (log2 counts per million (CPM)) of *RDH10* in each xARC sample, stratified by sex. Sample identifiers are given as platform (v or x), donor number (4 digits), replicate tissue section (A-D), followed by an underscore and donor sex.

Of our 5 xARC clusters (**Fig. S23**), 3 were readily identifiable based on known marker genes from the literature: xARC.1 as *AGRP*-expressing neurons, xARC.3 as a *TRH*-expressing subpopulation of *GHRH* neurons, and xARC.4 as *TAC3-* and *ESR1*-expressing neurons, possibly representing *KISS1*-expressing KNDy neurons (**Fig. S21**). For the two remaining clusters, xARC.5 was comparatively enriched in oligodendrocyte markers (*OLIG1*, *OLIG2*, *PDGFRA*) along with *POMC*, and thus designated “*POMC*-Oligo Mixed,” while xARC.2 was enriched for *SLC17A7* and glial markers and designated “*SLC17A7*-Glia Mixed” (**Fig. S21C**, **Table S3H**). Regarding relative enrichments of glial genes when comparing among xARC or xVMH clusters, we repeated our analysis using only nuclear-expressed transcripts and obtained similar results (**Table S3J-K**), suggesting close interactions between glial processes and neurons in select clusters.

We then sought to examine whether Xenium sex-DE patterns were consistent with Visium at the domain level. To achieve this, we first defined ARC and VMH domains analogous to those from Visium by developing a smoothing algorithm (*Methods*) to obtain anatomically contiguous domains corresponding to the xARC and xVMH (**Fig. 3B**). Consistent with Visium, Xenium domains for xVMH and xARC were respectively enriched for expected marker genes *NR5A1* and *POMC*, with qualitatively strong anatomical agreement within donors and across platforms (**Fig. S24**). With xARC and xVMH defined, we pseudobulked across all cells within xVMH or xARC and performed sex DE analyses, covaried by Xenium run (**Fig. 3C-D**), to assess cell type-agnostic sex DE (analogous to vVMH/vARC sex-DE analyses). While no genes achieved significance at an FDR<0.05 in Xenium domains, trends identified in Visium were replicated, such as male upregulation of *MC4R* in VMH (**Fig. 3E**) and female upregulation of *RDH10* in ARC (**Fig. 3F**). Systematic examination of sex DE across platforms revealed correlated autosomal effects of sex (*t* statistics; ARC, 277 genes, Spearman rho=0.63, p<2.2x10^-16^; VMH, 245 genes, Spearman rho=0.41, p<5.5x 10^-11^; **Fig. S8G**). The correlation increased when only considering the 36 genes earmarked for sex-DE follow-up on Xenium (ARC, 33 genes, rho=.80, *p*<5.8x10^-7^; VMH, 27 genes, rho=0.51, *p*<0.01; **Fig. S8H**). These findings suggest that sex effects at the domain level were consistent across platforms, regardless of DE significance.

Additionally, we examined domain-level expression of estrogen, progesterone, and androgen receptors using both platforms. As anticipated from rodent and human data (*99*), *ESR2* expression was weaker relative to *ESR1* in VMH and ARC (**Fig. S25A**). Visium expression of progesterone receptor (*PGR*) was moderate in both vVMH and vARC, consistent with its protein expression pattern in human HYP (*100*). *PGR* was also nominally sex-DE (male-elevated) in vARC (logFC 0.94, *p*<0.001) and trended similarly in vVMH (logFC 1.23, p=0.053; **Table S7**). *PGR* (Visium only), *ESR1*, and *AR* were detected in a greater fraction of ARC spots/cells compared to that of VMH. Contrary to expectations from rodent literature showing enrichment of *ESR1* in the vlVMH (*2–4*, *8*, *10*, *21*, *22*), *ESR1* expression was sparse and spatially uniform in human xVMH. smFISH on Visium-adjacent tissue sections reproduced this finding (**Fig. S26A**; see *Data Availability* for full-tissue *ESR1* smFISH microscopy). The dense expression of *ESR1* in human ARC compared to VMH further suggests that our observations of more marked sex-DE in the ARC domain are driven by hormonally-mediated (activational) signaling.

### Single-cell analysis reveals especially marked sex effects in ESR1-expressing neurons of ARC

Next, to determine whether VMH and ARC showed cell type-specific sex differential expression, we examined all Xenium-measured genes for sex-DE within individual cell types in xVMH or xARC domains (**Fig. S27**, **Table S10A-B**). In xVMH, we found the most significant (FDR<0.05) sex-DEGs in excitatory *CRHR2*-expressing cells, involving 39 genes, with another six DEGs in the xVMH-Glia mixed cluster (**Fig. 4A**, **Fig. S28**). Among these, *TAC1* was female-upregulated in the xVMH-Glia mixed cluster with strong trends (FDR<0.1) towards female upregulation in *CRHR2* and *LAMP5* clusters—consistent with sex-DE of *Tac1* observed in mouse VMH (*8*, *101*)—and *CYP26A1* trended toward female upregulation in *CRHR2* and xVMH-Glia mixed clusters (**Fig. 4B**). Male xVMH *CRHR2* neurons showed significantly greater expression of *MC4R* (**Fig. 4B**), an obesity-associated (*102*) melanocortin receptor, suggesting the *CRHR2* population of xVMH drives the observed domain-level sex DE. Likewise, *MYT1L*, a transcription factor (TF) mutated in a syndromic form of ASD involving obesity (*103*), was selectively male-upregulated in the *CRHR2* population (**Fig. 4B**).

**Fig. 4.**
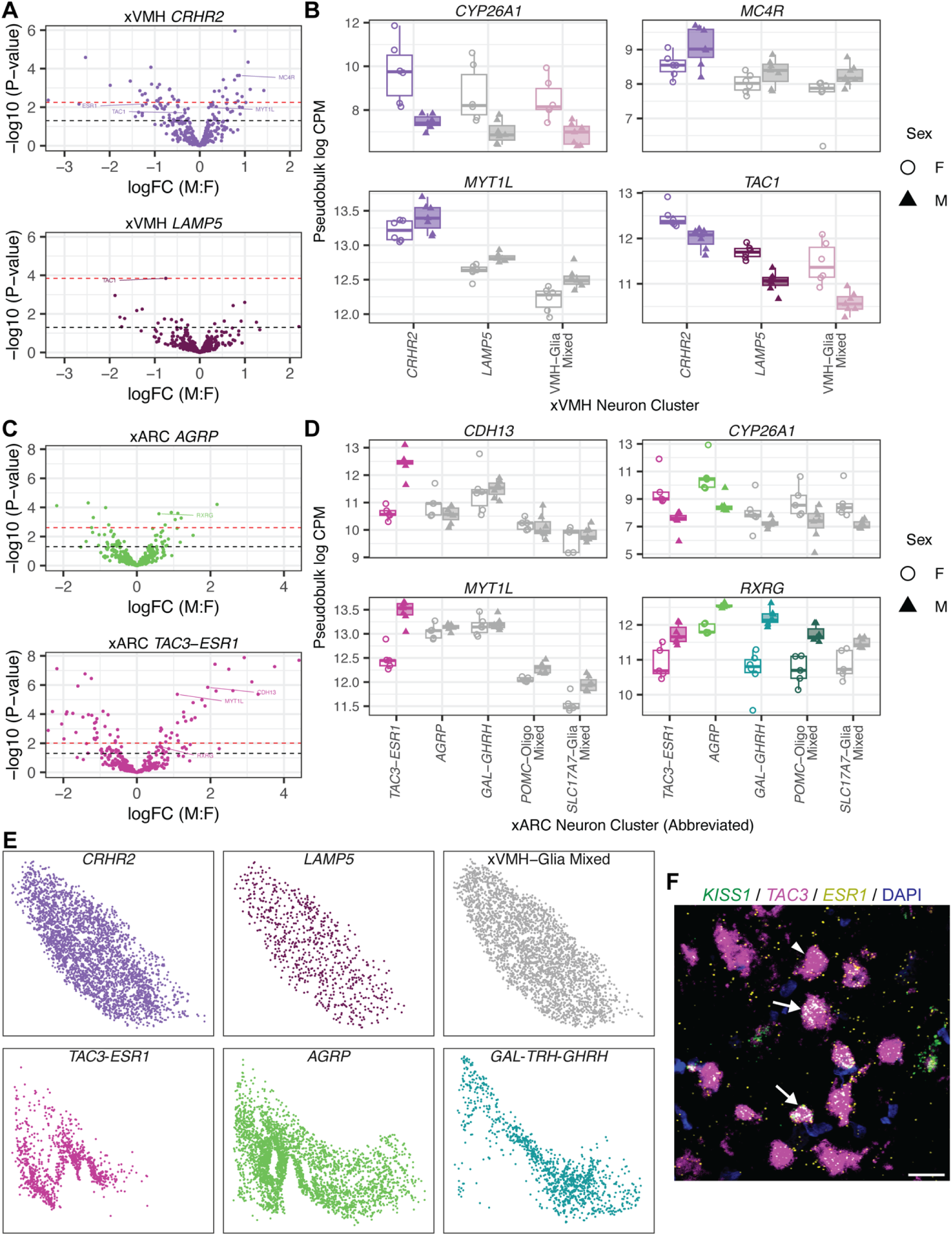
Sex-DE within Xenium-resolved cell types of the human xVMH and xARC. (**A**) Volcano plots of sex-DE for selected xVMH cell clusters. (**B**) Top candidate sex-DE genes from vVMH measured using Xenium. Each point represents the pseudobulk intra-xVMH gene expression for a cluster in one sample (color legend shared with panel A), stratified by sex. Cell types trending toward sex-DE (FDR<0.1) are colorized in this plot. (**C**) Volcano plots of sex-DE for selected xARC cell clusters highlighting that sex-DE is most prominent in terms of significance and effect size in *TAC3*-*ESR1* cells. (**D**) Top candidate sex-DEGs identified in vARC measured using Xenium. Each point represents the pseudobulk intra-xARC gene expression for a cluster in one sample (color legend shared with panel C), stratified by sex. Cell types trending toward sex-DE (FDR<0.1) are colorized in this plot. (**E**) Visualization of individual xVMH/xARC clusters wherein trending sex DE was most marked. Top row illustrates individual xVMH clusters in shared x-y space from one Xenium sample (x6588C_F); bottom row illustrates individual xARC clusters in the same manner. As with other spatial plots, down is ventral, left is medial. (**F**) Confocal image showing expression of *ESR1*, *TAC3*, and *KISS1* in an adjacent tissue section by smFISH in a region of ARC expressing all 3 genes (scale bar 25µm). Arrows indicate cells expressing all three genes of interest. Arrowhead indicates an example cell with only *TAC3* and *ESR1* expression. Sample identifiers are given as platform (v or x), donor number (4 digits), replicate tissue section (A-D), followed by an underscore and donor sex.

Sex-DE (FDR<0.05) in xARC was most extensive in *TAC3*-*ESR1* neurons (67 genes, **Fig. 4C**), and detected in all xARC populations except the *POMC*-Oligo mixed cluster (**Fig. S28**). As among xVMH cell types, we found male upregulation of *MYT1L* to be cell type-specific in xARC, only observed in *ESR1*-*TAC3* cells (**Fig. 4D**). *TAC3*-*ESR1* neurons also showed specific and marked male upregulation of *CDH13*, a receptor for insulin-sensitizing peptide adiponectin (*104*) implicated in obesity and type 2 diabetes (*105–107*) (**Fig. 4D**). Retinoid receptor *RXRG,* a male-upregulated DEG in vARC, showed less cell type-selective sex-DE, with significant male-upregulation in three xARC clusters and a trend toward the same in *ESR1*-*TAC3* neurons (**Fig. 4D**). Consistent with domain level findings, retinoid metabolism genes were female-upregulated in xARC, with *RDH10* showing DE in four populations (**Table S10A-B**) and vARC marker *CYP26A1* showing significant or trending DE in two xARC clusters (**Fig. 4D**). No significant DEGs showed opposite directions of effect among or across xARC and xVMH cell types. However, in agreement with vARC and vVMH findings, *HRH1* was significantly upregulated in two xARC clusters from females while trending toward male-upregulation (*p*<0.07) in xVMH *CRHR2* cells.

We next examined sex hormone receptor expression patterns at the finer resolution of individual cell clusters (**Fig. S25B**). The greatest proportion of cells with ≥1 count of *ESR1* included xARC *TAC3*-*ESR1*, xARC *AGRP*, and xVMH *CRHR2—*all populations notable for having a number of significant sex-DEGs—in addition to cells of microglia cluster 1 and oligodendrocyte progenitors (OPCs) residing in either xVMH or xARC. *AR* was generally found in a similar proportion of cells for each cluster-domain pair, with the notable exception of xARC *GAL*-*TRH*-*GHRH*, where it was detected in ∼2-fold more cells than was *ESR1*.

To confirm that our sex-DE findings were not confounded by ARC/VMH cell type intermixing at domain boundaries, we repeated the sex-DE analyses by reducing both the xVMH and xARC domain boundaries inward by 4%, which gave well-correlated t-statistics relative to the original domain boundaries for all 5 xARC clusters and the 3 main xVMH clusters (all Spearman rho > 0.99) (**Table S10F-H**). Similarly, we ruled out effects of cell segmentation by repeating the analyses using only nuclear transcript counts, which also yielded consistent results (**Table S10D-E** and **H**).

As several instances of sex-DE in xARC or xVMH were cell type-specific, we next examined whether these cell types displayed spatial organization that would suggest a specific anatomic subdomain where sex effects were most pronounced. xVMH clusters showed no spatial organization, with each occupying the entirety of their domain (**Fig. 4E**). xARC, on the other hand, showed exquisite spatial isolation of *TAC3*-*ESR1* cells in particular. Consistent with Visium SVG results (**Fig. S14**-**Fig. S15**), *GHRH* and *TAC3* expression was mutually exclusive within xARC (**Fig. S29**). Given both spatial and sex-DE features of our *TAC3*-*ESR1* cluster, we sought to more definitively annotate these cells. Although *KISS1* was not included in our Xenium panel, smFISH showed that ∼14% of *ESR1*+/*TAC3+* neurons in ARC also express *KISS1*, suggesting that KNDy neurons are included within this population (**Fig. 4F**, **Fig. S26**). Additionally, orthologues to secondary markers of mouse KNDy neurons (*108*, *109*) were also enriched in our *ESR1*-*TAC3* population (*ECEL1* and *NR4A2*, **Fig. S21**, **Table S3G**). These findings highlight a previously unrecognized ARC subdivision in which *ESR1*-*TAC3* neurons (both *KISS1*-positive and -negative), and thus much of sex-DE in the adult ARC, are restricted. Altogether, we replicated and refined several Visium findings at higher spatial resolution and identified specific xARC and xVMH cell types responsible for sex-DE signals. Moreover, we identified cell populations under the strongest transcriptional influence of sex in each domain and illustrate that genes—including those linked to known VMH/ARC functions such as metabolism and obesity (*e.g.*, *MC4R*, *CDH13*)— are influenced by sex in the human VMH and ARC.

### Male upregulated genes in VMH and ARC are enriched for neurodevelopmental disorders

We next sought to leverage our transcriptome-wide data to identify potential transcriptional regulators of adult vVMH- and vARC-defining expression patterns. To achieve this, we performed GSEA on one-vs-all marker analysis test statistics in each Visium domain using aggregated databases of putative transcription factor (TF) target genes across mammalian tissues and cell types (**Table S6**, **Data S3**; see *Methods*). Gene targets of the glucocorticoid receptor (*NR3C1*) and of a putative retinoic acid receptor (*NR2F2*) were enriched in the vVMH (*110*). Similarly, targets of the traditional retinoid receptors (retRs) *RXRA* and *RARA* were enriched in vVMH and vARC, respectively (**Fig. 5A**). Finally, several results suggested sex differences in vVMH and vARC, including androgen receptor (*AR*) target enrichment in VMH and *AR* interactors (*i.e.*, co-TFs)(*111*) target enrichment in ARC. Furthermore, genes upregulated or downregulated by perturbation of *EGR1,* a TF that sex-differentially modifies chromatin in the hippocampus (*112*), were respectively enriched in vARC or vVMH. These findings collectively highlight previously underappreciated signaling pathways in the human VMH and ARC.

**Fig. 5.**
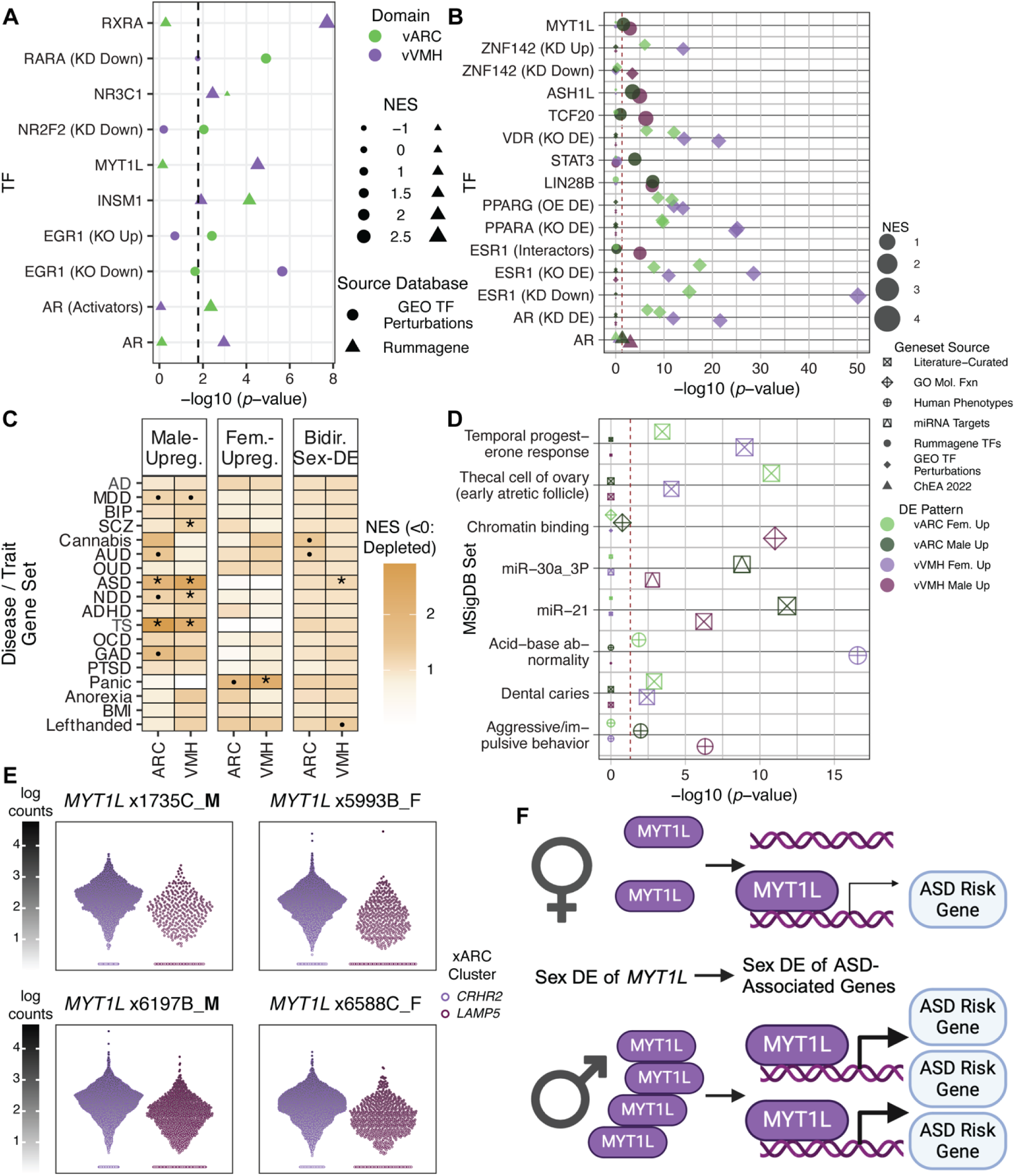
Sex-DE patterns of vARC/vVMH are enriched in ASD/NDD risk genes, including disease-associated TFs. (**A**) GSEA enrichment of vVMH/vARC markers, weighted by one-vs-all marker *t* statistics in neuropsychiatric disorder- and BMI-associated genes. * FDR<0.05; • uncorrected *p*<0.05. (**B**) GSEA enrichment of genome-wide vVMH/vARC sex-DE, weighted by signed DE *t* statistics, in neuropsychiatric disorder- and BMI-associated genes. Tests of enrichment for patterns of male and female upregulation are shown, along with a test considering sex-DE in both directions. * FDR<0.05; • uncorrected *p*<0.05. (**C**) GSEA enrichment of Visium sex-DE for ontologic and functional terms indexed in MSigDB, with points sized based on normalized enrichment score (NES). Legend shared with panel D. (**D**) GSEA enrichment of Visium sex-DE for TF targets/TF interactors, with points sized based on normalized enrichment score (NES). Multiple points are illustrated for a sex-domain pair for GEO TF perturbation sets reporting multiple experimental replicates. (**E**) ASD-associated TF, MYT1L, whose targets are overrepresented in genes with male-preferential vARC and vVMH expression (**D**), is also male upregulated in xVMH, predominantly in the *CRHR2* population. Cell-level expression is plotted for two samples per sex (sex denoted in sample name) for xVMH clusters *CRHR2* and *LAMP5*. (**F**) A hypothesized mechanism that may connect our observations of sex-DE genes, shared TFs regulating them, and sex-DE enrichment for ASD genes. *MYT1L* (itself an ASD risk gene) is a male-upregulated TF in our data (left side); likewise, male-upregulated genes in VMH are overrepresented in *MYT1L* regulatory targets, including ASD risk genes (as discussed in *Results*; right side), suggesting that at the level of domains or cell populations, sex-DE of TFs like *MYT1L* may also drive sex-DE of several further disease risk genes. AD: Alzheimer’s disease; ADHD: attention deficit-hyperactivity disorder; ASD: autism spectrum disorder; AUD: alcohol use disorder; BIP: bipolar disorder; BMI: body mass index (quantitative trait); Cannabis: cannabis use disorder; GAD: generalized anxiety disorder; MDD: major depressive disorder; NDD: neurodevelopmental disorder; OCD: obsessive-compulsive disorder; OUD: opioid use disorder; PTSD: post-traumatic stress disorder; SCZ: schizophrenia; TS: Tourette’s syndrome.

We then performed a similar GSEA analysis examining sex-DEGs and the same TF-target gene sets, again filtering results to TFs expressed in vVMH or vARC (see *Methods*) (**Fig. 5B**, **Data S4**). Sex hormone receptor targets were enriched for DE in both ARC and VMH. While *ESR1* target sets were enriched strongly in female vVMH and vARC, male vVMH was weakly enriched for one target set and for ESR1 protein interactors. Female vVMH showed stronger enrichment for inferred androgen receptor targets than did male vVMH, presumably representing female expression of genes repressed by AR activity. Finally, GSEA findings indicated enrichment of vARC sex-DEGs for *AR* targets specifically upregulated via hormone-independent *AR* activator (*113*) and chromatin modifier (*114*) *PKN1* (**Data S5**).

Given that the prevalence rates for many neurodevelopmental (NDD) and neuropsychiatric disorders are sex biased and that the HYP plays established functional roles in behaviors relevant for these disorders, we also hypothesized that expression patterns defining VMH and ARC may be enriched in NDD and neuropsychiatric disorder gene sets. Toward this end, we collected gene sets from genome-wide association studies (GWAS), rare-variant studies (where applicable), transcriptome-wide association studies (TWAS) (*77–81*), and the database DisGeNET (*82*), which includes curated disease-gene links (**Table S6**). These gene sets were combined into a single gene set for each of 18 disorders. We then performed GSEA to identify whether disease-associated genes demonstrated marker-like expression or sex-DE in vARC or vVMH. vARC did not show an enrichment of genes associated with neuropsychiatric disorders, while vVMH showed enrichment of genes associated with several sex-biased disorders, including autism spectrum disorders (ASDs), neurodevelopmental disorders (NDDs), and schizophrenia (SCZ) (**Table S11**). A similar GSEA on sex-DE patterns in vVMH or vARC revealed that male-upregulated genes from both vVMH and vARC were overrepresented in genes associated with risk for ASDs and Tourette’s syndrome (TS), which are disorders more prevalent in males (**Fig. 5C**, **Table S7E**). While our Visium analysis only identified three ASD-associated genes with nominal sex-DE in vARC, *TAC3*-*ESR1* cells in xARC showed sex-DE (FDR<0.1) for 11 such genes (*AR*, *CDH13*, *DCC*, *GABRA5*, *GABRQ*, *GLRA2*, *MYO16*, *MYT1L*, *NR4A2*, *PCLO*, *THBS1*; **Table S12**). *MYT1L*, a gene mutated in a syndromic form of ASD involving obesity (*103*), was significantly male-upregulated selectively in *ESR1*-*TAC3* neurons, providing previously unreported cell type context for this risk gene. These findings exemplify how gene panel prioritization using Visium improves our ability to detect sex-DEGs in anatomically-restricted cell types, while retaining transcriptome-wide information to link such genes to broader pathways.

We then asked whether vVMH or vARC sex-DE corresponds to broader aspects of HYP biology that may be relevant to the enriched neuropsychiatric disorders, or to gonadal hormone signaling, by performing similar GSEA analyses of sex-DE patterns with pathway and ontology gene sets (**Data S5**). Male-biased vVMH genes were enriched for gene sets associated with aggression, which is consistent with behavioral functions of male rodent VMH in regulating aggressive behavior (*21*, *23*). On the other hand, male-biased vARC genes were enriched for targets of *miR-30a*, a regulator of puberty onset downregulated by perinatal estrogen exposure in female rodents (*115*) (**Fig. 5D**). Progesterone-responsive genes were enriched among female-biased genes of vVMH (**Fig. 5D**), despite trends toward greater male *PGR* expression in both vVMH and vARC.

Last, we examined individual sex-DEGs on Xenium at FDR<0.1, focusing on 8 neuropsychiatric/neurodevelopmental disorders with prominent sex differences in prevalence and/or clinical features (ADHD, anorexia nervosa, ASD, MDD, Tourette’s syndrome, SCZ, generalized anxiety disorder and panic disorder). In total, we obtained 122 disease-associated genes with sex-DE in ≥1 cluster within ARC and 35 in ≥1 cluster within VMH, with 29 genes in common between the two domains. These clusters include glial cells, interneurons, and other broad cell types that may interact differently with VMH- and ARC-specific neuronal populations (**Fig. S27**). Strikingly, there were 70 ARC/22 VMH sex-DEGs associated with MDD, 68 ARC/16 VMH genes associated with SCZ, and 31 ARC/12 VMH associated with ASD (**Table S12**). MDD-, SCZ-, and ASD-associated genes were also frequent among sex-DEGs of domain-specific clusters, with 25 genes in ARC neurons and 22 in VMH neurons for MDD, 31 ARC/16 VMH for SCZ, and 13 ARC/12 VMH for ASD. While our Xenium data is not a genome-wide assessment, it establishes that genes associated with several sex-biased disorders are also sex-DE in individual cell types of the ARC and/or VMH, including their defining neuronal populations.

Our TF-target analysis also suggested sex-DE regulation by multiple TF-coding genes associated with ASDs and NDDs. Female vVMH and vARC were enriched for targets of ZNF142, which is associated with a rare, female-predominant neurodevelopmental syndrome (*116*) and had divergent target patterns between sexes. Genes upregulated by disruption of *ZNF142* were enriched among female-biased vVMH and vARC genes, while those downregulated by disruption of *ZNF142* were enriched among male-biased vVMH DEGs. As these targets are ascertained from TF perturbation experiments, this would suggest that ZNF142 activity in VMH is *ordinarily* greater in males than in females. On the other hand, male vVMH was enriched for targets of two syndromic ASD genes, *MYT1L* (*103*) and *ASH1L* (*117*) (**Fig. 5B**). In the case of MYT1L, for example, 5/38 of its target genes were nominally (*p*<0.05) sex DE in vVMH, all male-upregulated; similarly, all 14 genes in our ASD risk set that were also targets of MYT1L had a male-up direction of effect (regardless of DE significance). Moreover, *MYT1L* was itself male-upregulated at an FDR<0.1 in xVMH *CRHR2* neurons (**Fig. 4B**, **Fig. 5D**). These results provide further evidence that molecular sex differences in the VMH and ARC are highly relevant for NDDs and ASDs, suggesting a possible mechanism wherein sex-DE of a TF drives sex-DE of several additional disease-associated genes (**Fig. 5F**).

Finally, given that the HYP plays a key role in energy metabolism, we also investigated gene sets relevant for body mass index (BMI) and obesity. GSEA did not reveal overrepresentation of BMI-associated genes in either vARC- or vVMH-enriched genes. However, GSEA using TF-target gene sets showed that both male and female vARC, as well as female vVMH, were enriched in target sets for *STAT3*, a TF necessary for leptin signal transduction. We also found female-specific enrichment of both vARC and vVMH for targets of *PPARA* and *PPARG*, which play key roles in lipid and sterol metabolism (**Fig. 5B**). Interestingly, both of these TFs appear to compete with estrogen receptor α for shared DNA binding sites (*118–120*). In conclusion, these enrichments collectively illustrate that molecular sex differences in tuberal HYP are organized by gene pathways and regulatory hubs critical to HYP functions, gonadal hormone signaling, and disease risk.

## Discussion

The human tuberal HYP contains regions of broad interest in behavioral and endocrinologic neuroscience, but its molecular anatomy has been relatively unexplored, especially in the context of sex differences. Here, we performed comprehensive spatial molecular profiling of two hypothalamic regions, VMH and ARC, in a carefully selected set of healthy male and female control donors enabling comparative transcriptomic analyses in the absence of critical brain or endocrinologic perturbations. Our findings highlight features of HYP that are distinct from more commonly-studied cortical regions, including unexpected *LAMP5* expression in excitatory neurons of the VMH, complex/non-laminar cell type organization in the ARC, and distinctive neuropeptide expression in these domains and their surroundings. Moreover, we identified signatures of sex-regulated gene expression in VMH and ARC, which we localized to specific neuronal populations. Finally, we highlight *ESR1*-expressing neurons in ARC and *CRHR2*-expressing neurons in VMH as hubs for molecular sex differences. These studies provide a roadmap for using transcriptome-wide spatial assays to prioritize genes of interest for finer characterization at cellular resolution using imaging-based spatial transcriptomics.

Given that ARC and VMH are small in size and lack clear anatomical boundaries, generating comprehensive molecular atlases for these regions in the human brain has been challenging to date. Using Visium, we were able to perform an *in silico* dissection of VMH and ARC based on gene expression. While we did not observe discrete spatial clusters within vARC or vVMH, we identified a preponderance of highly spatially variable genes (SVGs) in vARC, which overlapped with established marker genes of postmitotic human fetal ARC neurons (*96*). We hypothesized that higher resolution assays, such as Xenium, would facilitate the identification of spatially-restricted neuronal populations. Indeed, Xenium experiments recapitulated vARC SVG patterns and identified xARC clusters with distinct spatial localization. While there is clear evidence for substructure within the human ARC, we could not detect spatial subdivisions in human VMH. With Visium, VMH clusters were distinguished primarily by extent of glial gene expression, and in Xenium, VMH neuronal clusters were intermingled. We also did not find clear expression of rodent VMH subpopulation marker genes, including those previously reported to be spatially localized, such as *Esr1*-expressing neurons in the posterior ventrolateral rodent VMH.(*3*) Supporting our findings, a single-nucleus RNA-seq study of the developing human HYP through gestational week 20 (a stage at which nearly all neuronal populations have emerged) revealed only one postmitotic VMH cluster.(*96*) There are several possible interpretations for a lack of VMH substructure in our datasets. First, human VMH neuronal cell types may not appreciably exist in spatial niches, or such a phenomenon may only be discernible with very large, cell-resolution datasets and granular clustering. Second, human VMH cell types might be defined by different genes than rodent VMH cell types and for which spatial variability could not be detected within human vVMH. Third, it is possible that molecularly-defined VMH subdivisions exist in a specific anatomical plane across the anterior-posterior axis that was not captured in our study (discussed below).

Nonetheless, marker genes of human vVMH and vARC strongly mapped to the corresponding domains in mice based on integrative studies with a previously published rodent dataset. Several marker genes, especially in ARC, point to conserved physiologic roles implicit in the expression of well-characterized neuropeptides such as *KISS1*, *GNRH*, *AGRP*, and *NPY*. Likewise, domain-enriched genes were enriched in functional terms consistent with rodent studies of VMH (e.g., regulation of blood glucose (*90*, *91*)) and ARC (e.g., primary cilia). Primary cilia are ubiquitous cell structures implicated in the role of ARC in detecting nutritional status. Mice deficient for primary cilium assembly show obesity (*24*); likewise, mutations disrupting primary cilium localization of the melanocortin 4 receptor, which is expressed in ARC, are associated with obesity in humans (*121*).

Given that the HYP is a sexually-dimorphic brain region, we examined sex-DE in ARC and VMH. While a greater number of significant genes and larger fold changes were observed in ARC compared to VMH, we identified biologically relevant sex-DE in both domains. These included genes with HYP-relevant functional roles and regulation: vARC showed greater female expression of *RAMP2* (logFC -1.49, FDR<0.05; **Table S7**), which directly interacts with calcitonin-family receptors and glucagon receptors (*122*), consistent with ARC’s role in metabolic regulation; female vVMH and xVMH both showed greater expression of *TAC1*, a neuropeptide transmitter; and male vVMH demonstrated elevated apelin receptor, *APLNR* (logFC 1.22, *p*<0.01). Interestingly, delivery of apelin into rodent VMH increases lordosis (reproductive behavior) in females (*123*), supporting a sex-differentiated role for this gene. In line with vVMH/vARC marker enrichment in retR-regulated genes, both vARC and xARC demonstrated female upregulation of a retinol dehydrogenase, *RDH10* (**Fig. 2E-F**, **Fig. 3F**) and male upregulation of the retR *RXRG* (**Table S7**). Unsurprisingly, sex-DEGs were also enriched for targets of gonadal hormone receptors. Consistent with rodent literature demonstrating the roles of androgens in sex-differential development of VMH populations (*10*) and in adult HYP sex-DE (*124*), our vARC and vVMH sex-DE signatures were enriched for androgen receptor (AR)-regulated genes, including those induced downstream of protein kinase PKN1 (**Data S5**), providing a possible mechanism for the establishment and/or maintenance of sex-DE. Similarly, GSEA analyses implicated a role for progesterone in observed sex-DE, consistent with observations that progesterone modulates morphology and electrophysiology of female mouse VMH neurons (*125*). Interestingly, *PGR* expression itself was nominally greater in male vVMH, while genes upregulated in female vVMH were enriched for progesterone-responsive genes. This pattern could be due to sex differences in circulating progesterone itself, in activity of PR co-TFs, or both.

Consistent with Visium analysis, we identified robust sex-DE using Xenium, which we resolved to specific neuron subtypes. For example, we pinpointed *ESR1*-expressing neurons as the predominant site of sex-DE within ARC. We also identified sex-DEGs common to several or all domain neuron types, including the retinol dehydrogenase *RDH10*, which was DE in 4/5 xARC neuron clusters. Finally, sex-DE was well-correlated between ARC and VMH, suggesting that sex similarly regulates expression in these two domains. Monosynaptic circuits connecting VMH-ARC have been traced in rodent HYP (*126*); if such circuits exist in human HYP, sex-DE concordance may be partially driven by capture and subsequent quantification of mRNA from neurites of VMH neurons that project to the ARC.

To support translational research efforts, we conducted cross-species analyses using available rodent datasets (*84*, *85*). First, we compared sex hormone receptor expression and distribution between human and mouse HYP. Notably, *Esr1* expression in human VMH and ARC diverged from previous findings in rodents. In mice, *Esr1*-expressing neurons in the VMH, especially in the ventrolateral subdivision, are a well-studied hub of rodent sex differences (*1*, *3*, *8*). By contrast, we observed uniformly sparse *ESR1* expression in human VMH (possibly due to anatomic considerations, discussed below), but dense, high expression in a ventromedial pocket of ARC. We validated this expression pattern using 3 complementary modalities: smFISH, Visium, and Xenium. *ESR1*-expressing cells in human ARC included a subpopulation of *TAC3*+/*KISS1*+ neurons likely representing KNDy neurons previously described in rodent ARC (*5*, *127*). KNDy neurons play a key role in hormone feedback and the luteinizing hormone (LH)/follicle stimulating hormone (FSH) cycle (*9*, *27*, *28*), so it is not surprising that the xARC population containing these neurons would show the most sex-DEGs. At the same time, it is difficult to infer what proportion of our ARC *ESR1*-*TAC3* population comprised probable KNDy neurons as we were not able to include KISS1 on our gene panel. While smFISH indicated it was likely a minority of *ESR1* neurons, it is possible that gonadal hormones may have repressed key transcriptional markers of KNDy cells resulting in lower *KISS1* expression. This speculation is based on mouse studies showing that KNDy marker genes (*Kiss1, Tac2*) are repressed substantially by gonadal hormones (*108*, *128*, *129*), which we assume were circulating in our subjects.

We noted that several results implicated the retinoid pathway in human VMH/ARC biology, including marker, sex DE, and GSEA findings relating to such genes. For example, despite generally shared molecular identities between rodent and human ARC and VMH, retinoid pathway gene *CYP26A1* was a unique marker to human ARC. Additional convergent lines of evidence for retinoid activity in human VMH and ARC included expression of both genes key to local synthesis of active retinoids (*ALDH1A1* and *RDH10*) and enrichment among VMH- or ARC-specifically expressed genes for targets of retRs. Interestingly, however, vARC did not display standout expression of *CRABP1*—a non-nuclear retinoid-binding protein capable of modulating signal transduction via kinase interactions (*130*)—while its mouse ortholog was recently shown to be highly enriched in a POMC-and AGRP-negative, diet-responsive mouse ARC neuron population (*131*). Similarly, divergence between mice and rat in the regulation/expression of genes encoding retinoid synthesis enzymes and RetRs in ARC has been observed (*132*). These findings collectively suggest that retinoid pathway gene expression and regulation is ‘evolutionarily plastic’ in the tuberal HYP, in service of either species-specific (divergent) functions or a shared (conserved or convergent) function via distinct genes.

Last, we explored relationships between HYP gene expression/sex-DE and genetic factors influencing BMI and neuropsychiatric disorders. Surprisingly, we did not find BMI-associated genes to be overrepresented in VMH or ARC, despite the role of HYP in appetite and metabolism. However, our method of gathering genes for this analysis yielded an improbable number of BMI-associated genes (over 8000, *i.e.*, a null expectation that >50% of query genes would be BMI-associated; results using subsets of the ascertained genes for each phenotype are in **Table S11** and **Table S7**). Nonetheless, we did note sex-DE of *CDH13*, a receptor for the insulin-sensitizing hormone adiponectin, in ARC, and sex-DE of the appetite-regulating and obesity-associated (*102*) melanocortin receptor *MC4R* in VMH. The most noteworthy finding, supported by multiple analyses, was the robust enrichment of male-upregulated genes of both vVMH and vARC for ASD and NDD risk gene sets. Moreover, male-upregulated genes were overrepresented in targets of TFs encoded by genes mutated in syndromic ASDs/NDDs, including *ZNF142* (*116*), *TCF20* (*133*), *ASH1L* (*117*), and *MYT1L*—a syndrome which includes obesity (*103*, *134*). An interesting single-gene link to MDD, which occurs more often in females, also emerged: the sex-DEG *CDH13* is repressed by a human-specific lncRNA *FEDORA*, which has been shown to be sex-DE in MDD cortex and to exert sex-differential behavioral effects when expressed in mouse cortex (*135–137*). Moreover, *MYT1L* was itself significantly male-upregulated in *TAC3*-*ESR1* KNDy neurons of the ARC and *CRHR2* neurons in VMH. These findings highlight VMH and ARC as candidate HYP regions susceptible (perhaps sex-differentially) to common- and rare-variant mediated neurodevelopmental disorder risk.

We note several limitations in the present study. First, consistent with previous efforts to map the human hypothalamus (*138*), we only sampled one position along the AP axis per donor. We note that domain shape and size varied across donors, which we attribute to variability in cutting angle during brain slabbing and tissue block sectioning, and potentially to anterior-posterior (AP) variation in sampling. Based on our findings, we estimate that there is only a modest degree of AP sampling variation: we screened tissue for VMH content and study inclusion using smFISH for *NR5A1*, which is only expressed in anterior/middle VMH of adult rodents (*3*). Meanwhile, *Esr1*+ neurons are most abundant in the posterior mouse VMHvl (*3*) consistent with anterior *NR5A1* expression and limited VMH *ESR1* expression throughout our dataset. Finally, despite the known presence (*139*, *140*) and important function in lactation (*141*) of dopamine neurons in ARC, this population was not identified or characterized in our data due to insufficient dopaminergic marker expression. Collectively, our design and results suggest that we sampled across the anterior-to-middle portion VMH, specifically where ARC is also present. We further consider it unlikely that systematic AP variation impacted our sex-DE findings as tissue blocks across our cohort were sampled by cutting A-to-P in some cases and P-to-A in others; if AP variation in gene expression affected sex-DEGs, then we might expect poor agreement between Visium and Xenium with regards to sex-DE (contrary to our observations). Furthermore, while heterogeneous gene expression *within* cell types along the AP axis has been observed in other areas of rodent HYP, this phenomenon only involved a few genes (*142*). Recent molecular atlasing of the human HYP identified several VMH clusters from snRNA-seq, all of which were represented in a single AP plane (*44*) indicates that AP sampling variation should not affect our ability to detect (non-*ESR1*) VMH populations and is consistent with identification of cells from all xVMH/xARC clusters in all Xenium samples. Focusing our Xenium sex-DE analyses on cell clusters or ‘types’ found across the entire sample set, we found sex differences far more widespread than could be supported by evidence of intra-regional AP expression gradients alone.

Second, there are methodological limitations to both spatial transcriptomic approaches employed. While Visium is transcriptome-wide, it is lower resolution and prohibits granular spatial-molecular organization of small tissue areas such as VMH and ARC. We addressed this limitation by employing Xenium, which resolves single nuclei in space, but at the expense of measuring only a fraction of the transcriptome. Our Xenium data cover only 366 genes, potentially preventing identification of cell types, including tanycytes and cells of VMH. In the former case, tanycyte labels were assigned to rare cells in the median eminence bordering VMH/ARC as would be expected (*143*), but also to cells intermingled with connective and vascular clusters in many samples (**Fig. S30**), illustrating the limitations of cell-resolution spatial transcriptomics as a method for cell type discovery. Separately, findings regarding *ESR1*, KNDy neurons, and sex-DE could be potentially affected by menstrual stage and/or oral contraceptive use by female donors, though this information was neither systematically collected nor necessarily accurate (only available when reported by next of kin). For example, while we find *MC4R* to be male-upregulated in the VMH of our cohort, it has been previously reported female-upregulated in mice in an estrus stage-dependent manner (*94*). At the same time, findings on *ESR1* distribution were consistent between sexes, and our smFISH experiment for *ESR1* and *KISS1* was performed on tissue from a male donor. While other recent human atlasing efforts have made headway in profiling a broader extent of HYP (*43*, *44*), they have not been powered to address sex differences in any specific area. Future studies should build upon this and other foundational molecular work in human HYP by profiling its domains in-depth—that is, at similar or greater sampling depths to ours here—in order to power studies contrasting the effects of sex or physiology (BMI, reproductive status, etc.) on HYP.

In sum, these data represent a unique transcriptomic resource focused on the human tuberal HYP, an area important for endocrine, reproductive, and behavioral functions. We provide a spatially-resolved single cell atlas of the adult human HYP and demonstrate that the molecular identities of VMH and ARC are generally similar between mice and humans. Further, we show that autosomal gene expression differs across the sexes in both VMH and ARC, often in a cell type-specific manner. Finally, we show that sex differences in gene expression are associated with ASDs and NDDs, disorders with greater prevalence in males. To augment ongoing HYP research, we generated interactive web-based data browsers for the scientific community to explore these data at https://bit.ly/libdVisHYP (Visium) and https://bit.ly/libdXenHYP (Xenium).

## Supporting information

Figures S1-S30 and Legends for All Supplemental Materials

Table S1-Demographics and Sequencing Quality Metrics for Visium

Table S2-SVG Analysis of Visium Sections

Table S3-One-vs-All Marker Analyses for all Visium and Xenium Clusters

Table S4-One-vs-Other Marker Analyses Among vVMH or vARC Clusters

Table S5-Marker Gene Registration Between Humans and Mice

Table S6-Gene Sets Collated from Outside Sources for GSEA

Table S7-Visium Sex DE and Disease-GSEA Analyses

Table S8-Xenium Gene Panel Prioritization and Notes

Table S9-Intra-vVMH and Intra-vARC SVG Analyses

Table S10-Xenium Sex DE Results

Table S11-GSEA Analysis of Visium Marker Genes

Table S12-Disease Genes, TFs, and Sex-DE

## Acknowledgments

We thank the Joint High Performance Computing Exchange (JHPCE) at Johns Hopkins University for providing computing resources for these analyses. We thank the JHU Single Cell Transcriptomics core for Illumina sequencing and Xenium data generation. We thank the families of Connie and Stephen Lieber and Milton and Tamar Maltz for their generous support of this work. We thank the LIBD neuropathology team, particularly James Tooke and Amy Deep-Soboslay, for curation of the brain samples and assistance with tissue dissections, and the families of the brain donors for their generosity. We also would like to thank William Reay, PhD, for insightful discussion regarding retinoid genes/pathways in the brain and disease. We thank Madhavi Tippani for reviewing and testing code for smFISH puncta quantification and visualization in R. Schematic illustrations were generated using Biorender and Adobe Illustrator.

## Author contributions

Conceptualization: BM, KM, KDH, KRM

Methodology: AC, BM, KM, KDH, KRM, SCH, SCP, YW

Software: AC, BM, KDH, RAM, SCH, YW

Validation: BM, HRD, KDH, KDM, SVB, YD

Formal Analysis: AC, BM, HRD, KDH, KRM, RAM, YW

Investigation: AC, HRD, KDM, SC, SCP, SVB, TMH, YD

Resources: JEK, TMH

Data Curation: BM, KDH, RAM

Writing-original draft: AC, BM, HRD, KM, KDH, KDM, KRM, SC, SVB

Writing-review and editing: BM, KM, KDH, KRM

Visualization: AC, BM, HRD, KDH, KRM, RAM, SC, SVB, YD

Supervision: KM, KDH, KRM, SCH

Project administration: KM, KDH, KRM

Funding Acquisition: KM, KDH, KRM

## Data and materials availability

### From this study

The source data described in this manuscript, including FASTQ files from Visium and unfiltered sample-level xeniumranger 1.7 outputs (transcript-level data) compatible with Xenium Explorer software are respectively available through GEO accessions **GSE280316** and **GSE280460**. Supplemental Data files, smFISH microscopy images, the Visium SpatialExperiment (including bundled H&E images) and Xenium SpatialFeatureExperiment objects as filtered, clustered, and analyzed, and Xenium hematoxylin and eosin images (not used in the study) are available through http://research.libd.org/globus/ via the Globus endpoint **jhpce#HYP_suppdata**. Source code for all processing/analyses and plotting is available on GitHub (https://github.com/LieberInstitute/spatial_HYP), with contents at the time of submission archived to Zenodo (https://zenodo.org/records/14285059). Data can be interactively visualized and annotated in a web browser through our Samui (*144*) portals for the Visium data (with clustering available at four different resolutions from BayesSpace) at bit.ly/libdVisHYP (permanent URL: https://research.libd.org/spatial_HYP) and for Xenium (with annotated cell clusters from Banksy, their higher-order cell groups, and domains) at https://bit.ly/libdXenHYP (permanent URL: https://research.libd.org/spatial_HYP). Documentation for the data portals can be accessed through https://research.libd.org/spatial_HYP.

### Outside data/databases

Raw h5ad files with single-cell RNA-seq data from adult mouse HYP (*85*) were downloaded from https://allen-brain-cell-atlas.s3-us-west-2.amazonaws.com/expression_matrices/WMB-10Xv3/20230630/WMB-10Xv3-HPF-raw.h5ad and https://allen-brain-cell-atlas.s3-us-west-2.amazonaws.com/expression_matrices/WMB-10Xv2/20230630/WMB-10Xv2-HY-raw.h5ad, then were filtered to cells that received cluster labels found in https://allen-brain-cell-atlas.s3-us-west-2.amazonaws.com/metadata/WMB-10X/20230630/views/cell_metadata_with_cluster_annotation.csv. TF-target gene sets (Transcription_Factor_PPIs, ChEA_2022, Rummagene_transcription_factors, TF_Perturbations_Followed_by_Expression, TF-LOF_Expression_from_GEO, TRRUST_Transcription_Factors_2019) were obtained from the gene set libraries used by the Enrichr tool (*75*) at https://maayanlab.cloud/Enrichr/#libraries on 12/06/2023. Our assertion for low expression of transcription factor RNAs is based on an interactive plot from (*76*) illustrating the transcripts per million for each likely human TF in each tissue in data from the Human Tissue Atlas: http://humantfs.ccbr.utoronto.ca/.

### Non-Bioconductor/CRAN R Packages

scCoco and the Allen Brain Atlas API interface it utilizes, cocoframer, are respectively available at https://github.com/lsteuernagel/scCoco and https://github.com/AllenInstitute/cocoframer.

