## Supplementary material for "Spatially-resolved molecular sex differences at single cell resolution in the adult human ventromedial and arcuate hypothalamus": Figures S1-S30 and Legends for All Supplemental Materials

### Supplementary Figures S1-S30, Legends for Data S1-S5, and Legends for Supplementary Tables S1-S12 from

***Spatially-resolved molecular sex differences at single cell resolution in the adult human ventromedial and arcuate hypothalamus***

Bernard Mulvey *et al.*

#### Supplementary Tables

**Table S1. Experimental metadata (separate file).**

**Sheet A**: Sample processing information and sequencing metrics (by sample, with rotations and reflections columns included for how spatial data was manipulated)
**Sheet B**: Sample demographics.

**Table S2. Highly-variable and spatially variable genes for Visium (separate file).**

**Sheet A**-Table of gene name and whether it was a gene in the HVG 10%ile / HVG 20%ile / nnSVG 10%ile / nnSVG 20%ile set.
**Sheet B**-Full output of *nnSVG* analyses per gene per sample (including features not in either feature set from sheet A).
**Sheet C**-*nnSVG* per-sample ranks and mean ranks (taken across nominally significant samples) for all genes achieving nominal significance in at least one sample.

**Table S3. Marker analyses of Visium and Xenium clusterings (separate file).**

**Sheet A**: Inferred predominant cell population for each Visium *k*=15 cluster.
**Sheets B-E**: *spatialLIBD* one-vs-all ("enrichment") outputs for *k* = 15 (collapsed VMH/ARC), 15, 20, 31, respectively. VMH and ARC clusters are not collapsed in the latter three sheets.
**Sheet F:** Nomenclature reference for Xenium analyses. Cluster assignments from Banksy (X1, X2…X41), their generic annotations ('ARC 1', 'ARC 2', etc.), and for ARC/VMH neuron clusters, the descriptive annotations as used in illustrations ('ARC-AGRP', etc.). Donor/sample-specific clusters that were excluded in downstream analyses are not listed.
**Sheet G**: *spatialLIBD* one-vs-all ("enrichment") outputs across all 33 retained Xenium clusters.
**Sheet H**: *spatialLIBD* one-vs-all ("enrichment") outputs when only considering cells within the xARC domain and labeled as one of the 5 xARC clusters.
**Sheet I**: *spatialLIBD* one-vs-all ("enrichment") outputs when only considering cells within the xVMH domain and labeled as one of the 4 xVMH clusters.
**Sheet J**: *spatialLIBD* one-vs-all ("enrichment") outputs when only considering **nuclear counts** for cells within the xARC domain and labeled as one of the 5 xARC clusters.
**Sheet K**: *spatialLIBD* one-vs-all ("enrichment") outputs when only considering **nuclear counts** for cells within the xVMH domain and labeled as one of the 4 xVMH clusters.

**Table S4. Pairwise marker analyses between k=15, 20, or 31 VMH clusters and between k=15, 20, or 31 ARC clusters (separate file).** *scran* pairwise marker analyses between individual VMH or ARC clusters for *k*=15, 20, and 31. Sheet name denotes the BayesSpace cluster being tested for marker status and comparator cluster. (VMH1/VMH2 and ARC1/ARC2 are the VMH/ARC clusters from k=15 clustering). Note that these metrics can be interpreted bidirectionally as each comparison is between pairs of clusters; that is, for example, that a gene highly enriched in the tested cluster relative to the comparator is necessarily highly depleted in the comparator cluster compared to the tested cluster.

**Table S5. Visium marker gene spatial registration results in mouse: *scCoco* results for all mouse hypothalamic areas mapped from Allen ISH atlas; registration to mouse HYP subclasses from single-cell RNA-seq (*85*) (separate file).**

**Sheets A-D**: scCoco domain-to-mouse brain atlas HYP region scores using 50-150 top human markers per domain from the k=15 (collapsed VMH, ARC), k=15 using individual domains, k=20 domains, k=31 domains respectively.

**Sheet E:** Spearman correlations between (i.e., spatial registration of) mouse HYP cell "subclasses" and each Visium cluster. Mouse VMH and ARC ("ARH") subclasses were aggregated into single domains for comparability to Visium domains. Results are included for Visium clusterings at k=15 (with collapsed VMH, ARC clusters), k=15 using individual clusters, k=20, or k=31. Correlations were tested using variable numbers of top *n* marker genes (ranked by *t*-statistic) per human cluster. Only genes with 1-1 mouse orthologs and present in both datasets were used.

**Table S6. Aggregated disease-gene sets and TF-target gene sets (separate file).**

**Sheet A**: Source list delineating GWASes and databases from whence genes for each diagnosis or trait were collected.

**Sheets B-E**: GWAS, TWAS, disgenet, and union gene sets per diagnosis from above.

**Sheet F**: FUMA gene mapping results for PGC Panic disorder (run for preparing these gene sets).

**Sheet G**: FUMA parameters for PGC Panic disorder gene mapping.

**Sheet H:** Aggregated TF-target pairs from all databases queried.

**Sheet I**: Curated primary cilia gene sets obtained from (*93*).

**Table S7. Visium pseudobulk sex-DE results for k=15, k=15 (collapsed), k=20, k=31, and domain sex-DE pattern enrichments in disease-gene sets (separate file).**

**Sheets A-D:** Pseudobulked sex-DE results for Visium cluster assignments using each k-value, including single k=15 domains and the collapsed vVMH/vARC domains as primarily reported in the main text.

**Sheet E:** Collated results from all Visium sex-DE GSEA analyses for BMI and neuropsychiatric disorders. All BayesSpace classifications (k=15 (collapsed VMH/ARC), k=15, k=20, and k=31) and disease-gene sets/subsets (GWAS publication genes only, TWAS genes only, DisGeNet ("dgn") genes only, union of genes from these sources) are presented in a single table.

**Table S8. Genes included on Xenium assay (separate file).** For custom genes in the assay, notes taken for gene prioritization (based on examinations of preliminary Visium analyses) are included.

**Table S9. Results of *nnSVG* analysis performed only on Visium spots classified as ARC or only classified as VMH (separate file).**

**Sheet A**-Full output of *nnSVG* analyses of ARC, per gene per sample.
**Sheet B**-Full output of *nnSVG* analyses of VMH, per gene per sample.
**Sheet C**-ARC-only *nnSVG* per-sample ranks and mean ranks (taken across nominally significant samples) for all genes achieving nominal significance in at least one sample in this analysis.
**Sheet D**-VMH-only *nnSVG* per-sample ranks and mean ranks (taken across nominally significant samples) for all genes achieving nominal significance in at least one sample in this analysis.

**Table S10. Xenium pseudobulk sex-DE results for each cluster, considering only cells within boundaries of xARC or of xVMH; sex-DE when pseudobulking all cells in the xARC or xVMH domain; confirmatory Xenium sex-DE analyses (separate file).** Positive logFC indicates male expression greater than female.

**Sheet A:** Pseudobulk sex-DE results by Xenium cluster only considering cells within xARC.
**Sheet B:** Pseudobulk sex-DE results by Xenium cluster only considering cells within xVMH.
**Sheet C**: Pseudobulk sex-DE results at the level of Xenium domains (all cells within xARC or all cells within xVMH).
**Sheet D**: Pseudobulk sex-DE results by Xenium cluster, only considering transcripts within segmented nuclei in xARC.
**Sheet E**: Pseudobulk sex-DE results by Xenium cluster, only considering transcripts within segmented nuclei in xVMH.
**Sheet F**: Pseudobulk sex-DE results by Xenium cluster, only considering cells within a shrunken xARC domain (so as to exclude the domain boundary, where xVMH-xARC populations would likely comingle).
**Sheet G**: Pseudobulk sex-DE results by Xenium cluster only considering cells within a shrunken xVMH domain (so as to exclude the domain boundary, where xVMH-xARC populations would most likely comingle).
**Sheet H**: Correlation of sex-DE *t*-statistics between all pairs of Xenium xARC/xVMH cluster analyses (primary domain-level analysis, primary clusterwise-within-a-domain analysis, nuclear-counts, shrunken domain counts) amongst themselves and Visium domain or *k*=15 VMH/ARC clusters. The analysis and cluster of the first comparator are listed in "*Cluster/Domain 1 Analysis*" and "*Cluster/Domain 1*", respectively. Individual Xenium clusters were only compared to themselves and to Visium clusters/domains.

**Table S11. GSEA results: Visium domain-enriched gene overlap with disease/trait-associated genes (separate file).** Collated results for all domains from all BayesSpace classifications for source data-subsetted disease/trait-gene sets (**A**) union gene set of all sources, (**B**) GWAS publication genes only, (**C**) TWAS genes only, (**D**) DisGeNet genes only for each diagnosis/trait using BayesSpace k=15 (collapsed VMH/ARC), k=15, k=20, and k=31 clusterings.

**Table S12. Each gene assayed on Visium and/or Xenium, whether it is a TF tested in the TF-target GSEA analyses or in 1 or more disease-gene sets, and information about its expression and sex-DE from Visium and Xenium (separate file)**. Each gene measured in Visium and/or Xenium is listed. Columns indicate whether the gene was assayed with Xenium; which of the disease(s) gene lists the gene appears in; whether the gene is expressed in vVMH/vARC overall and whether it is expressed in each Visium domain on a single-sex basis; whether the gene is nominally sex-DE or significantly sex-DE in each Visium domain; each Xenium cluster within xARC wherein the gene was sex-DE, if any; and each cluster within xVMH wherein the gene was sex-DE, if any.

#### Supplementary Data File Legends

**Data S1. Visium PCA and UMAP, before and after HARMONY dimensionality reduction (separate file).** Plots depicting PCs 1 vs 2 and 1 vs 3 or UMAP with dimensionality reductions on the 10%ile nnSVG feature set, without and with HARMONY batch correction. Plot legends indicate the variable by which points (Visium spots) have been colored, including BayesSpace cluster assignments (ascertained using HARMONY dimensionality reduction on the 10%ile SVG feature set), demographic characteristics, and technical variables.

**Data S2. GSEA results for Visium domain/cluster-enriched genes in MSigDb (separate file).** File is provided in an R data format (*.RDS*). Composed of an R list with names corresponding to which domain assignments were tested. Each entry of the list holds a dataframe of collated, unabridged results produced by *fgsea::fgseaMultilevel*, with a column added to mark whether or not that row was one of the 'collapsed' results following independent pathway analysis with *fgsea::collapsePathways(). fgsea* output tables for each Visium cluster in a BayesSpace run were collapsed together into one table, with a column added to indicate the analyzed Visium cluster. For a cluster-by-BayesSpace run-by-gene set combination, the gene set name was checked against the corresponding output from *collapsePathways()* for labeling as in the 'collapsed' gene set results. Additional information on columns output by *fgseaMultilevel* can be found in the package's documentation.

**Data S3. GSEA results for Visium domain/cluster-enriched genes in aggregated TF-target gene sets (separate file).** File is provided in an R data format (*.RDS*). Contains a list, named according to the BayesSpace *k* value and cluster tested (*k*=15, 20, 31, or *k*=15 with ARC and VMH collapsed into singular domains). Each entry of the list holds a dataframe of collated results produced by *fgsea::fgseaMultilevel*, with a column added to mark whether or not that row was one of the 'collapsed' results following independent pathway analysis with *fgsea::collapsePathways().* Only gene sets with a positive enrichment score (*i.e.*, overrepresented in the tested Visium genes) are presented. Each table was collated as described for File S2.

**Data S4. GSEA results for Visium sex-DE pattern enrichment in TF-target gene sets (separate file).** File is provided in an R data format (*.RDS*). Contains a list named as described for file S5. Each entry of the list holds a dataframe of collated, unabridged results produced by *fgsea::fgseaMultilevel*, with columns added to indicate whether the TF qualified as expressed in male and/or female samples (at least three samples from a given sex with the TF detected in ≥1% of spots in a cluster, and the TF detected in at least 2.5% of spots overall for that cluster and sex). For signed test results, the DE effect direction explicitly tested (enrichment in male-upregulated genes or in female-upregulated genes) is indicated, which corresponds to a positive or negative value, respectively, for enrichment score and normalized enrichment score.

**Data S5. GSEA results for Visium sex-DE pattern enrichment in MSigDb (separate file).** File is provided in an R data format (*.RDS*). Contains a list named as described for file S2, but with two result tables produced by *fgsea::fgseaMultilevel* for every cluster tested: one "signed", indicating a GSEA run including direction of DE to identify enrichments among male-upregulated or female-upregulated genes, and one "unsigned", indicating a GSEA run with the absolute values of DE statistics to identify terms enriched for sex effects bidirectionally. For signed test results, the DE effect direction explicitly tested (enrichment in male-upregulated genes or in female-upregulated genes) is indicated, which corresponds to a positive or negative value, respectively, for enrichment score and normalized enrichment score.

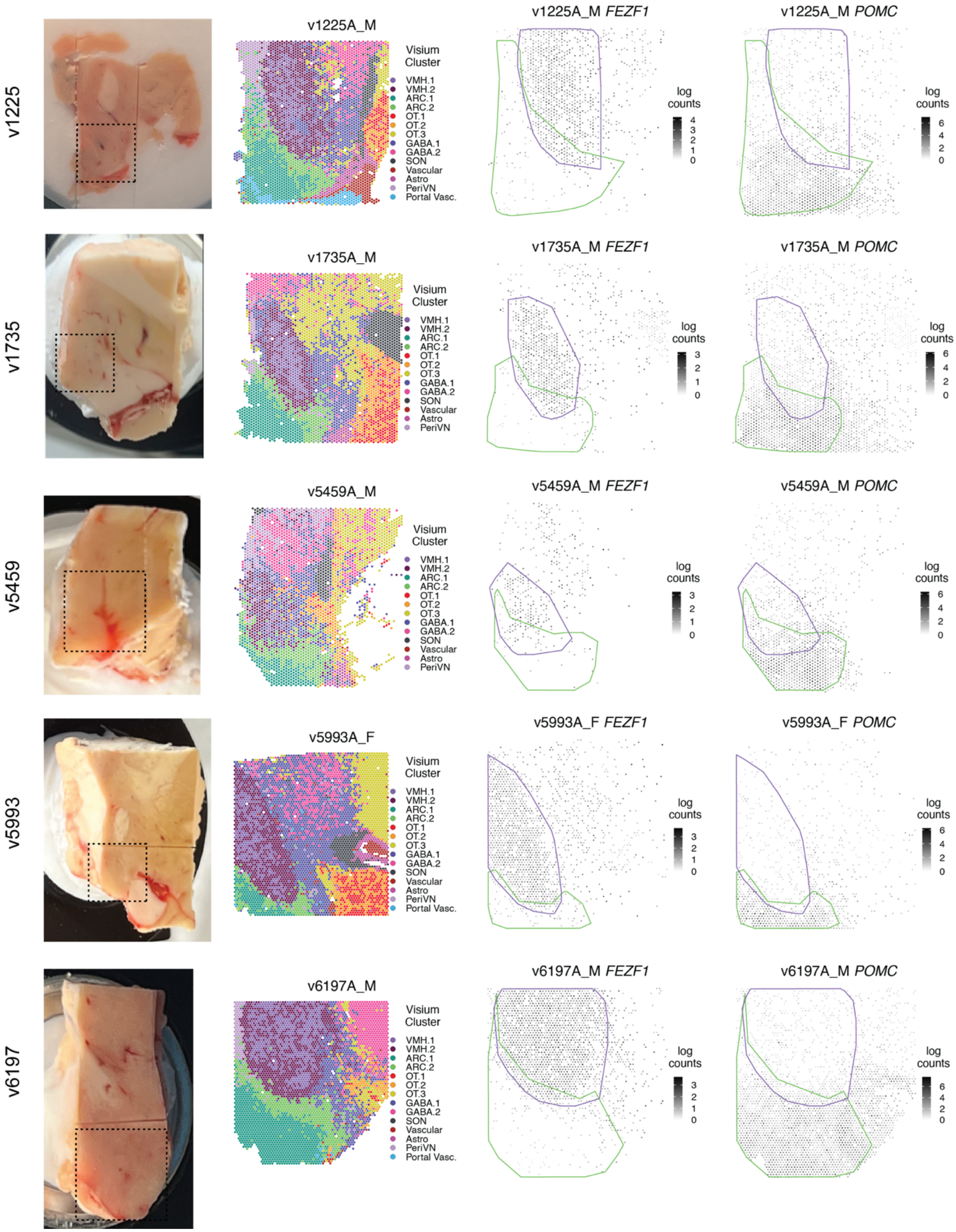


**Figure S1. Identification of VMH and ARC boundaries in Visium samples. (Continues on next page.)**

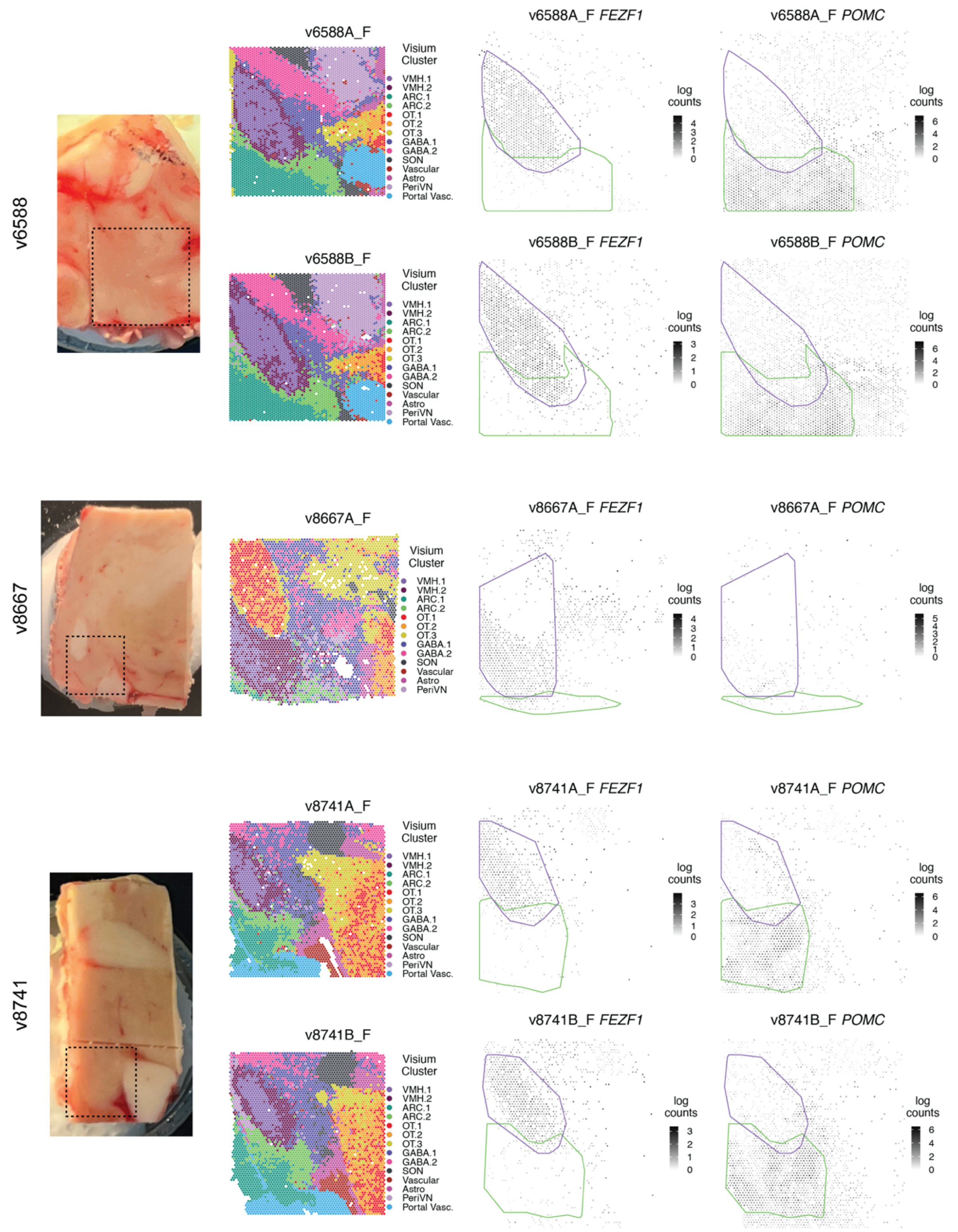


**Figure S1. Identification of VMH and ARC boundaries in Visium samples.** Left: Representative image of HYP tissue blocks containing VMH and ARC. Boxes denote the approximate portion of tissue used for Visium/Xenium assays. Middle: *BayesSpace* clustering at k=15 identifying 2 VMH and 2 ARC clusters. Right: Spotplots showing log transformed normalized expression (logcounts) of *FEZF1* (VMH marker) and *POMC* (ARC marker) per donor and sample(s). Approximate Visium (v)VMH and Visium (v)ARC domain boundaries are illustrated for reference—note that these boundaries, in the case of Visium, are for illustrative purposes only and were *not* used for any analyses.

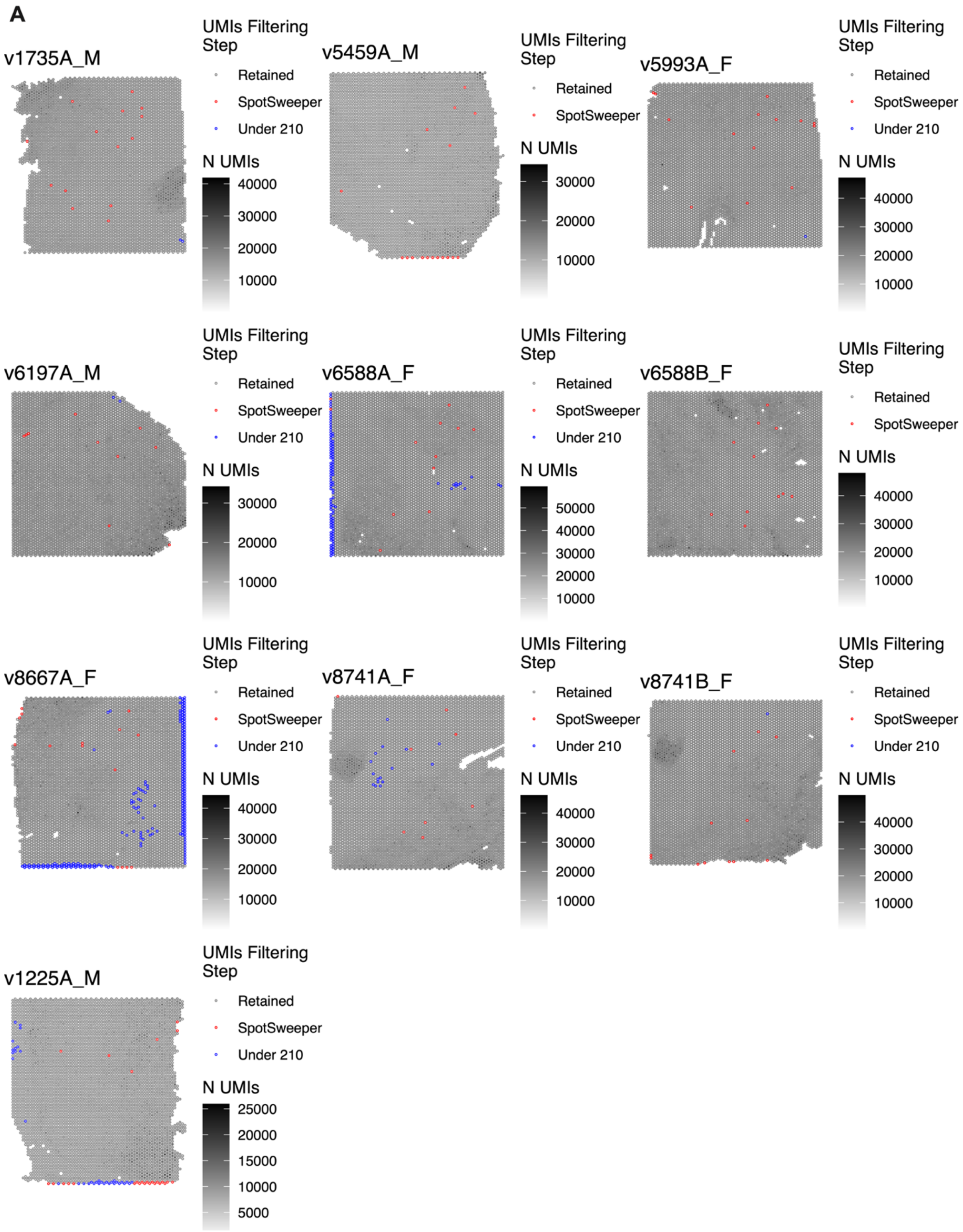


**Figure S2. Spots removed from each Visium sample by local outlier quality control with *SpotSweeper* followed by manual filtering.** **(*Continued on next page*)**

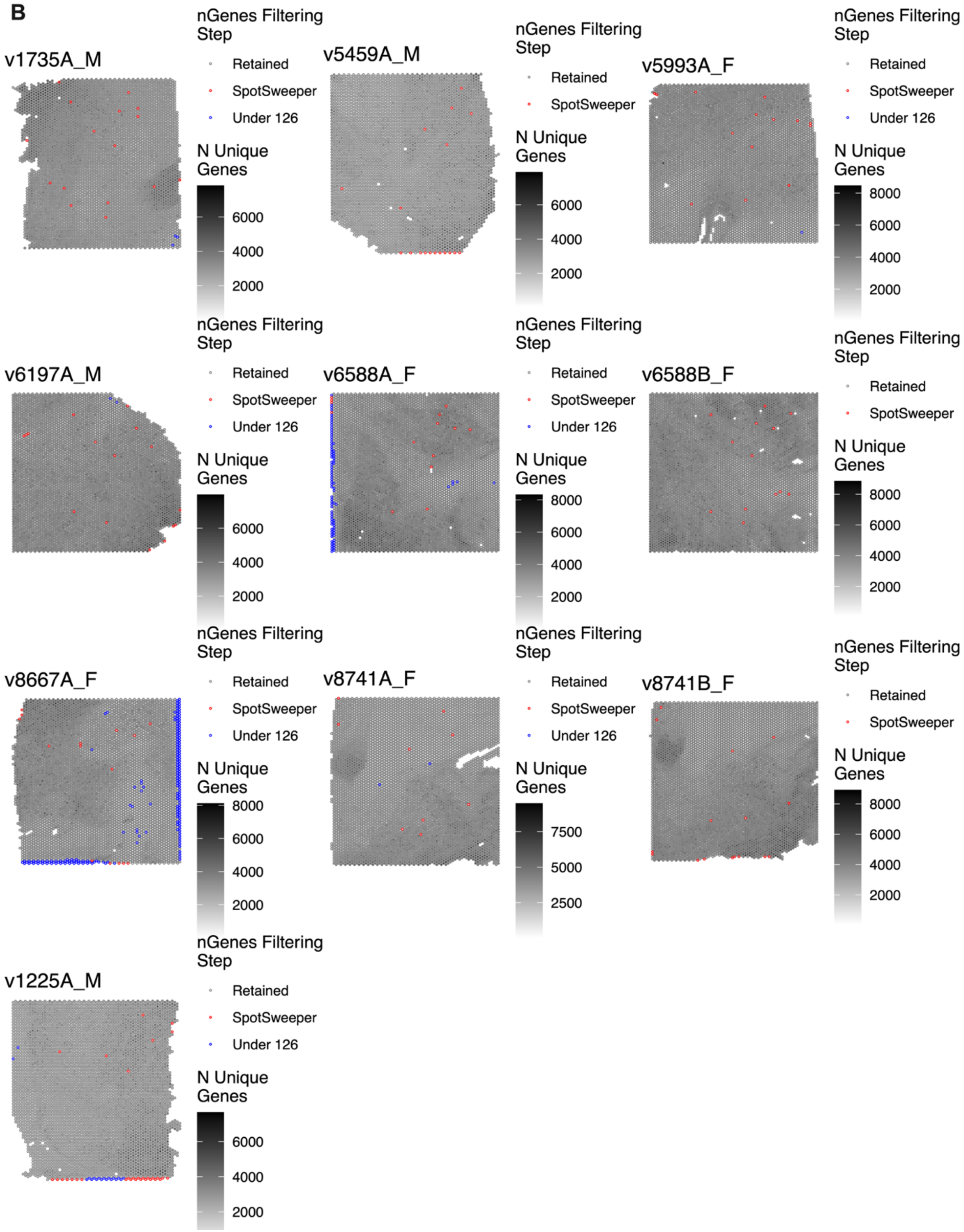


**Figure S2. Spots removed from each Visium sample by local outlier quality control with *SpotSweeper* followed by manual filtering.** **(*Continued on next page*)**

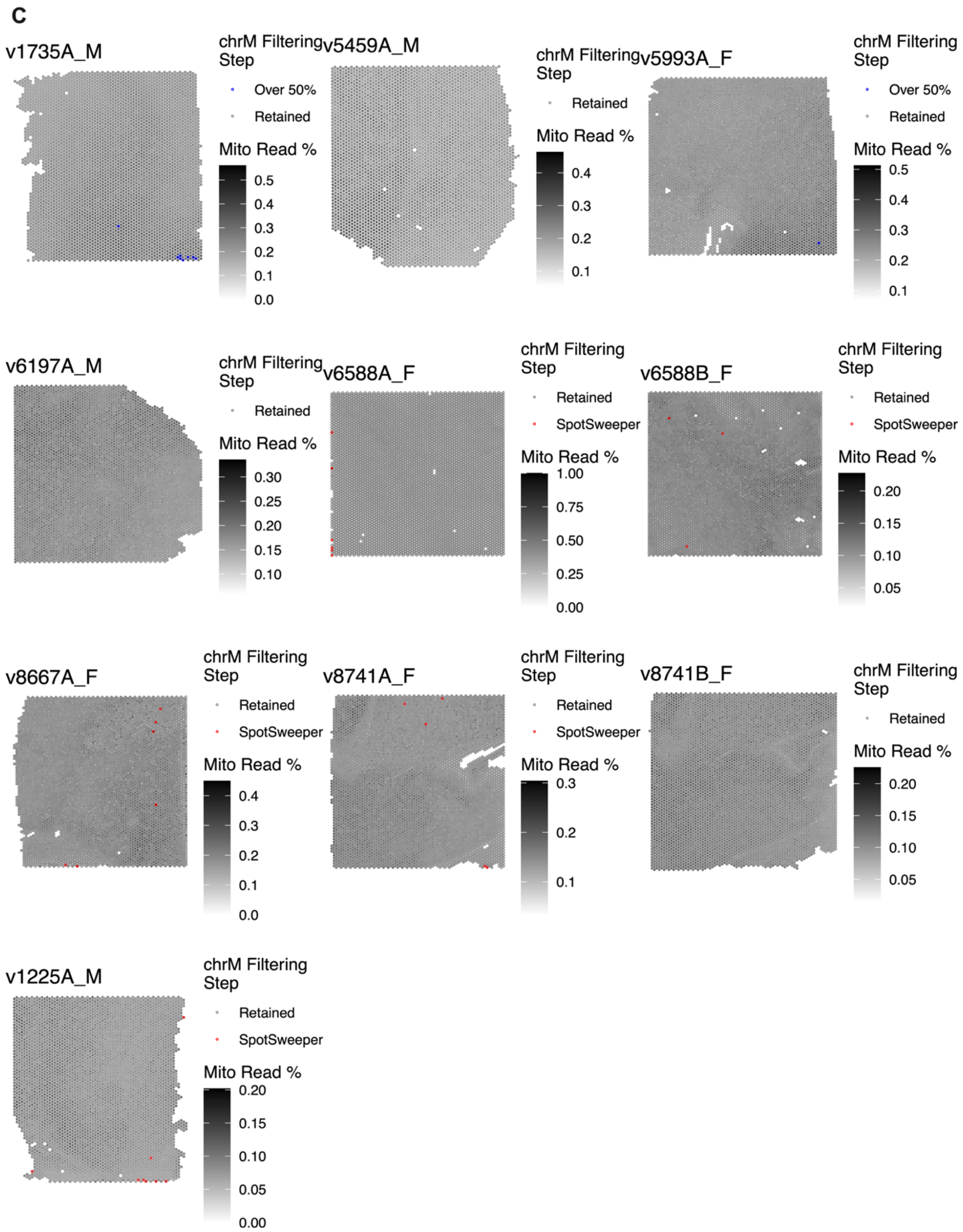


**Figure S2. Spots removed from each Visium sample by local outlier quality control with *SpotSweeper* followed by manual filtering. A)** Filtering of local outliers for UMIs per spot and subsequently for low UMI content (under 210 UMIs/spot). **B)** Filtering of local outliers for number of unique genes per spot and subsequently for low number of unique genes/spot (under 126 genes/spot). **C)** Filtering for proportion of mitochondrial reads on the basis of local outliers, and subsequently for spots exceeding a numerical threshold (50%).

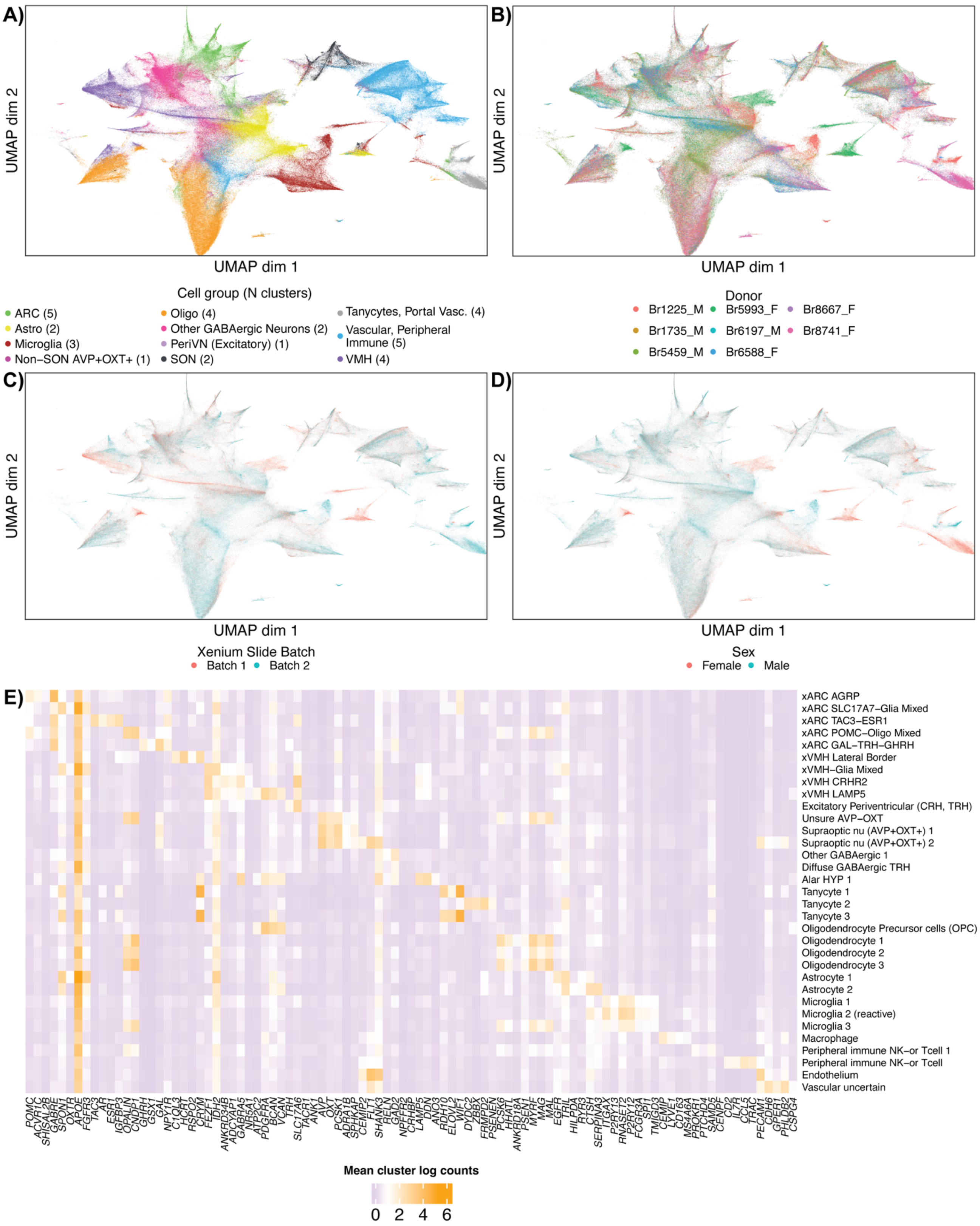


**Fig S3. UMAP of Xenium cell data annotated with major cell type clusters, sex, donor, or experimental batch; heatmap of top markers for all retained Xenium clusters. A)** UMAP illustrating Xenium cell type clusters grouped by the overarching cell type (e.g., astrocyte) and/or inferred domain of origin (e.g., periventricular nucleus). **B)** Same UMAP dimensions as in A, colored by brain donor. **C**) Same UMAP dimensions as in A, colored by experimental batch. **D**) Same UMAP dimensions as in A, colored by donor sex. **E**) Heatmap displaying average log counts top cluster marker genes for each of the 33 Xenium clusters retained in downstream analyses. All UMAPs illustrate dimensionality reduction following normalization of non-log adjusted count data, which was used as input for Banksy clustering (as its authors suggest[87]).

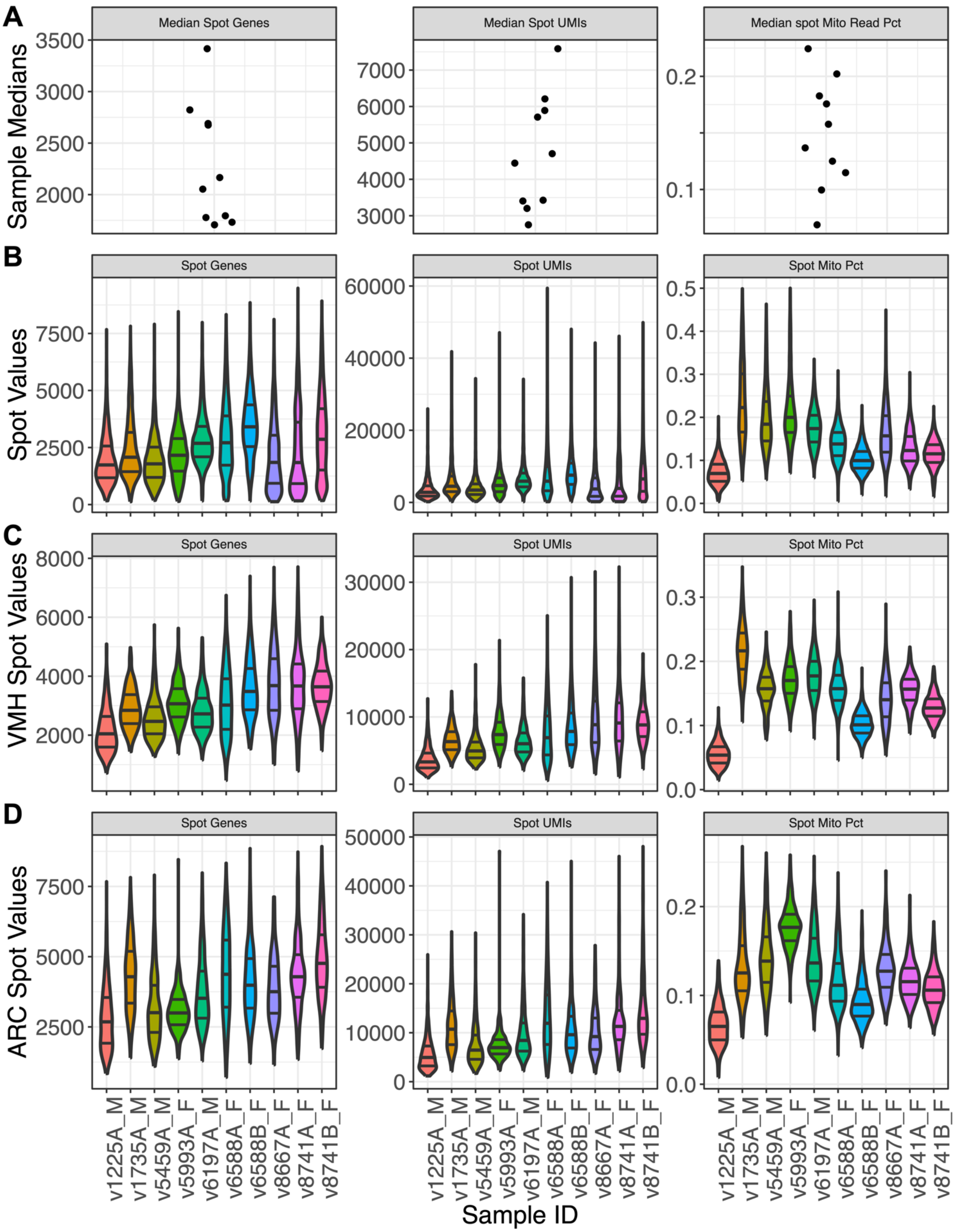


**Fig S4. Quality control metrics for the final Visium dataset.** Number of unique genes per spot, UMIs per spot, and % mitochondrial reads per spot are shown in columns. **A)** Samplewise medians for each metric. **B)** Spot-level values per sample. **C)** Spot values per sample only among spots labeled xVMH. **D)** Spot values per sample only among spots labeled ARC. Each violin in B-D shows a dotted line to demarcate the sample median, and solid lines to demarcate 25th and 75th %iles. All plots reflect the filtered dataset as used in downstream analyses.

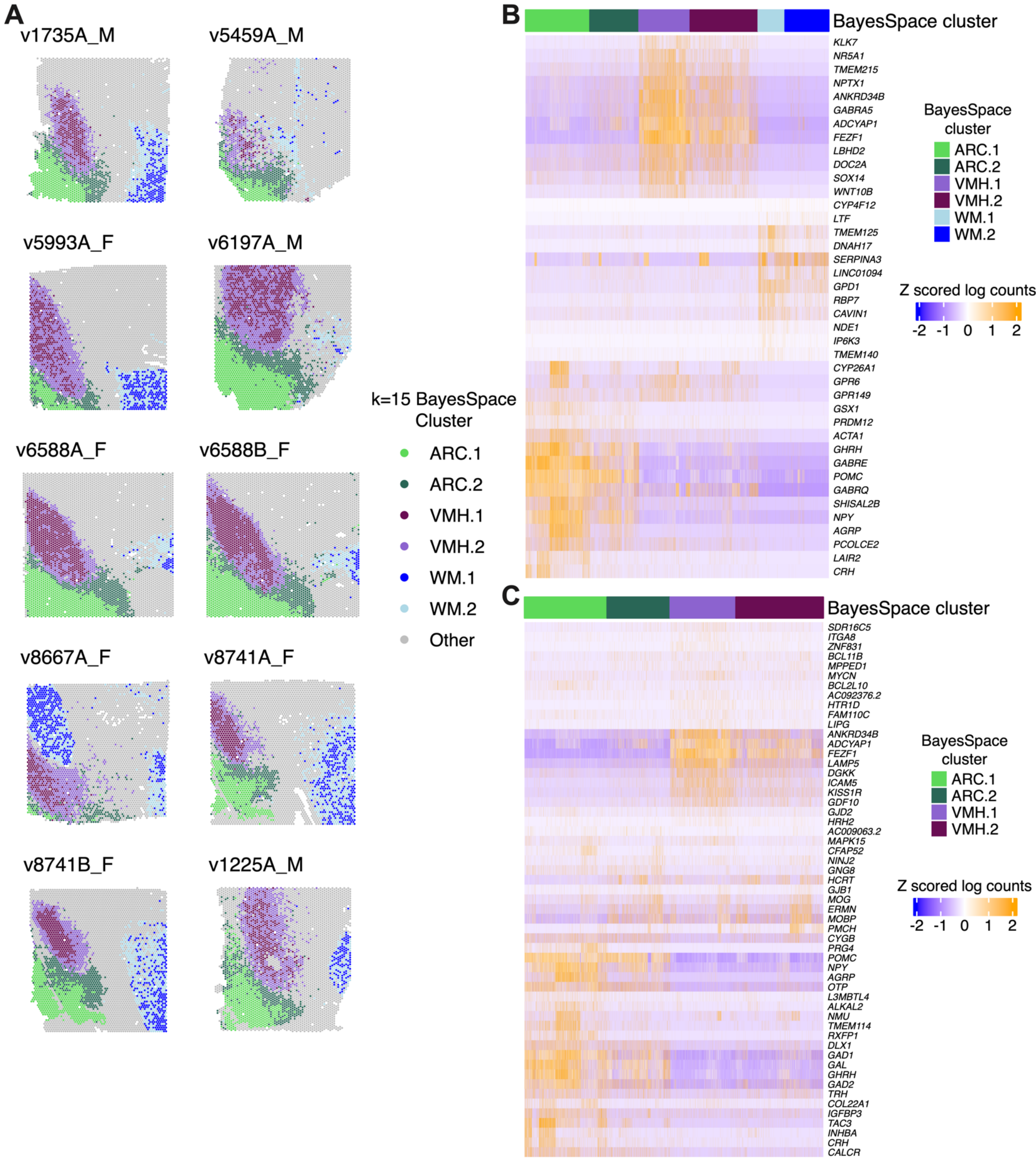


**Figure S5. Characterization of VMH and ARC clusters and identification of marker genes with BayesSpace at *k*=15**. **A)** Spotplots of individual VMH, ARC, and optic tract (OT) white matter (WM) clusters at *k*=15 for all Visium samples. **B)** Heatmap of top marker genes for VMH,

ARC, and WM (for comparison) clusters using one vs. all marker gene analyses. **C)** Heatmap of top marker genes using pairwise comparisons between VMH clusters and ARC clusters.

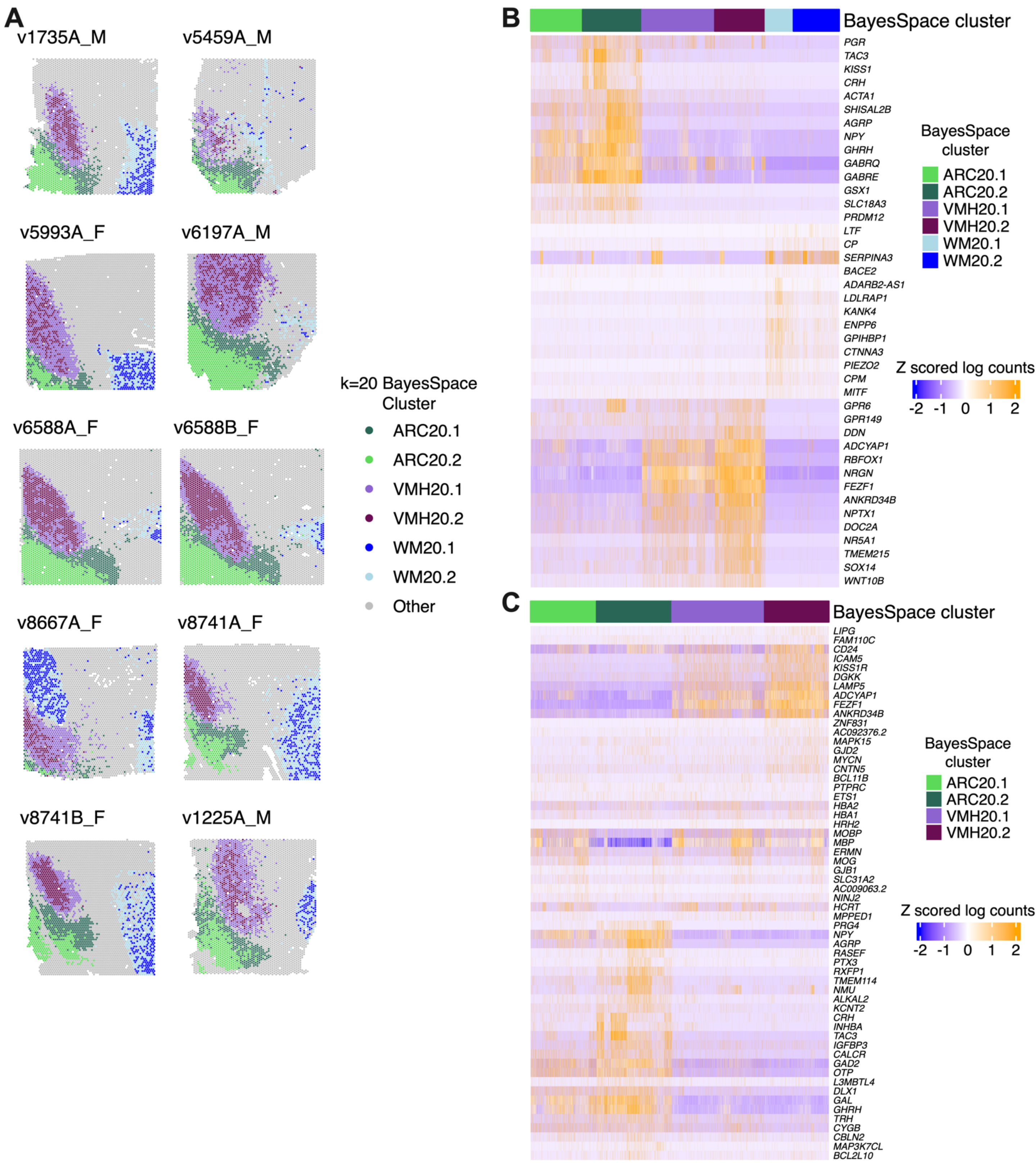


**Figure S6. BayesSpace clustering of Visium data at *k*=20. A)** Spotplots of individual VMH, ARC, and optic tract (OT) white matter (WM) clusters at *k*=20 for all Visium samples. **B)** Heatmap of top marker genes for VMH, ARC, and WM (for comparison) clusters using one vs. all marker gene analyses. **C)** Heatmap of top marker genes using pairwise comparisons between VMH clusters and ARC clusters.

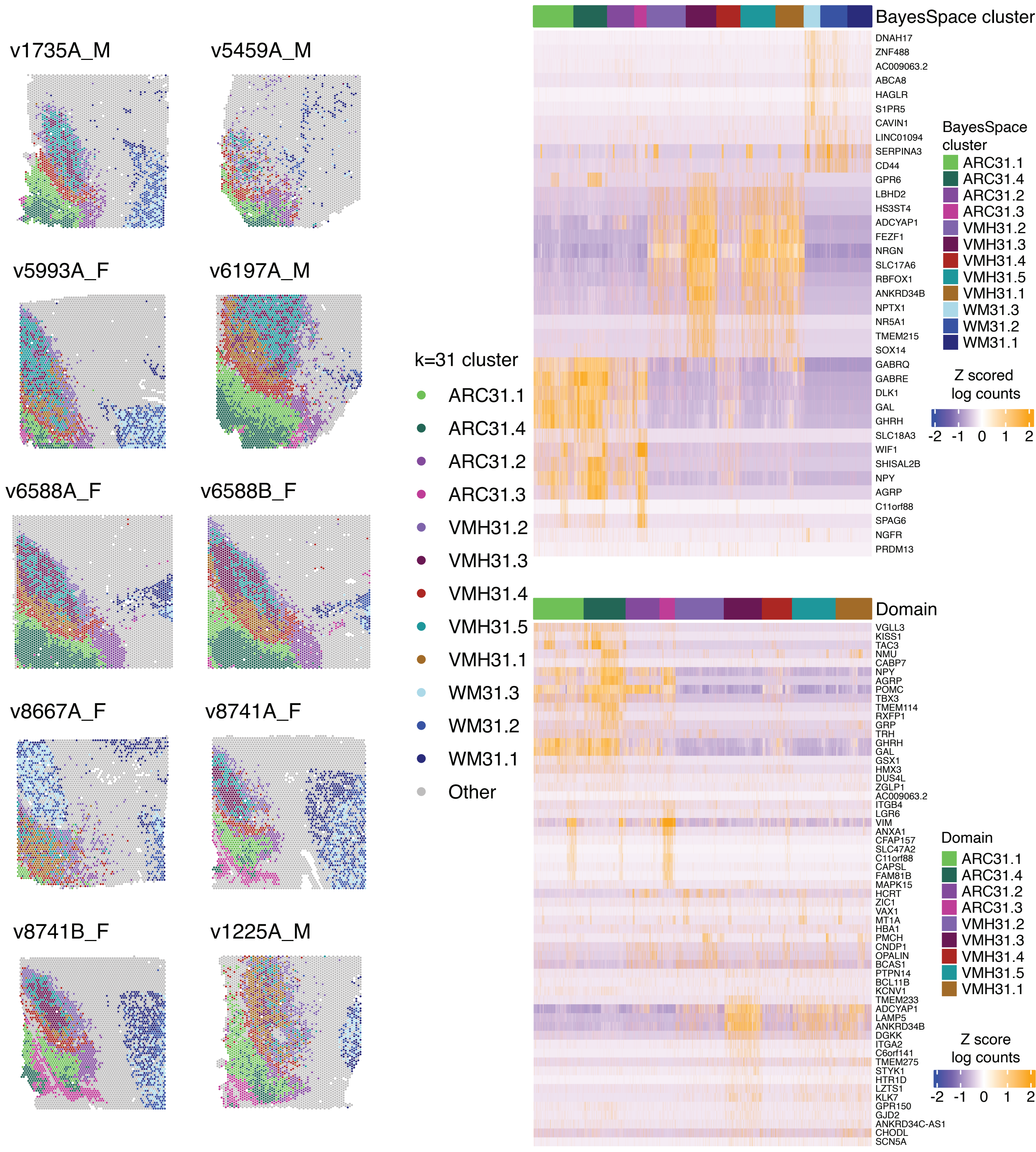


**Figure S7. BayesSpace clustering of Visium data at *k*=31. A)** Spotplots of individual VMH, ARC, and optic tract (OT) white matter (WM) clusters at *k*=31 for all Visium samples. **B)** Heatmap of top marker genes for VMH, ARC, and WM (for comparison) clusters using one vs. all marker gene analyses. **C)** Heatmap of top marker genes when comparing each VMH cluster to all other VMH clusters combined, or each ARC cluster to all other ARC clusters combined.

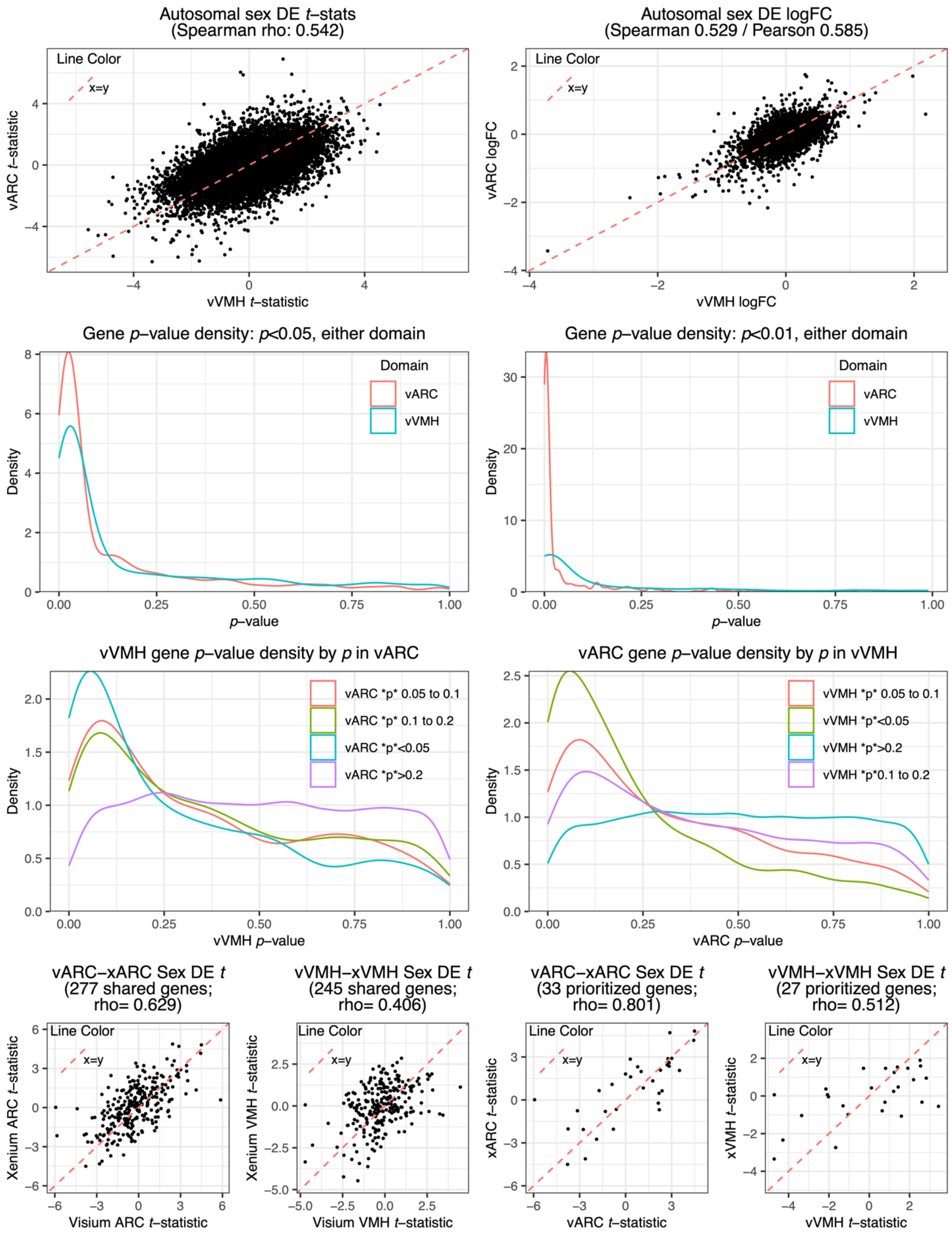


**Figure S8. Shared patterns of sex-DE in vVMH and vARC; reproducibility of sex-DE in ARC and VMH across platforms. A)** Visium domain-domain correlations among sex-DE *t*

statistics or **B)** logFCs. **C)** *p*-value densities for sex-DE analysis considering the union of domain sex-DEGs as called using an uncorrected *p* value threshold of 0.05 or **D)** 0.01. **E-F)** Conditional *p* value density plots for (*i.e.*, vVMH *p* values for genes under a given threshold, plotted for the same genes in vARC, or vice versa). **G)** Visium-Xenium correlation of domain-level sex-DE analyses among all shared genes or **H)** Genes earmarked for sex-DE follow-up with Xenium during panel design based on preliminary Visium sex-DE analyses.

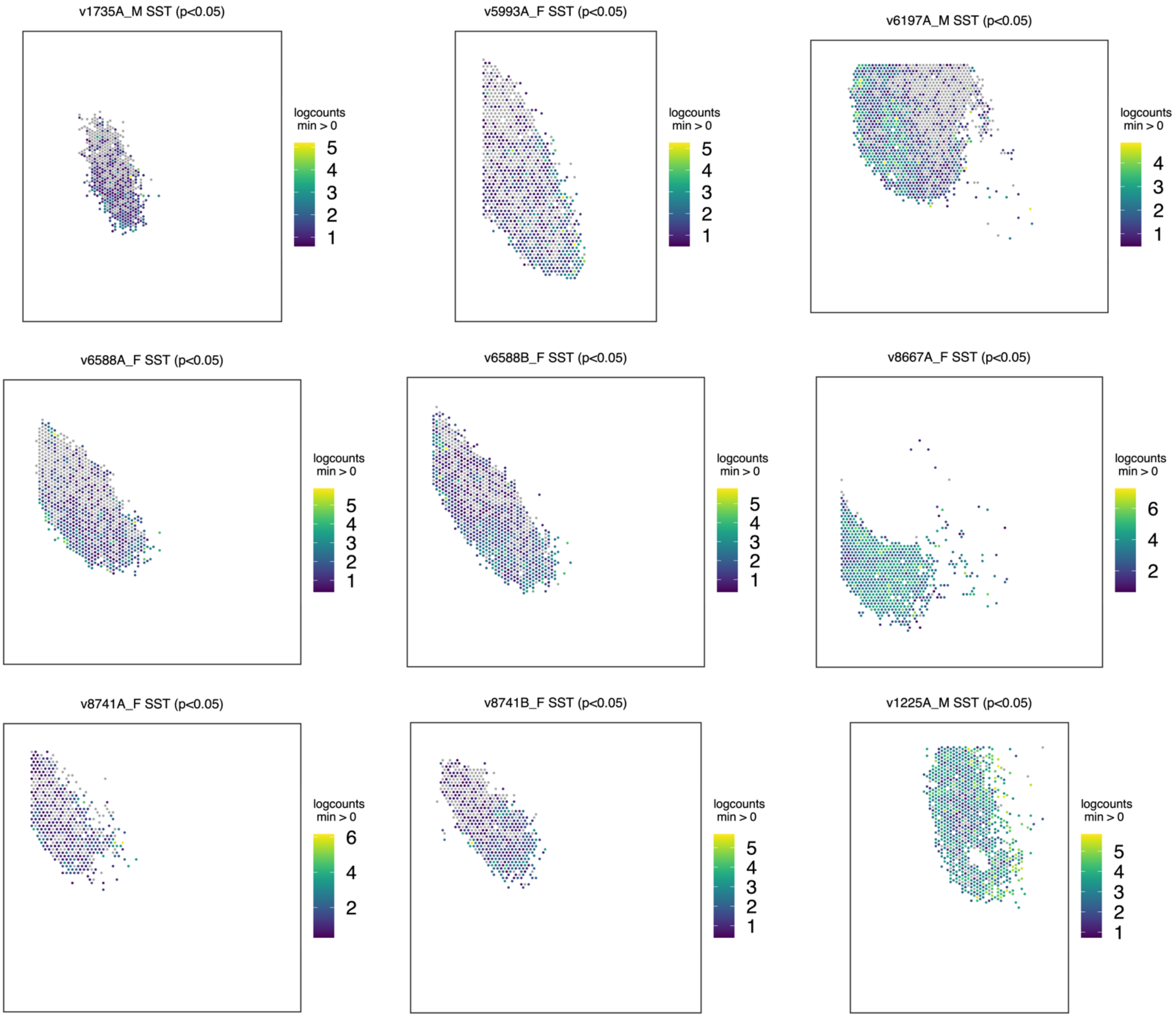


**Figure S9. Spatially variable genes (SVGs) within vVMH: *SST*.** Spotplots of *SST* in vVMH for each sample. *SST* is a within-vVMH SVG that appears ventrally biased. Sample titles indicate whether spatial variation was significant. Expression is only shown for spots labeled as vVMH for readability.

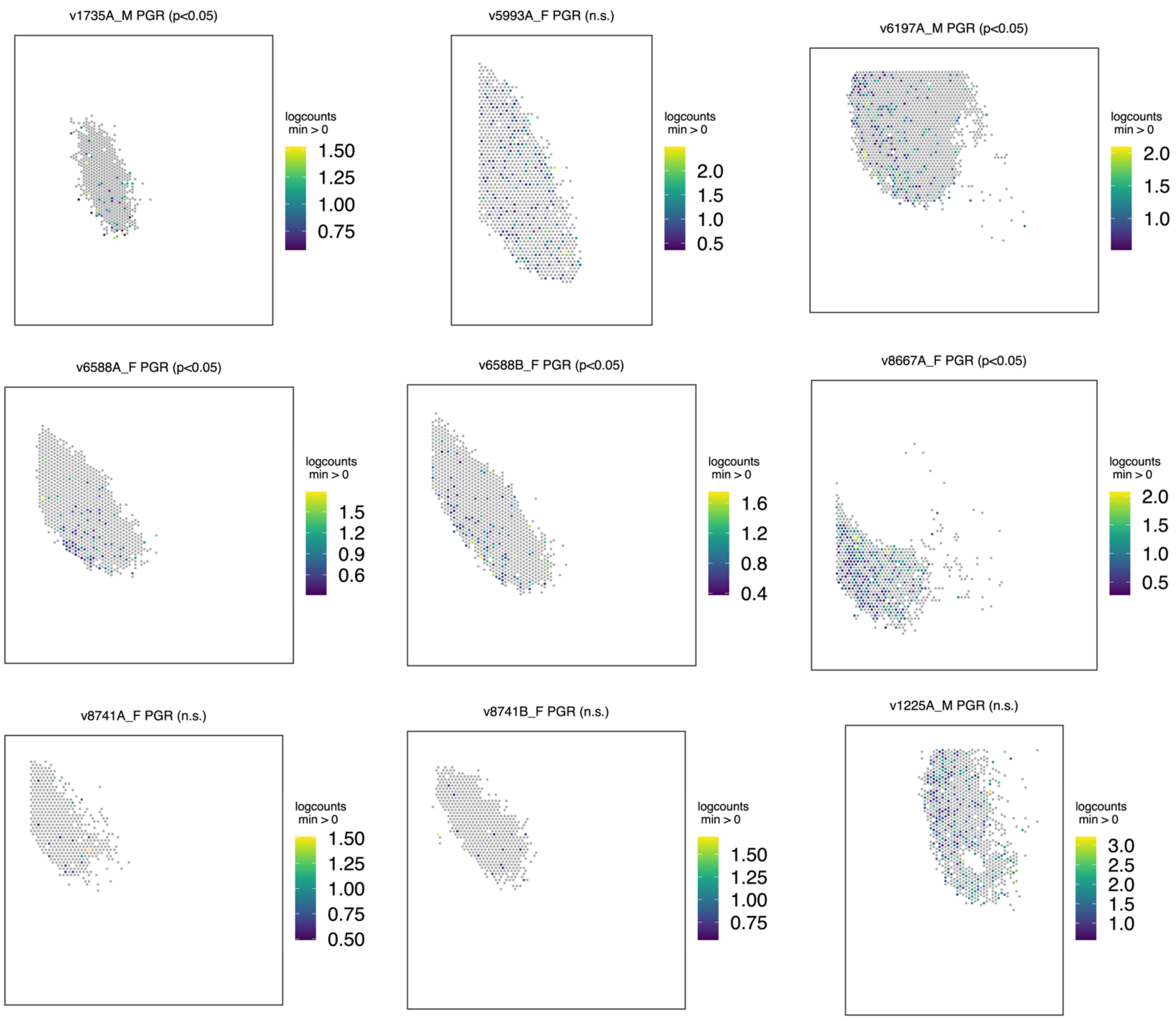


**Figure S10. Visium SVGs within vVMH: *PGR*.** Spotplots of *PGR* in vVMH for each sample. *PGR* is a within-vVMH SVG that appears ventrally to ventro-medially biased. Sample titles indicate whether spatial variation was significant. Expression is only shown for spots labeled as vVMH for readability.

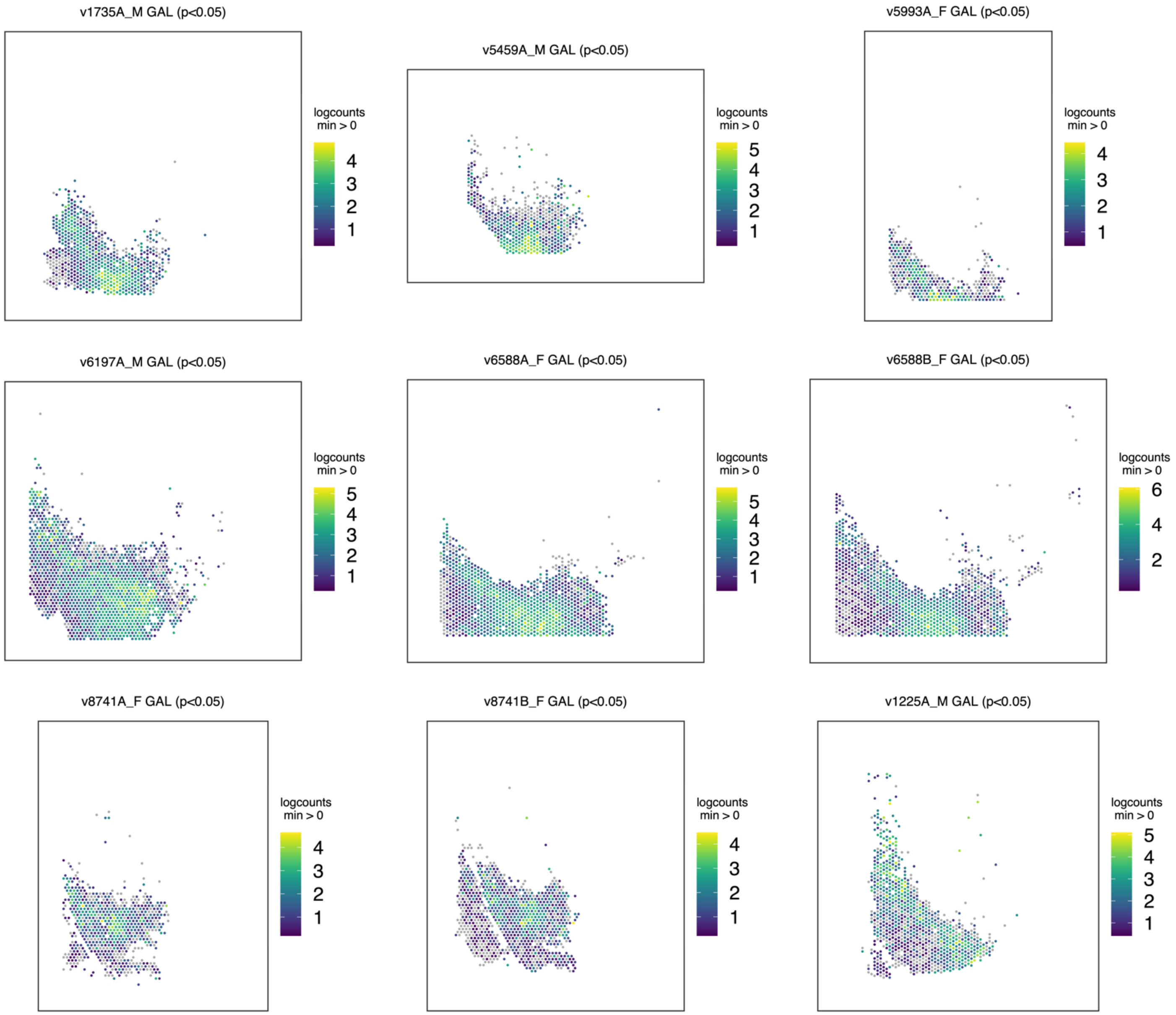


**Figure S11. Visium SVGs within vARC: *GAL***. Spotplots of *GAL* in vARC for each sample. *GAL* is a within-vARC SVG that appears biased to lateral ARC (along long medial edge of VMH) and center of ARC (ventral to VMH). Sample titles indicate whether spatial variation was significant. Expression is only shown for spots labeled as vARC for readability.

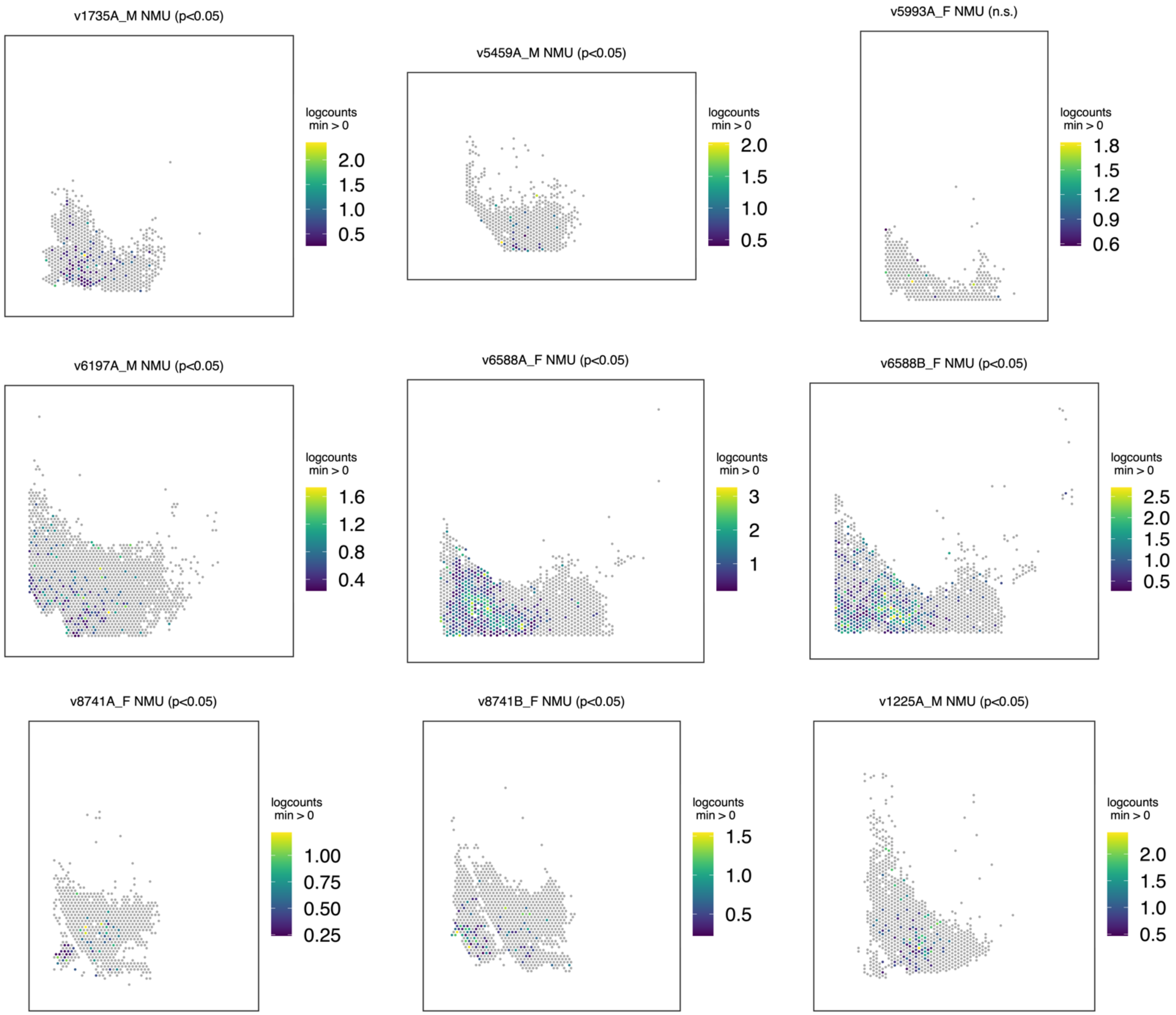


**Figure S12. Visium SVGs within vARC: *NMU*.** Spotplots of *NMU* in vARC for each sample. *NMU* is a within-vARC SVG with ventral localization in the center portion of ARC. Sample titles indicate whether spatial variation was significant. Expression is only shown for spots labeled as vARC for readability.

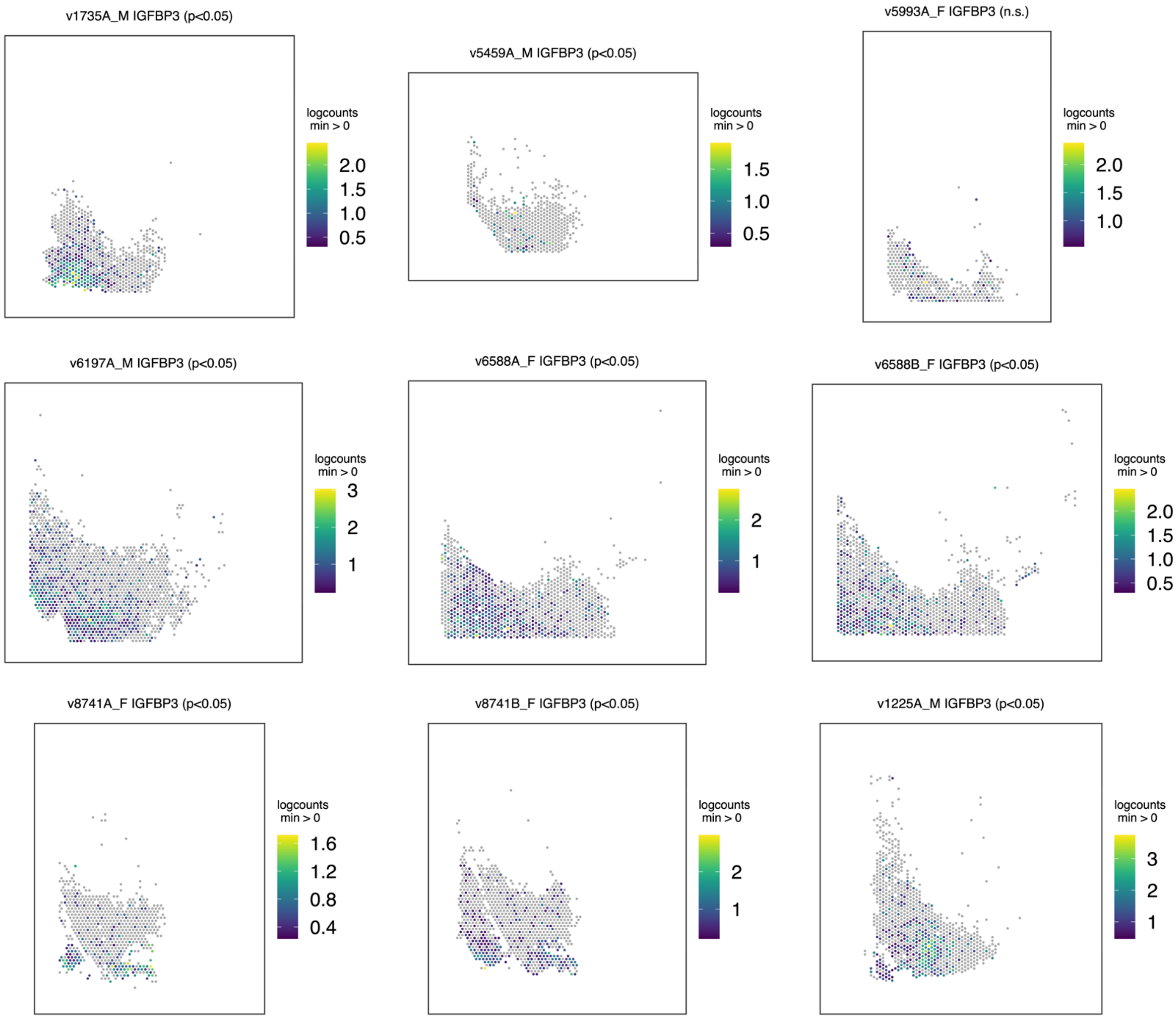


**Figure S13. Visium SVGs within vARC: *IGFBP3*.** Spotplots of *IGFBP3* in vARC for each sample. *IGFBP3* is a within-vARC SVG that appears ventromedially biased. Sample titles indicate whether spatial variation was significant. Expression is only shown for spots labeled as vARC for readability.

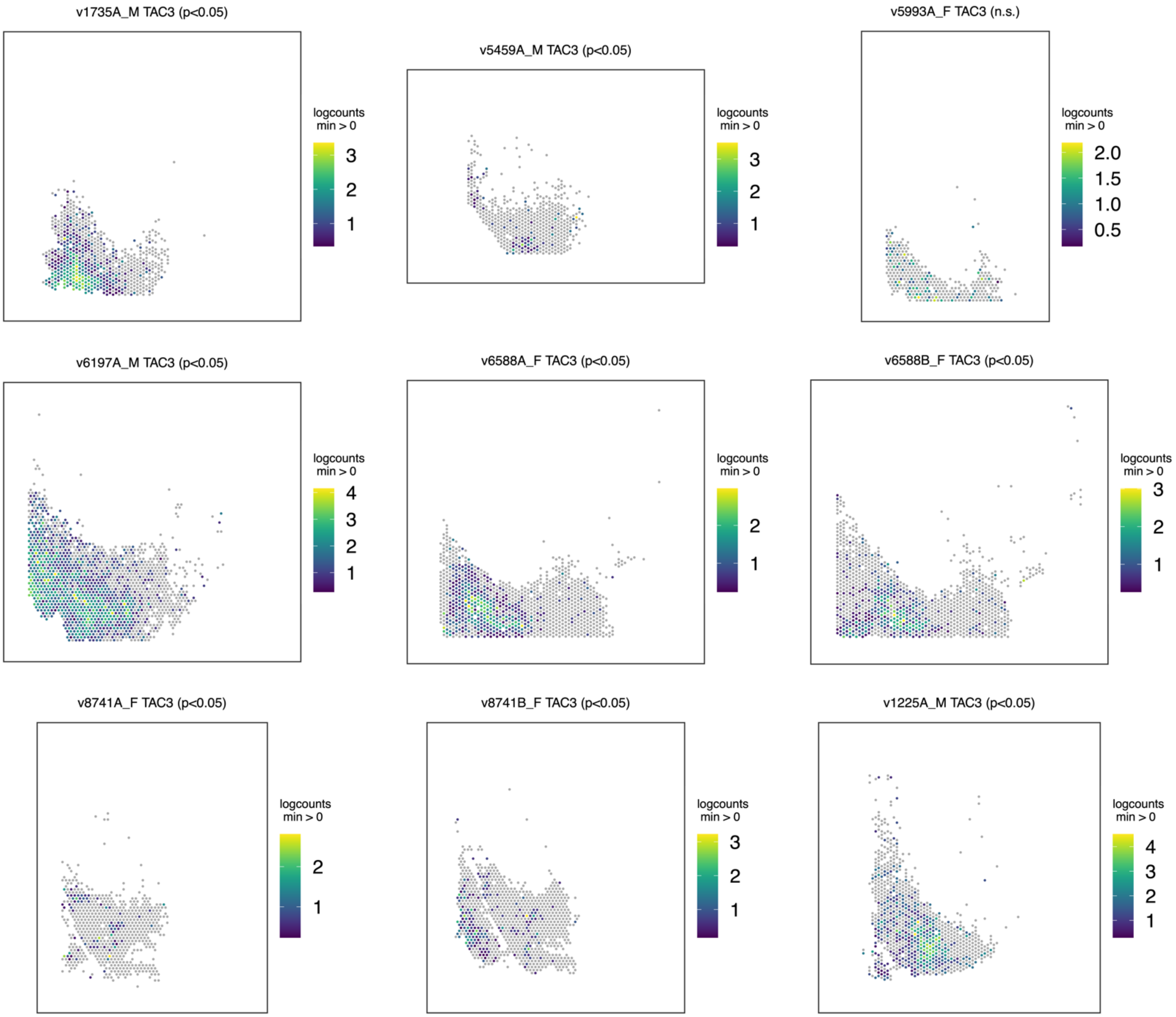


**Figure S14. Visium SVGs within vARC: *TAC3*.** Spotplots of *TAC3* in vARC for each sample. *TAC3* is a within-vARC SVG (ventral localization in center ARC, similar to *NMU*), for each sample. Sample titles indicate whether spatial variation was significant. Expression is only shown for spots labeled as vARC for readability.

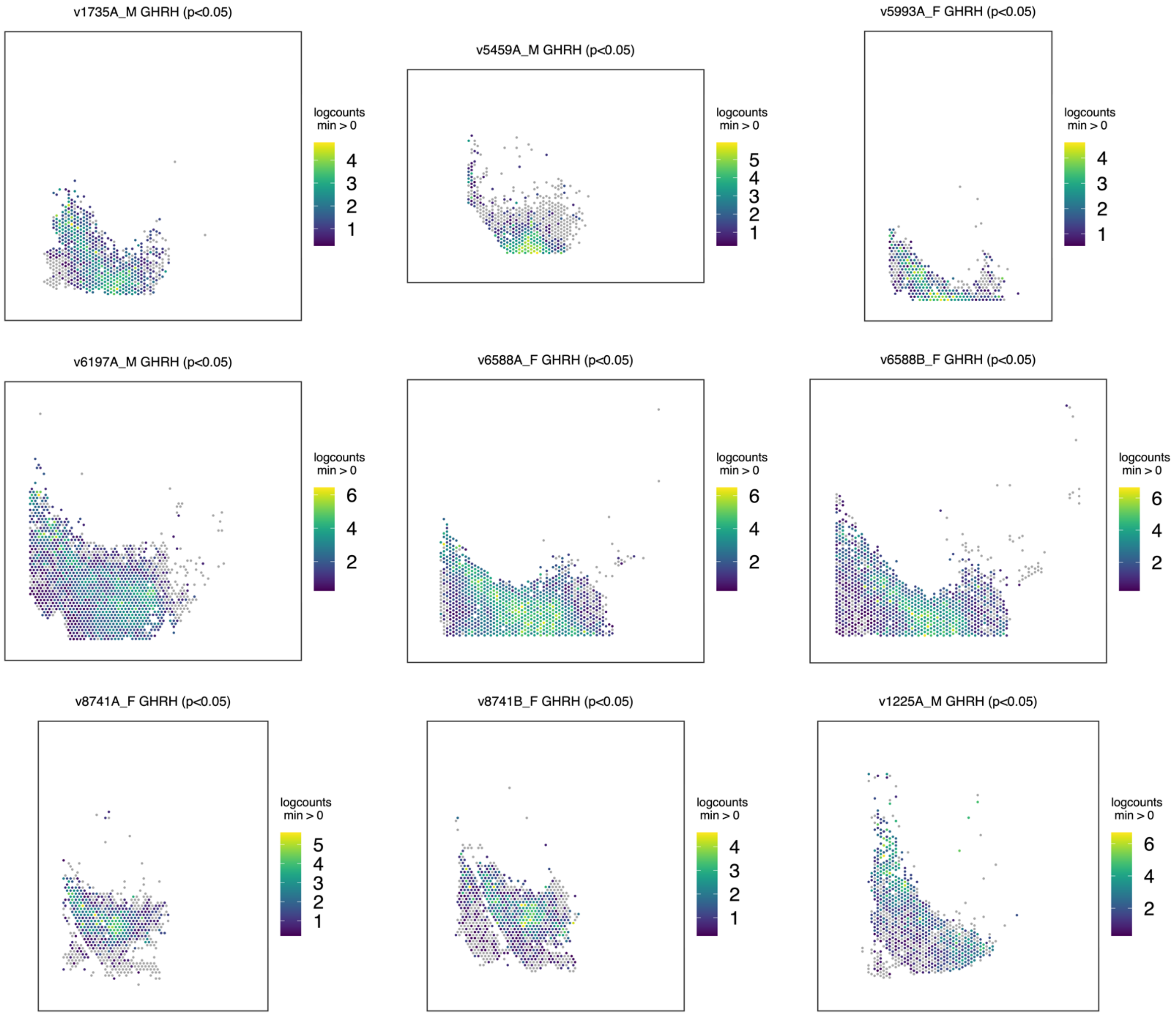


**Figure S15. Visium SVGs within vARC: *GHRH*.** Spotplots of *GHRH* in vARC for each sample. *GHRH* is a within-vARC SVG that appears ventrally biased. Sample titles indicate whether spatial variation was significant. Expression is only shown for spots labeled as vARC for readability.

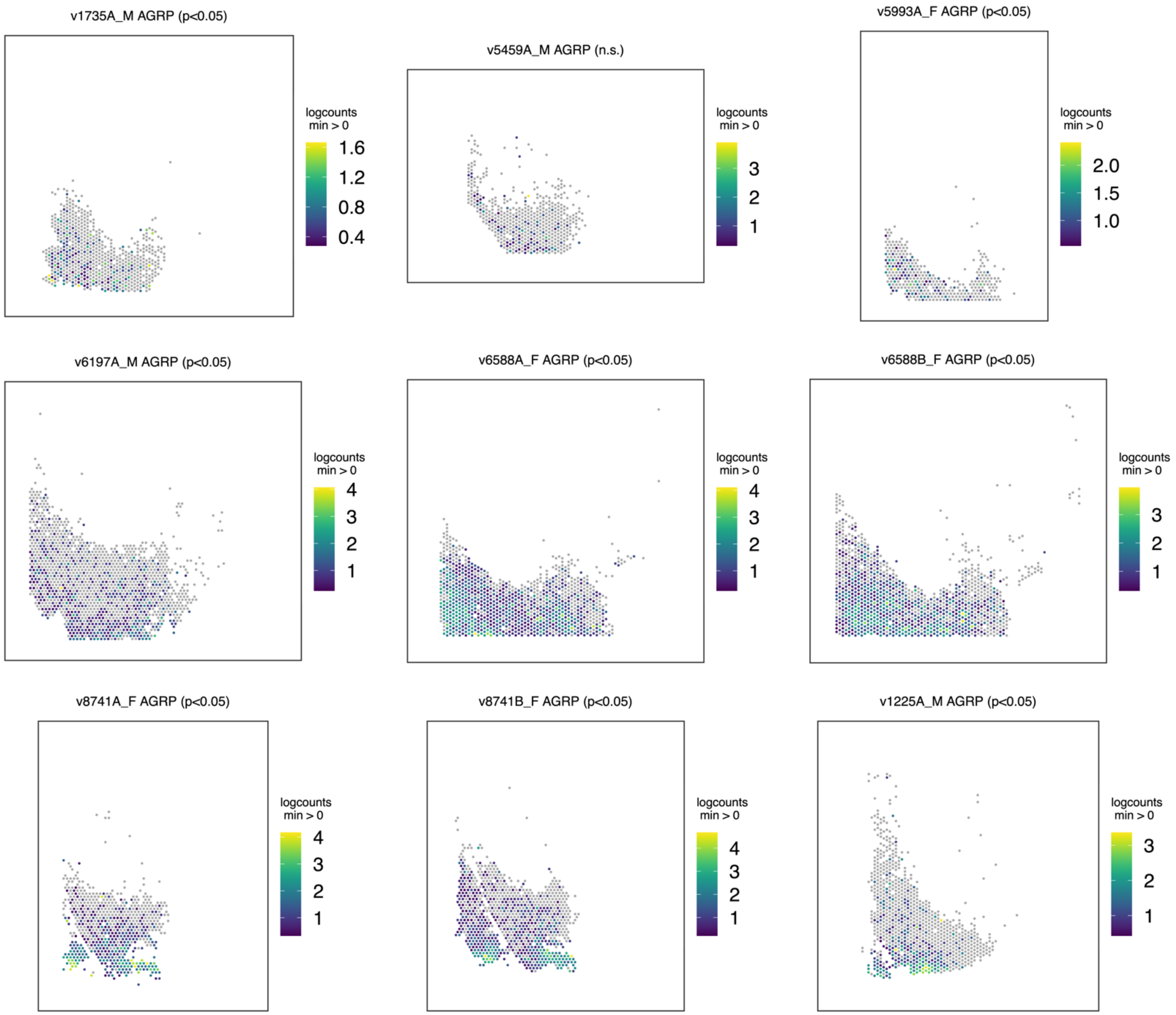


**Figure S16. Visium SVGs within vARC: *AGRP*.** Spotplots of *AGRP* in vARC for each sample. *AGRP* is a within-vARC SVG that shows a dorsal-to-ventral gradient. Sample titles indicate whether spatial variation was significant. Expression is only shown for spots labeled as vARC for readability.

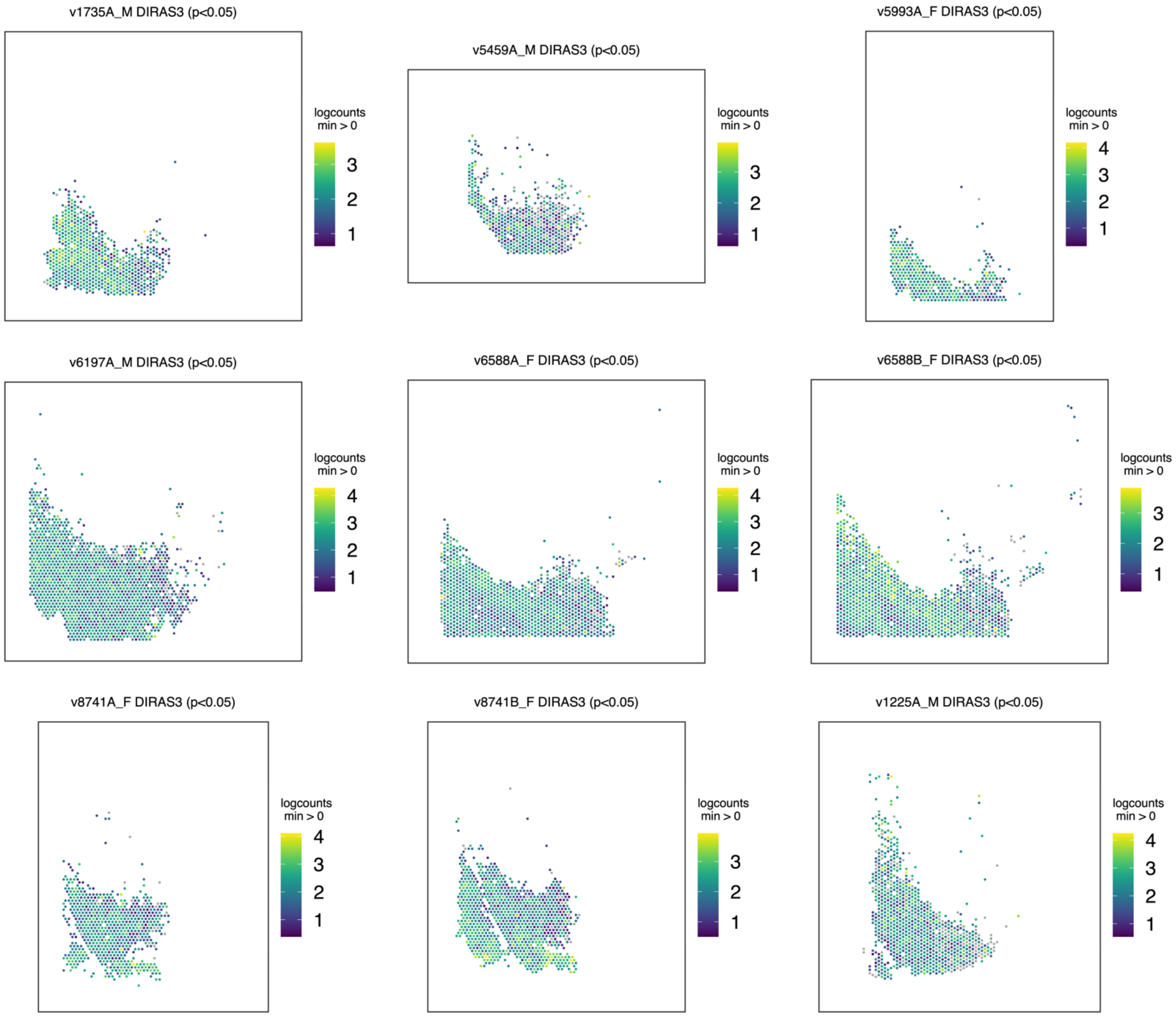


**Figure S17. Visium SVGs within vARC: *DIRAS3*.** Spotplots of *DIRAS3* in vARC for each sample. *DIRAS3* is a within-vARC SVG that shows a central band of expression in ARC with exclusion from lateralmost area. Sample titles indicate whether spatial variation was significant. Expression is only shown for spots labeled as vARC for readability.

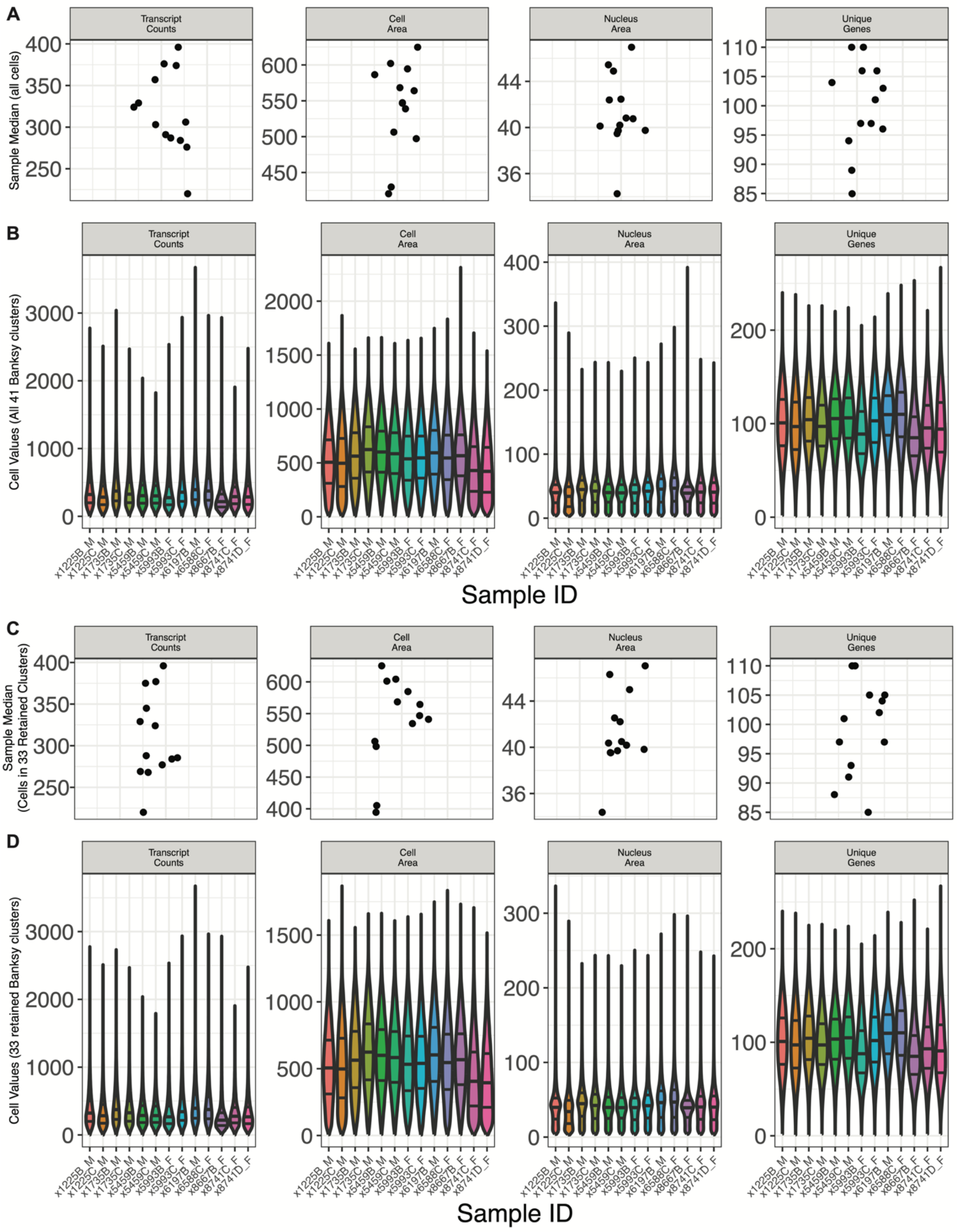


**Figure S18. Quality control metrics from the final Xenium dataset. (*Continued on next page*)**

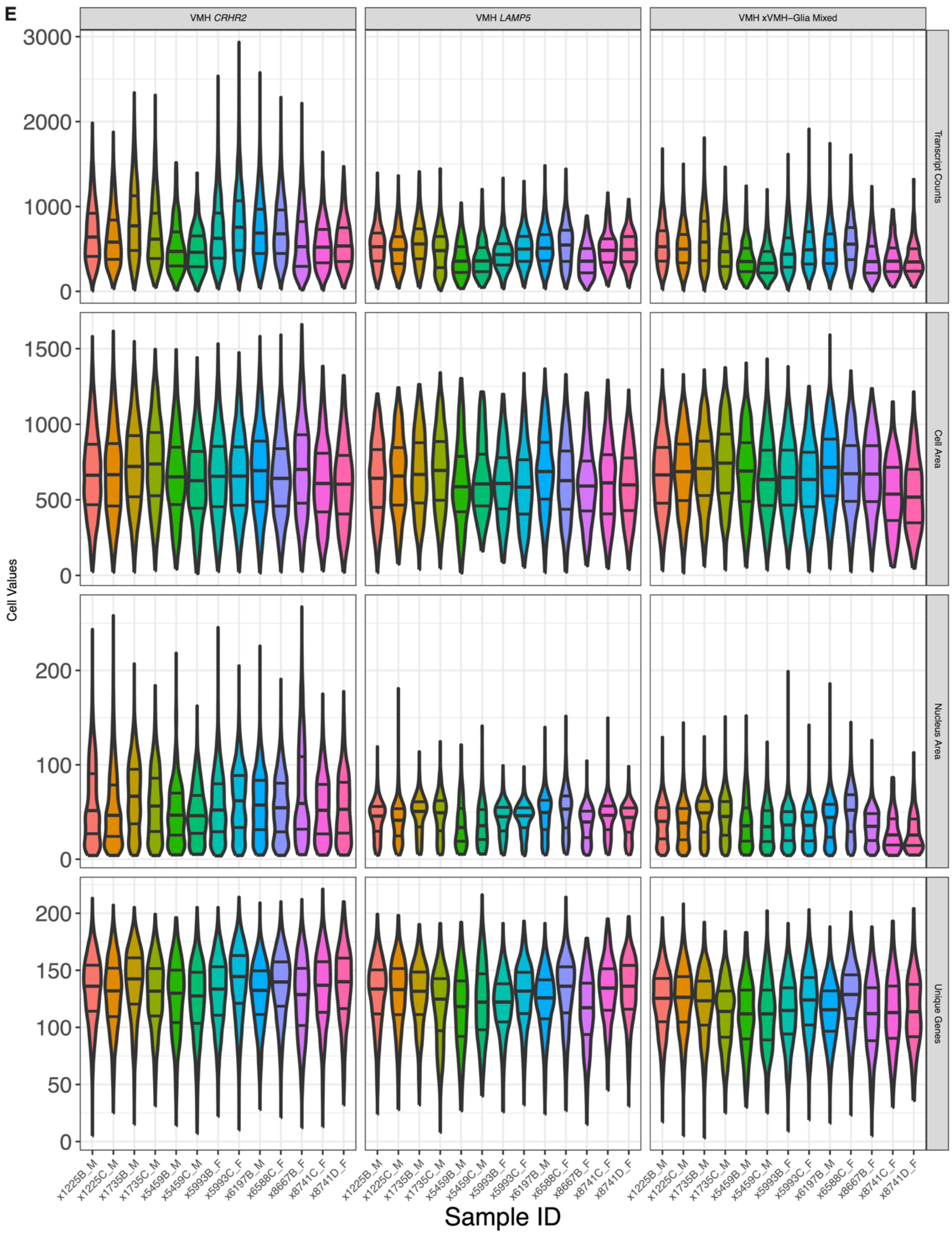


**Figure S18. Quality control metrics from the final Xenium dataset.** (*Continued on next page*)

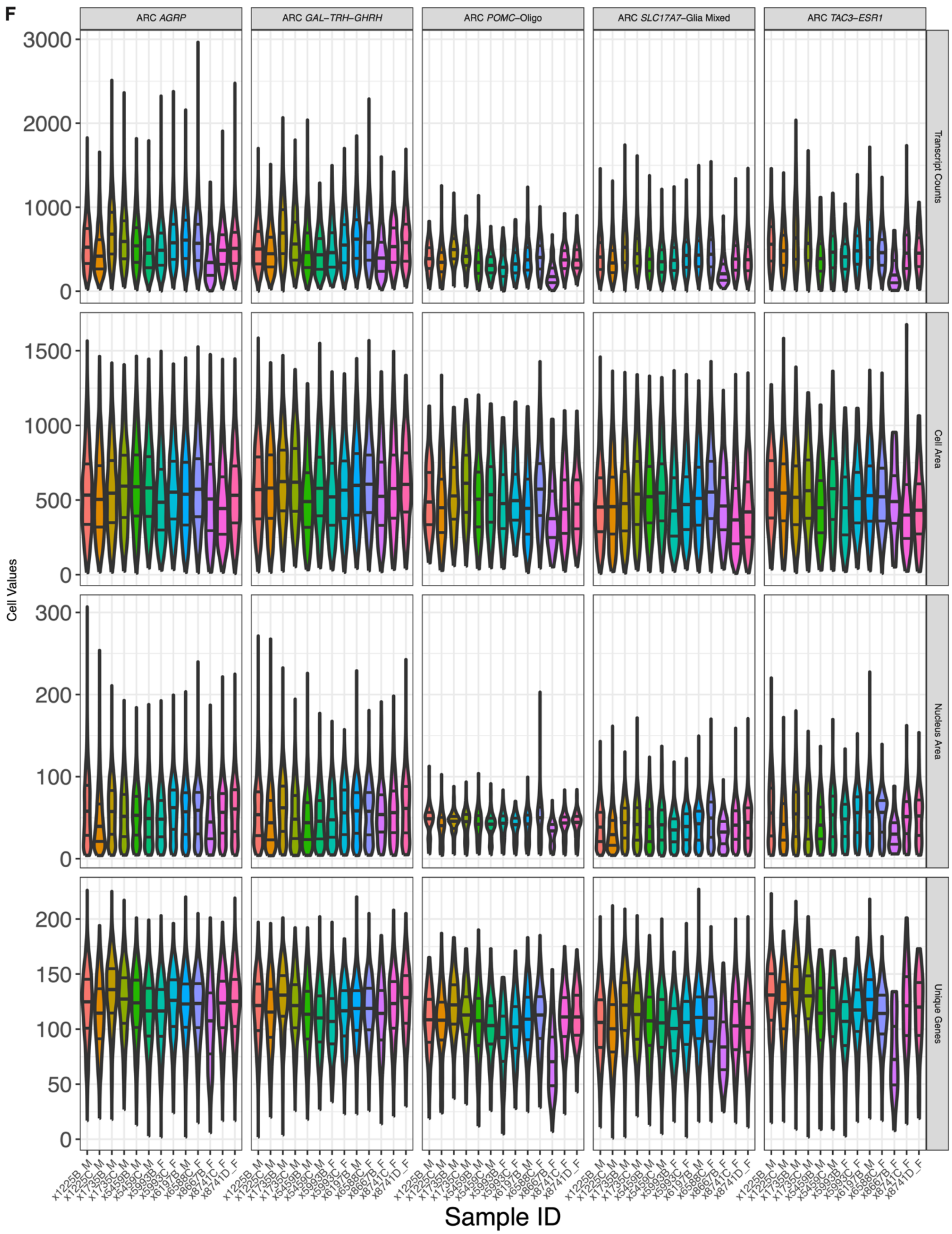


**Figure S18. Quality control metrics from the final Xenium dataset.** Number of transcripts detected per cell, segmented cell area, segmented nucleus area, and number of unique genes per cell are shown for various subsets of the Xenium data. Across all 41 cell clusters initially resulting from *Banksy* analysis, shown are **A)** samplewise medians and **B)** the distributions of these cell-level metrics within samples. After excluding eight sample/donor-specific *Banksy* clusters, shown are **C)** samplewise medians and **D)** the distributions of these cell-level metrics within samples. **E)** Violin plots of these metrics are shown for each xARC neuron cluster. Clusters are named along the top and metrics are listed along the right margin. **F)** Violin plots of these metrics are shown for each xVMH neuron cluster. Clusters are named along the top and metrics are listed along the right margin.

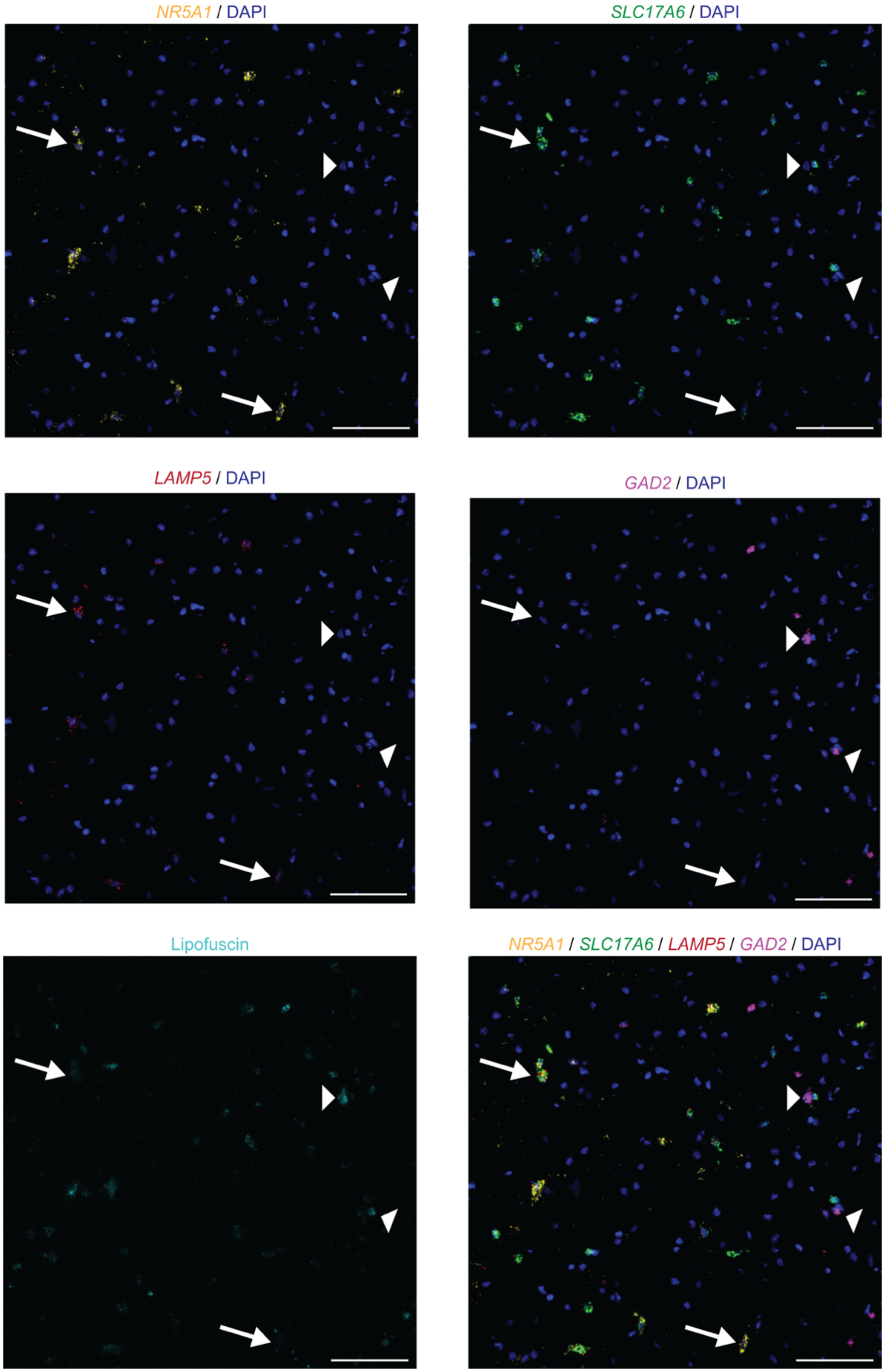


**Figure S19. *LAMP5* is expressed in *SLC17A6*+ excitatory neurons of VMH, but not local *GAD2*+ neurons.** Representative confocal image showing expression of *GAD2*, *LAMP5*, *NR5A1*, and *SLC17A6* in the VMH by smFISH on an adjacent tissue from Br1735. Each gene is shown individually overlaid with nuclear DAPI staining along with a merge of all channels. Arrows indicate example cells co-expressing *NR5A1*, *LAMP5*, and *SLC17A6,* but negative for *GAD2*; arrowheads indicate example *GAD2*-expressing cells. Lipofuscin autofluorescence (without DAPI) is shown in the bottom left image. Scale bar: 100µm.

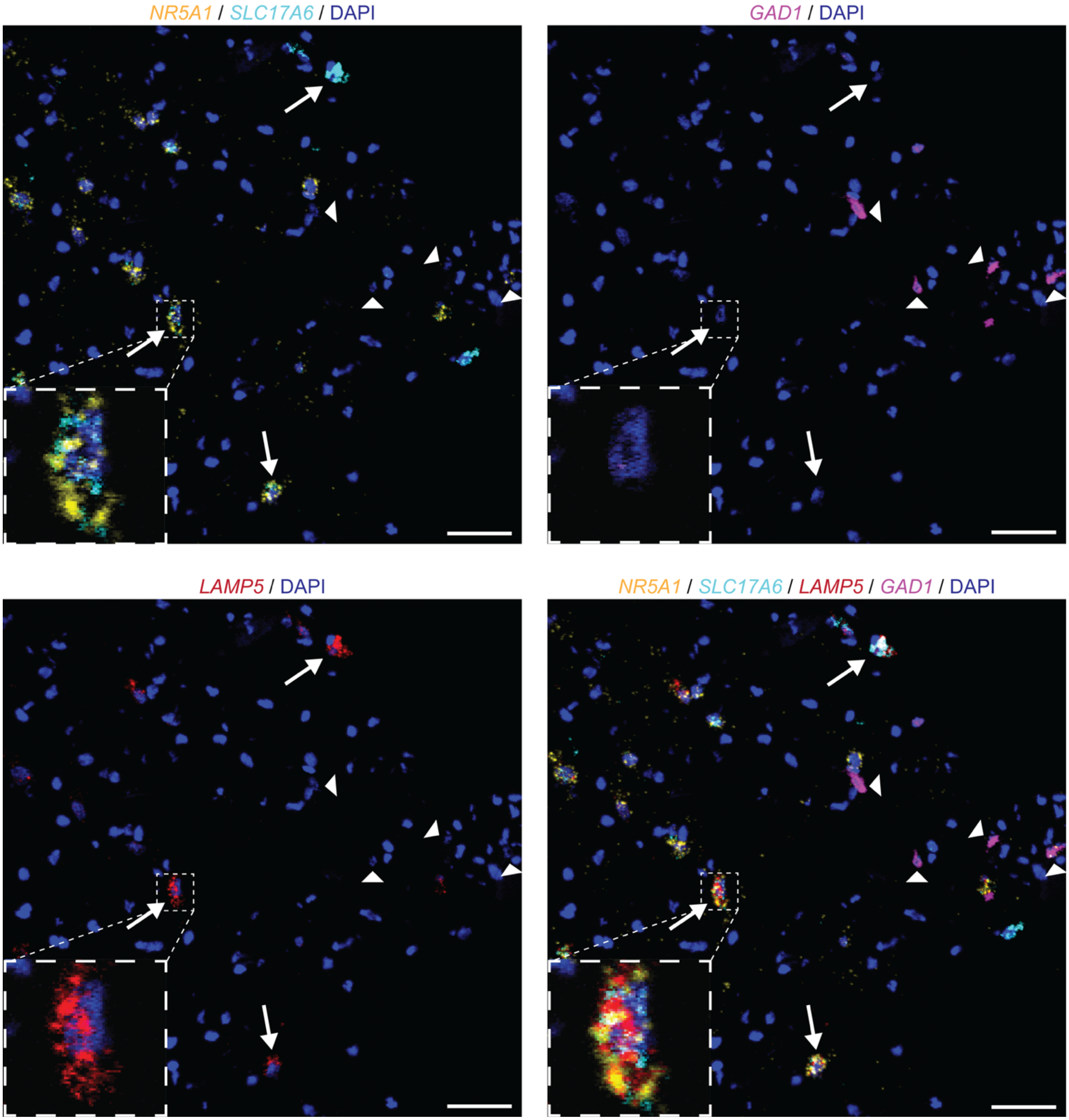


**Figure S20. *LAMP5* is expressed in *SLC17A6*+ excitatory neurons of VMH, but not local *GAD1*+ neurons.** Representative confocal image showing expression of *NR5A1*, *SLC17A6*, *GAD1*, and *LAMP5* by smFISH in an adjacent tissue section from Br1735. Arrows indicate *LAMP5*+ excitatory VMH neurons; arrowheads indicate *GAD1*+ inhibitory neurons ventral/lateral to VMH. Scale bar: 50µm.

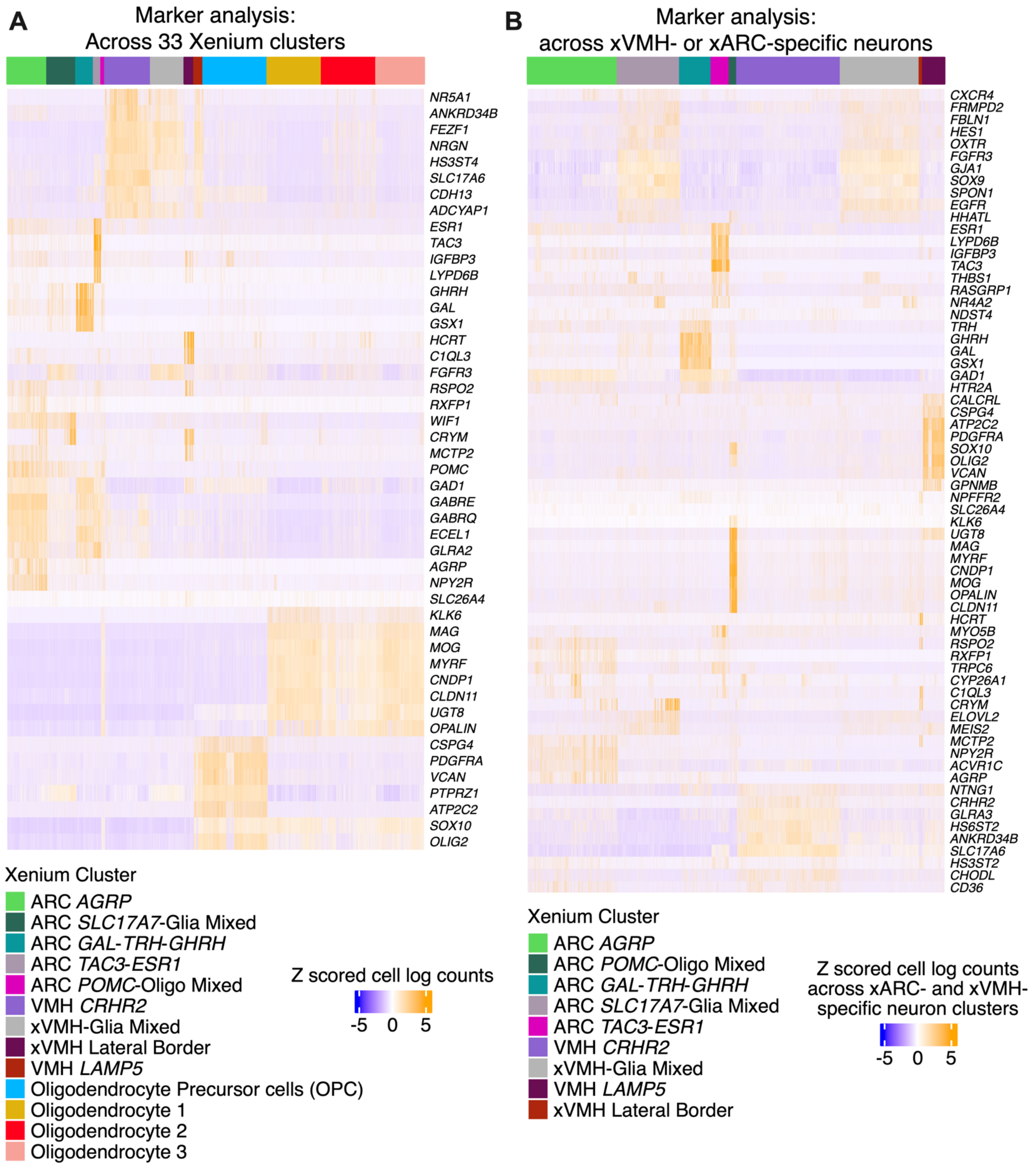


**Figure S21. Marker gene heatmaps of selected xVMH and xARC cell clusters. A)** Top gene markers of xVMH and xARC clusters alongside those of white matter (WM) clusters for comparison, as identified using one-vs.-all analysis across all 33 retained Xenium clusters. **B)** Top gene markers for each xVMH or xARC neuron cluster (constrained to cells with those labels falling within their domain boundaries), compared only to other xVMH or xARC neuron clusters, respectively.

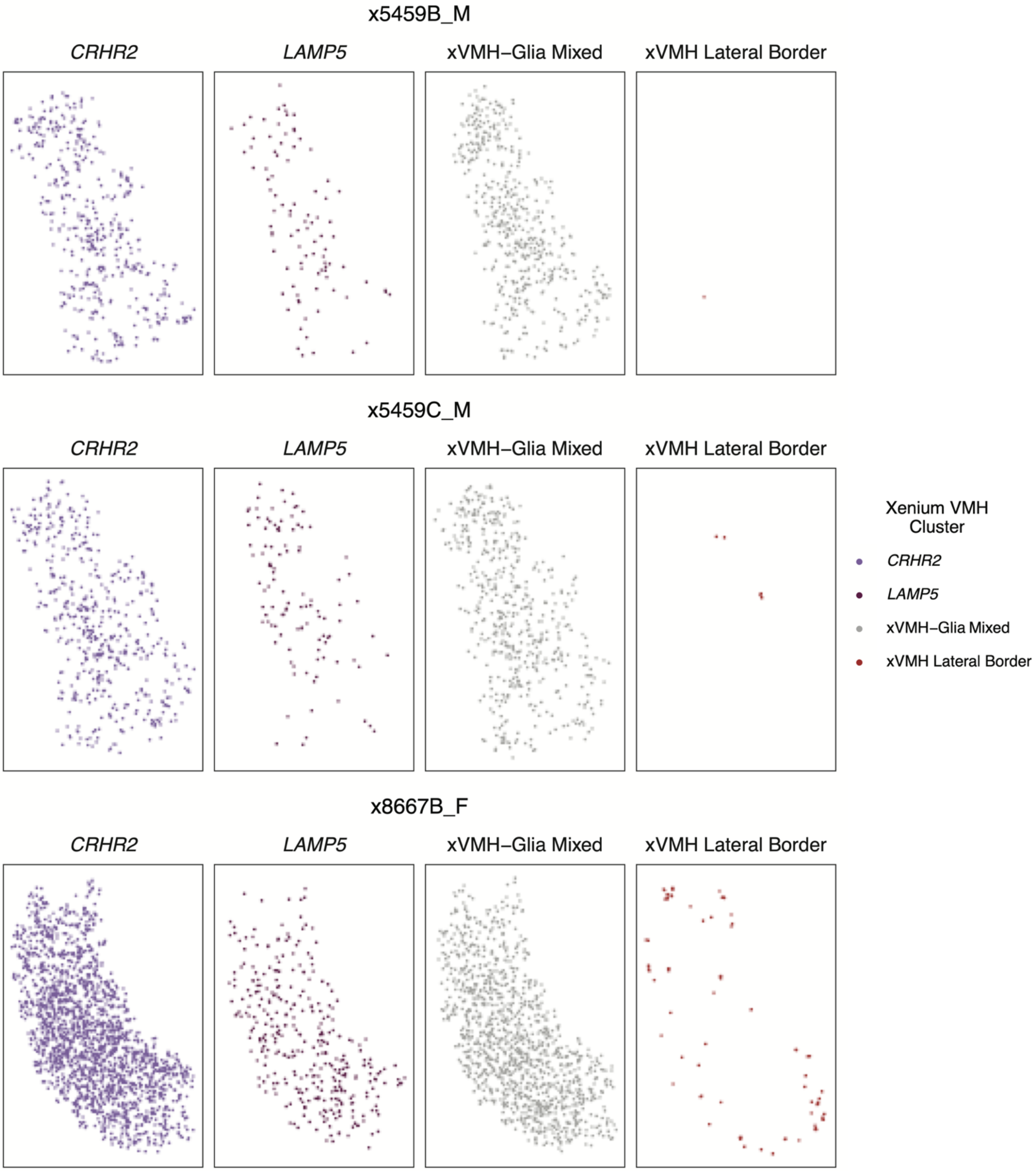


**Figure S22. Spatial organization of xVMH cell clusters plotted within the xVMH domain for all samples.** **(*Continued on next page*)**

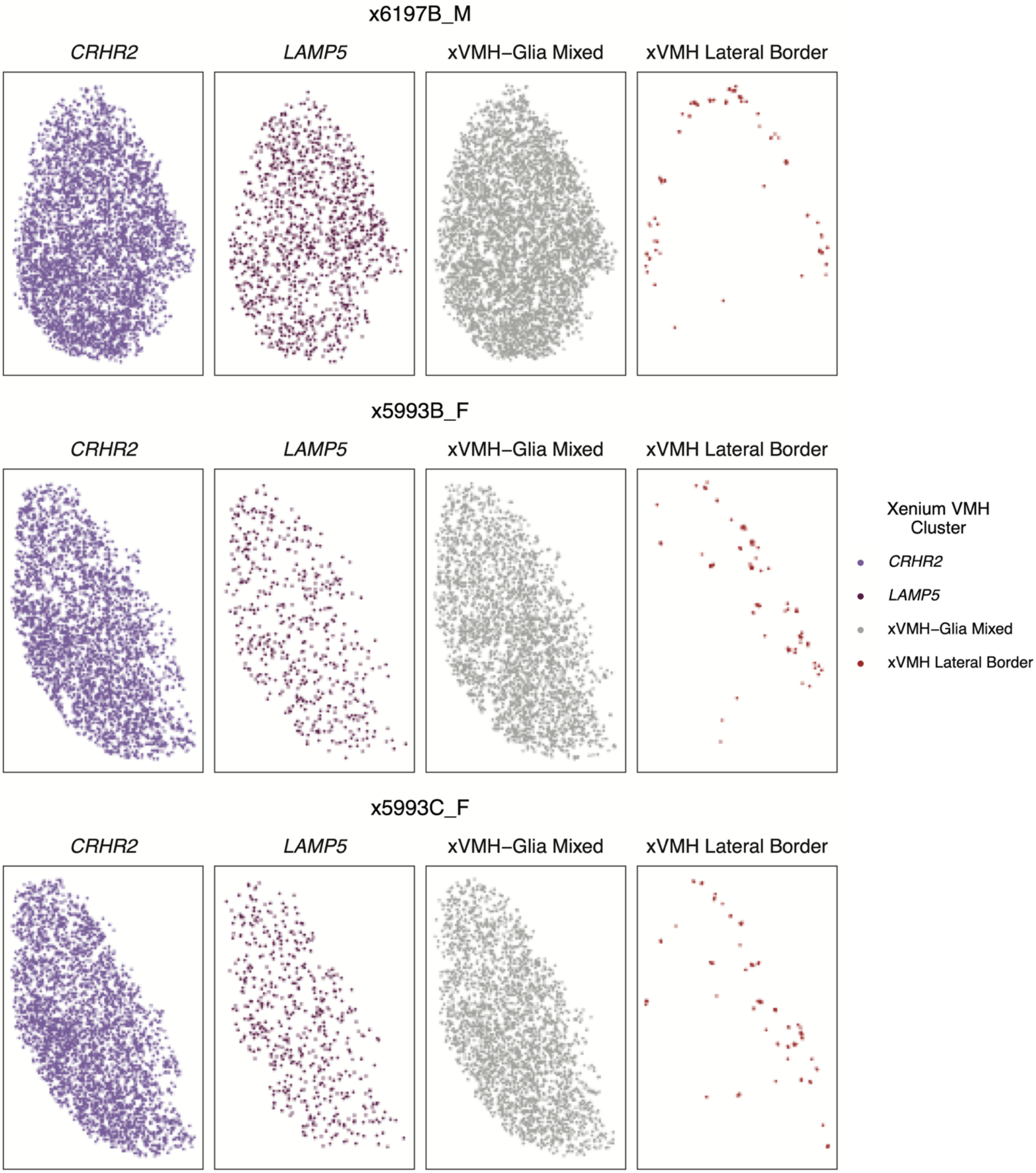


**Figure S22. Spatial organization of xVMH cell clusters plotted within the xVMH domain for all samples.** **(*Continued on next page*)**

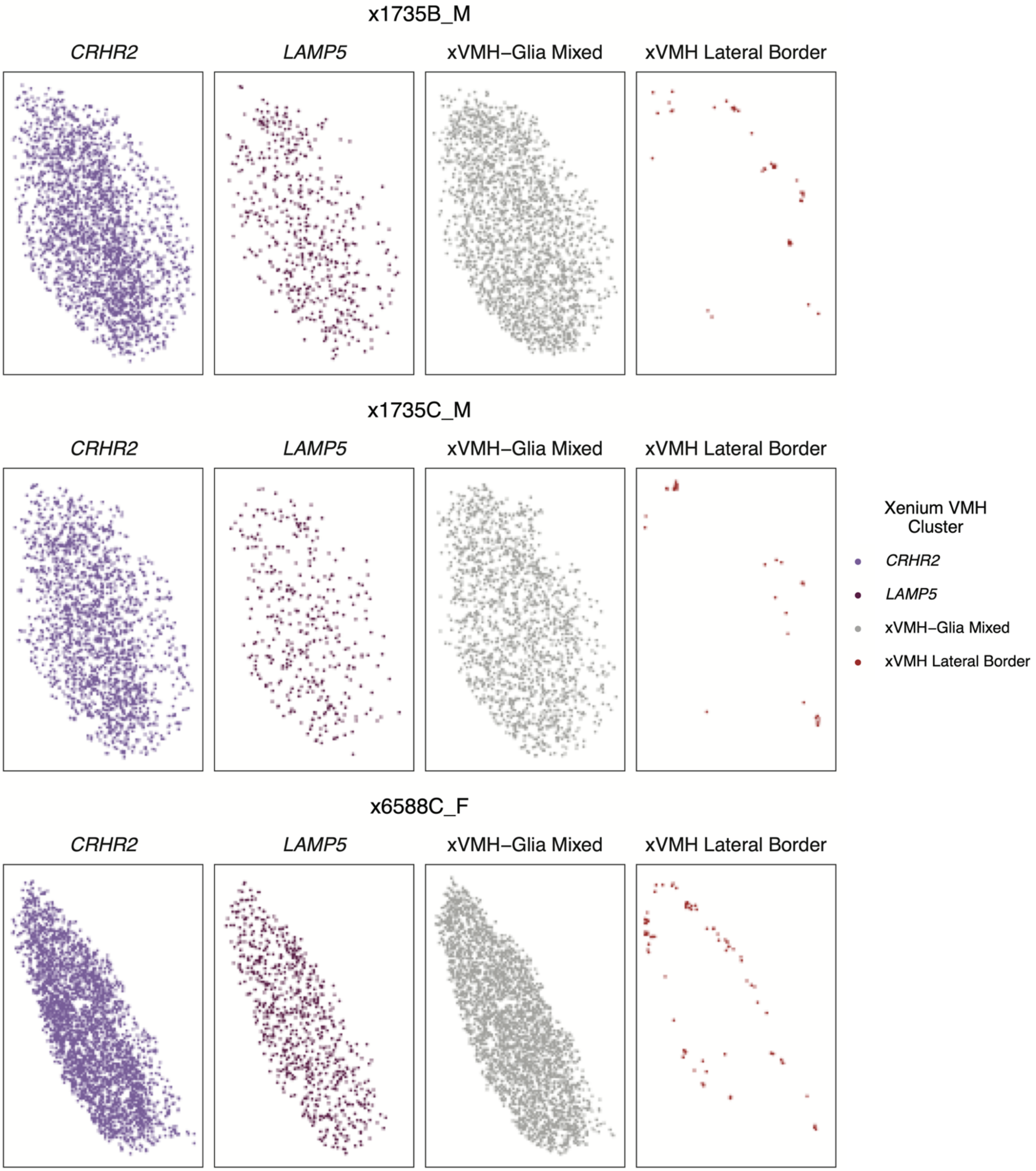


**Figure S22. Spatial organization of xVMH cell clusters plotted within the xVMH domain for all samples.** **(*Continued on next page*)**

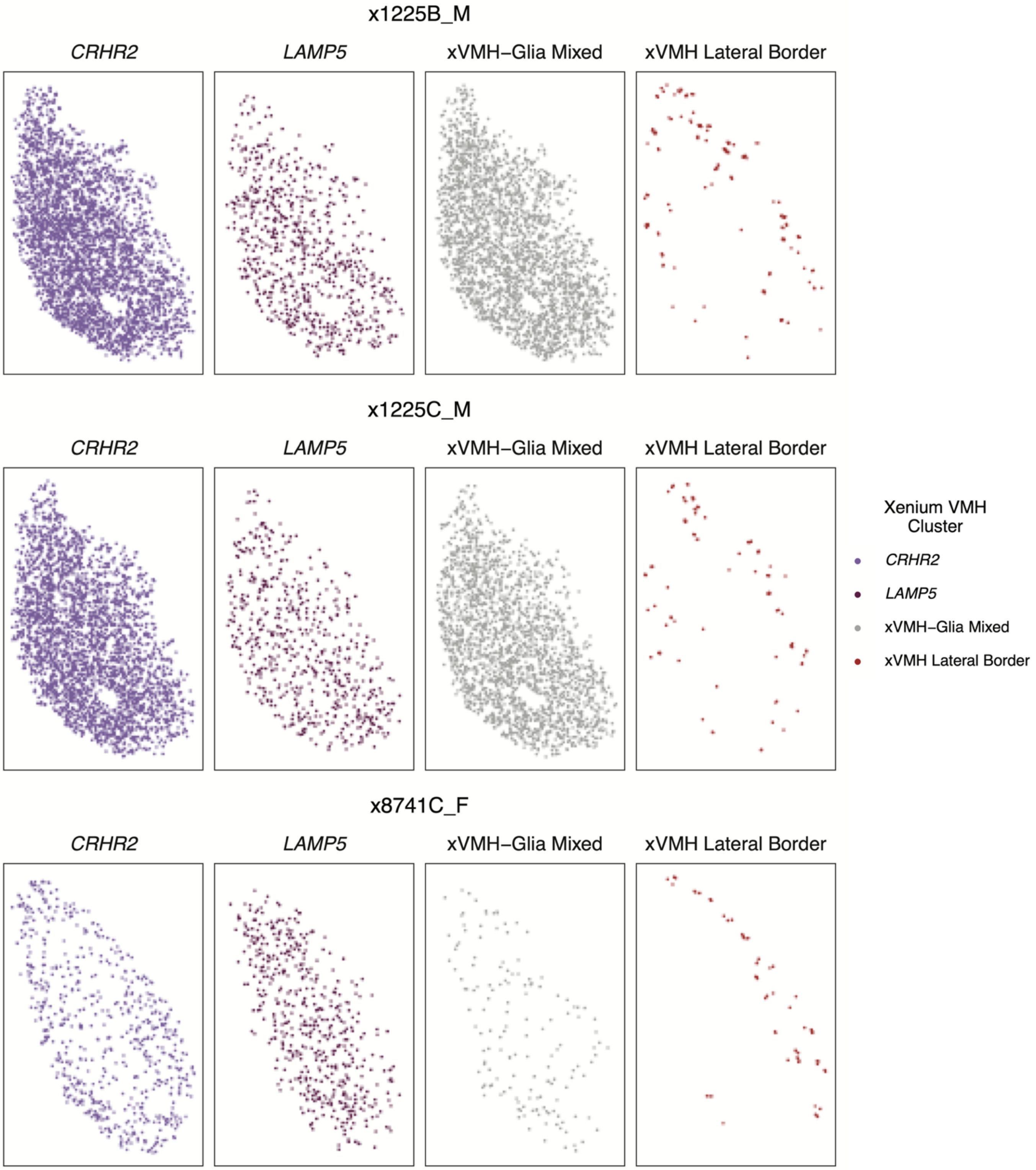


**Figure S22. Spatial organization of xVMH cell clusters plotted within the xVMH domain for all samples.** **(*Continued on next page*)**

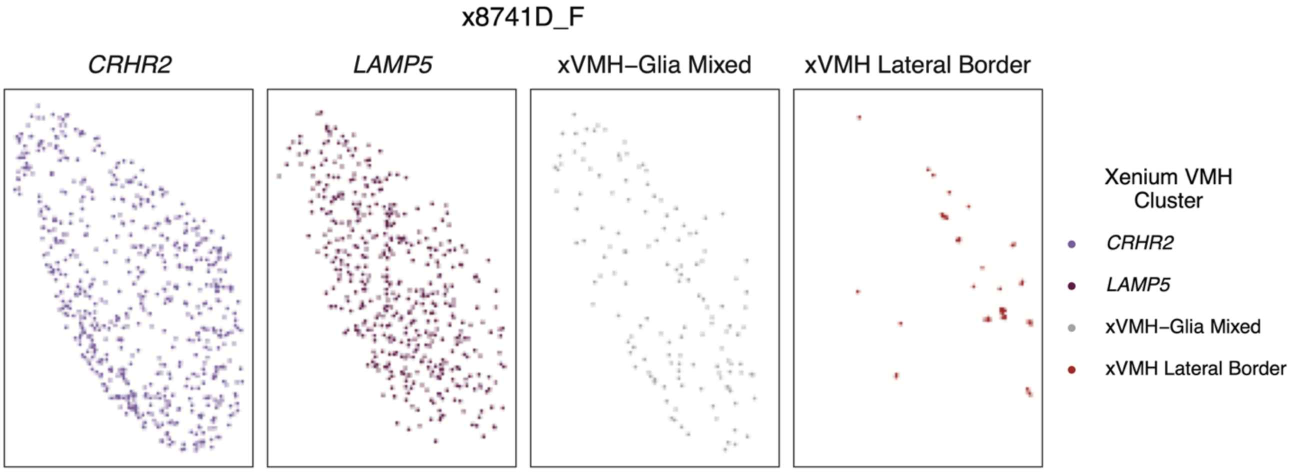


**Figure S22. Spatial organization of xVMH cell clusters plotted within the xVMH domain for all samples.** For each sample, plots illustrate the location of each cell from *CRHR2*, *LAMP5*, xVMH-Glia Mixed, and xVMH Lateral Border clusters. Only cells within the xVMH domain area are plotted. Samples from donor Br8741 contained a sample-specific VMH cluster (not shown) overlapping those displayed; such cells were excluded from all analyses.

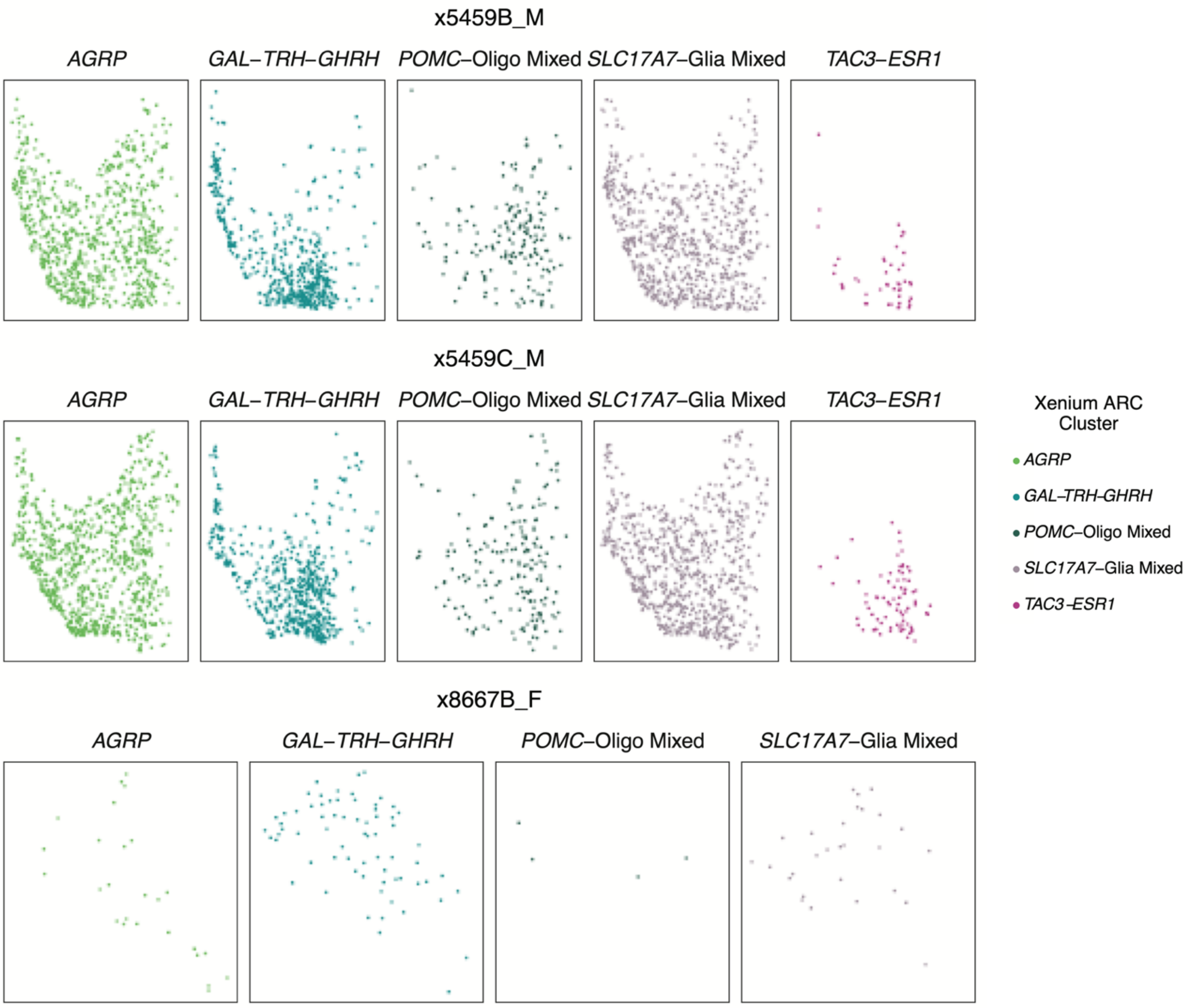


**Figure S23. Spatial organization of xARC cell clusters plotted within the xARC domain for all samples.** **(*Continued on next page*)**

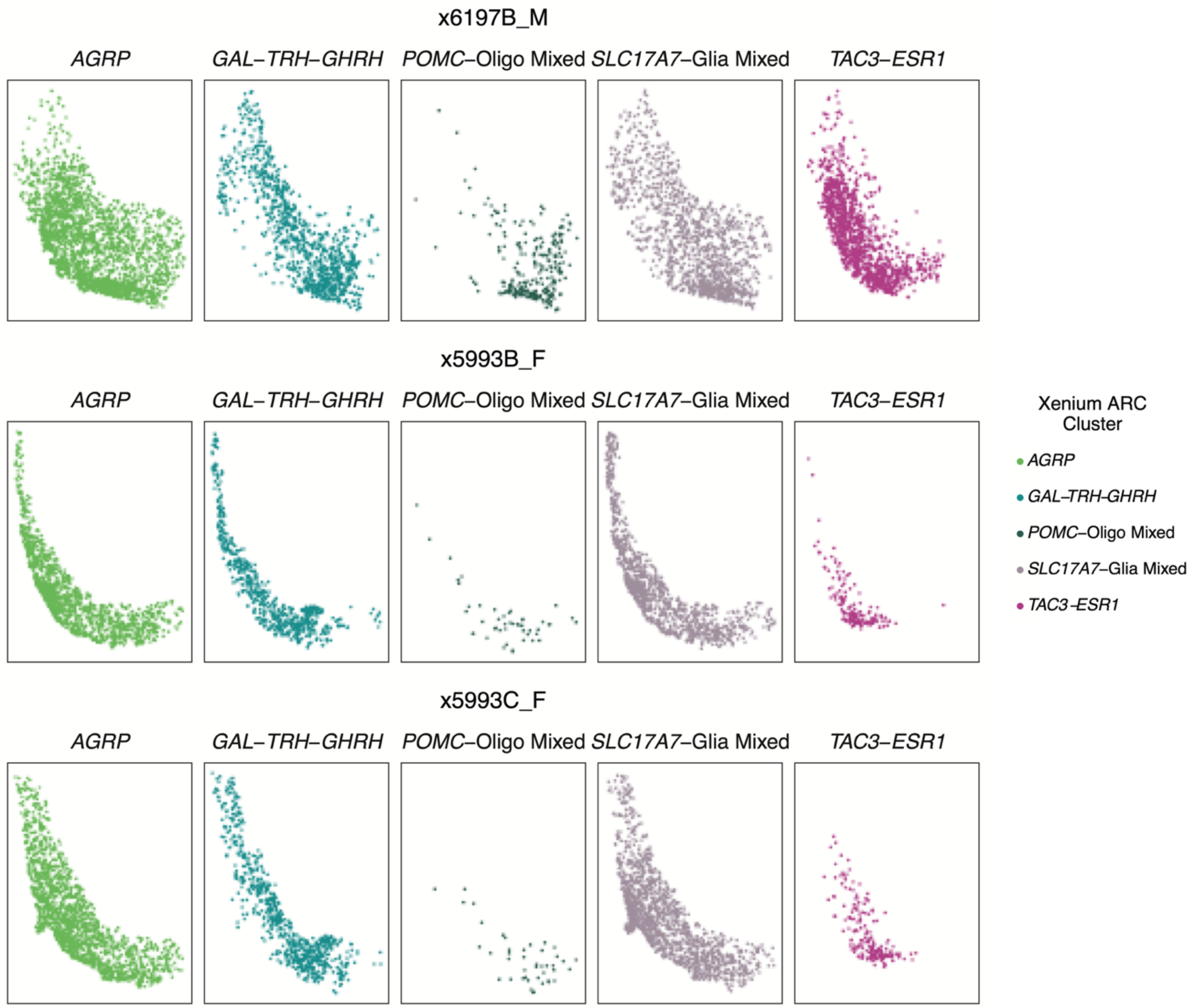


**Figure S23. Spatial organization of xARC cell clusters plotted within the xARC domain for all samples. (*Continued on next page*)**

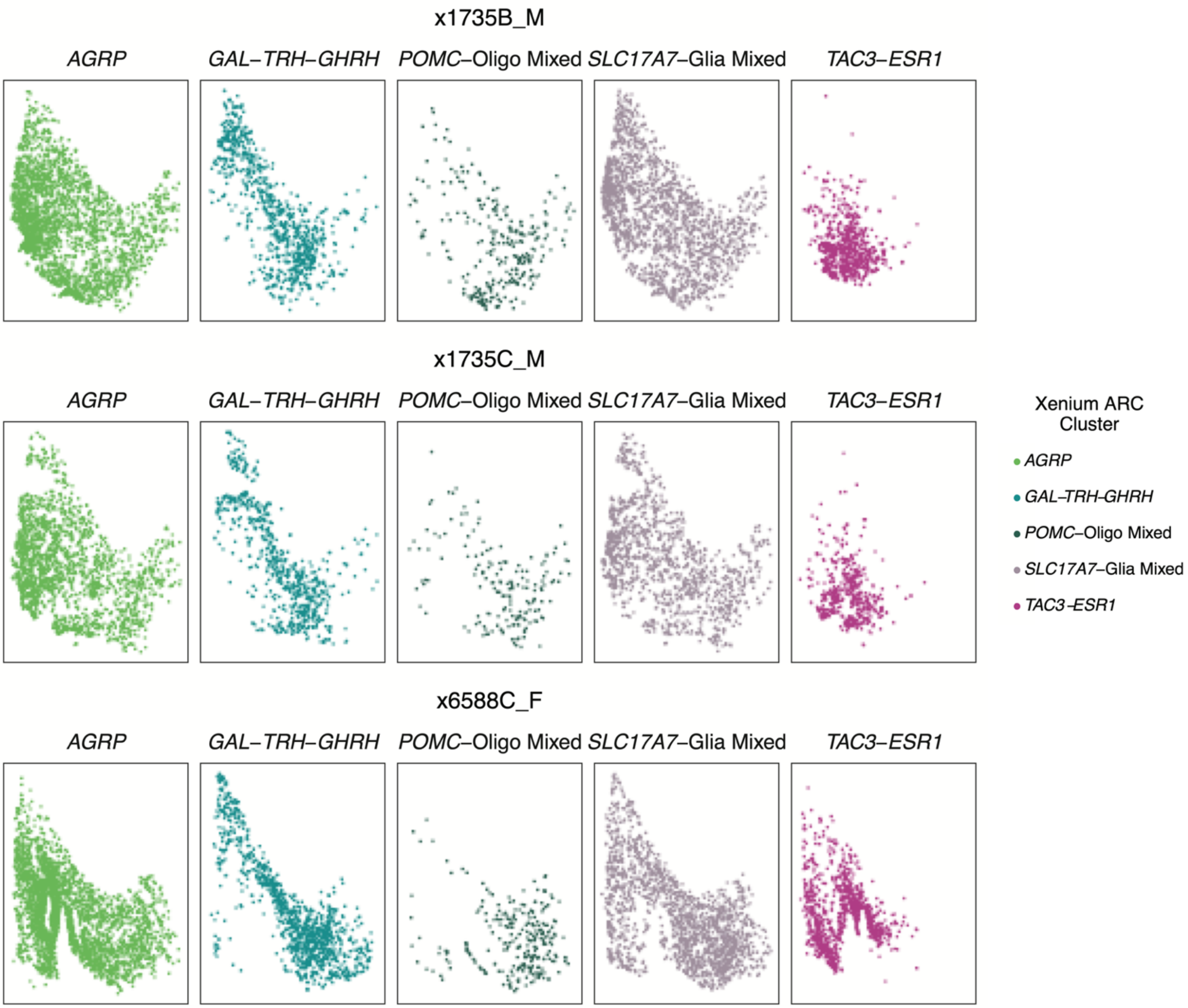


**Figure S23. Spatial organization of xARC cell clusters plotted within the xARC domain for all samples.** (***Continued on next page***)

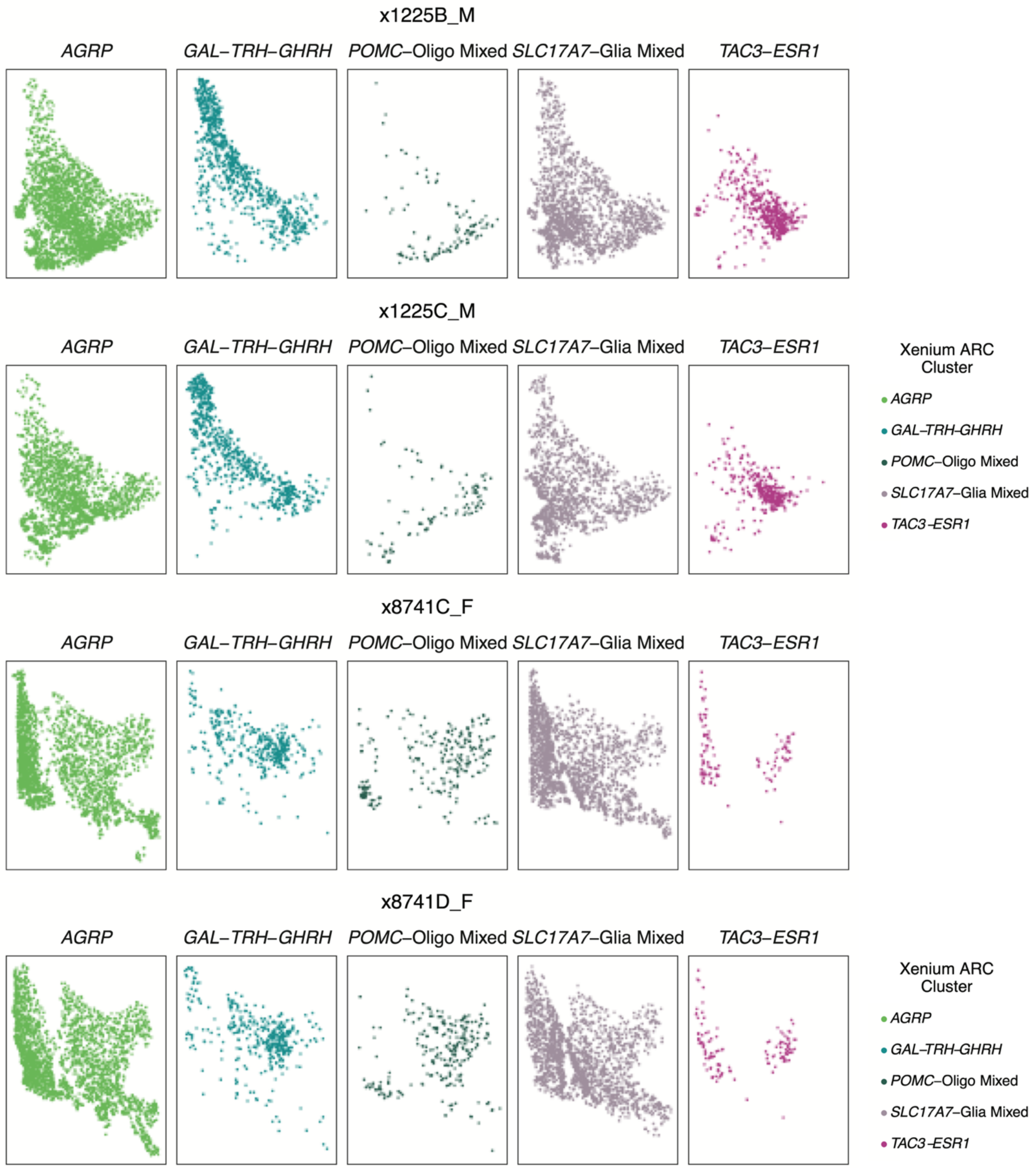


**Figure S23. Spatial organization of xARC cell clusters plotted within the xARC domain for all samples.** For each sample, plots illustrate the location of each cell from *AGRP*, *GAL-TRH-GHRH*, *POMC*-Oligo Mixed, *SLC17A7*-Glia Mixed, and *TAC3-ESR1* clusters. Only cells within the xARC domain area are plotted. Note that x8667B_F contains very little ARC tissue (similar to v8667A_F), and no *TAC3-ESR1* cells.

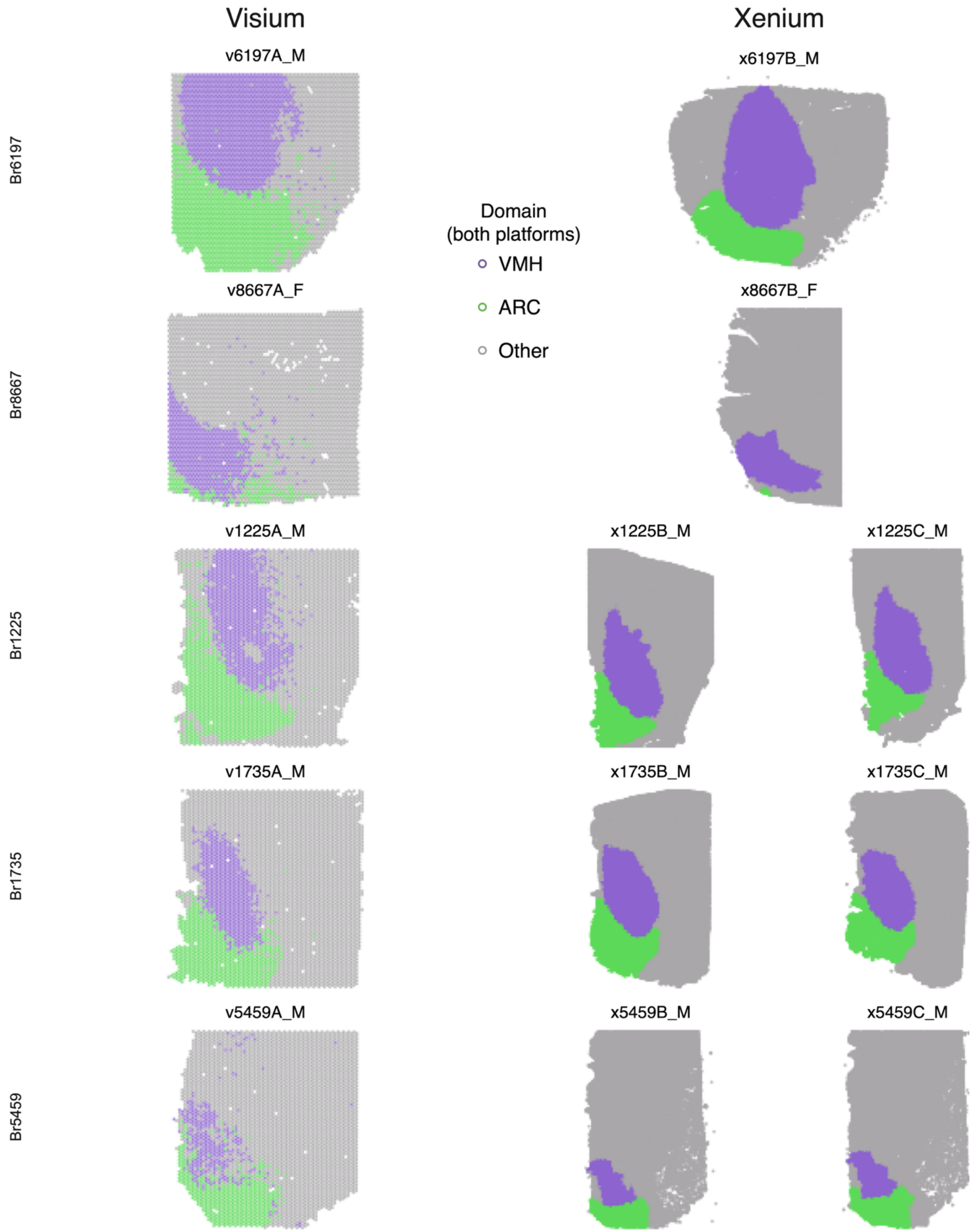


**Figure S24. Anatomically consistent VMH and ARC definitions across transcriptomic platforms. (*Continues on next page*)**

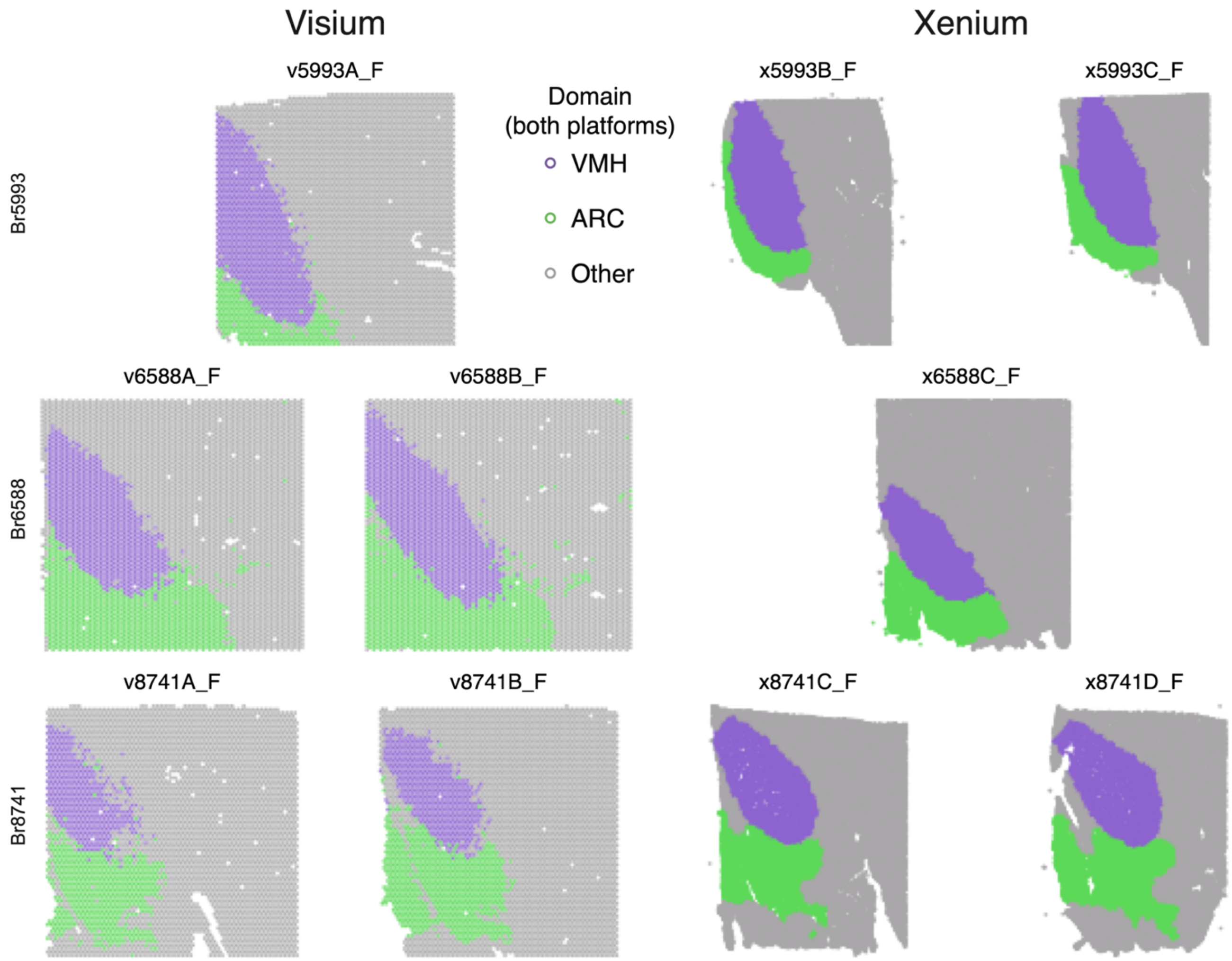


**Figure S24. Anatomically consistent VMH and ARC definitions across transcriptomic platforms.** Visium and Xenium samples are shown with spots or cells, respectively, colored according to their domain label. Left column displays Visium samples, and the right column Xenium samples. Each row shows all samples from a given donor (aspect ratios are not to scale). Sample nomenclature denotes "v" for Visium or "x" for Xenium, followed by donor ID (BrXXXX), "A" through "D" for tissue replicates, and "_M" for male or "_F" for female. Tissue samples from donor Br8667 (v8667A_F / x8667B_F) contained limited ARC. Aspect ratios are not to scale between platforms or among Xenium samples; several Xenium samples cover a larger dorsal-ventral span than is accommodated by Visium.

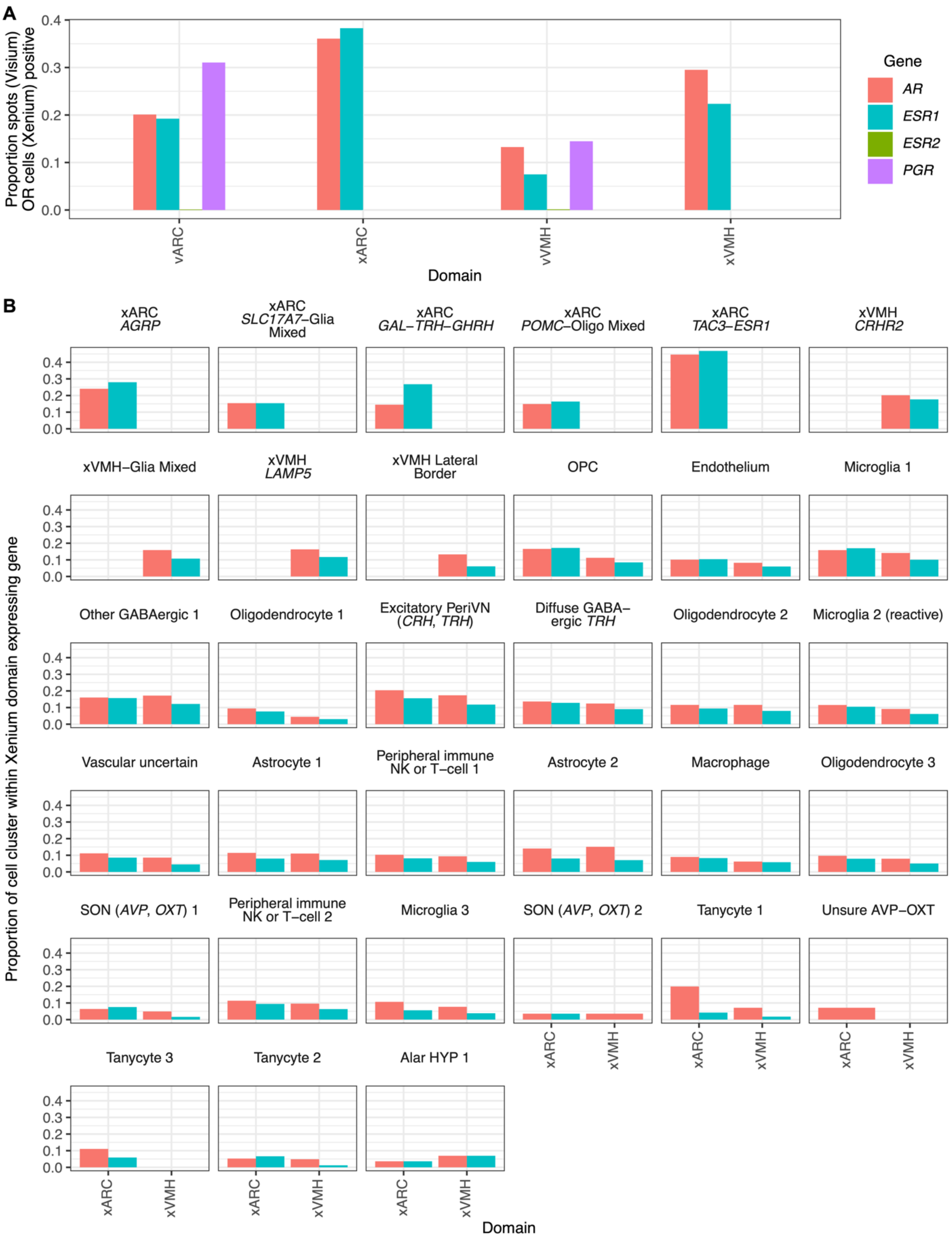


**Figure S25. Proportion of Visium spots and Xenium xVMH/xARC neurons expressing sex hormone receptors (*AR*, *ESR1*, *ESR2*, *PGR*) by domain or Xenium cluster-within-domain.** **A)** Proportion of spots vVMH/vARC spots (Visium) or xVMH- or xARC-specific neurons within their respective domains (Xenium) in which at least one count of a given gene (color-coded) was detected. *PGR* and *ESR2* were not assayed with Xenium. Note that *ESR1* and *AR* were both detected in a ~2-fold greater proportion of Xenium cells of a domain relative to Visium spots of that domain; this indicates improved detection of genes with generally low expression (*e.g.*, those encoding TFs) via imaging-based transcriptomics. **B)** Each plot shows the proportion of cells from a given Xenium cluster with at least one count of *AR* or *ESR1*. Only cells within the Xenium domain (*x*-axis) were considered.

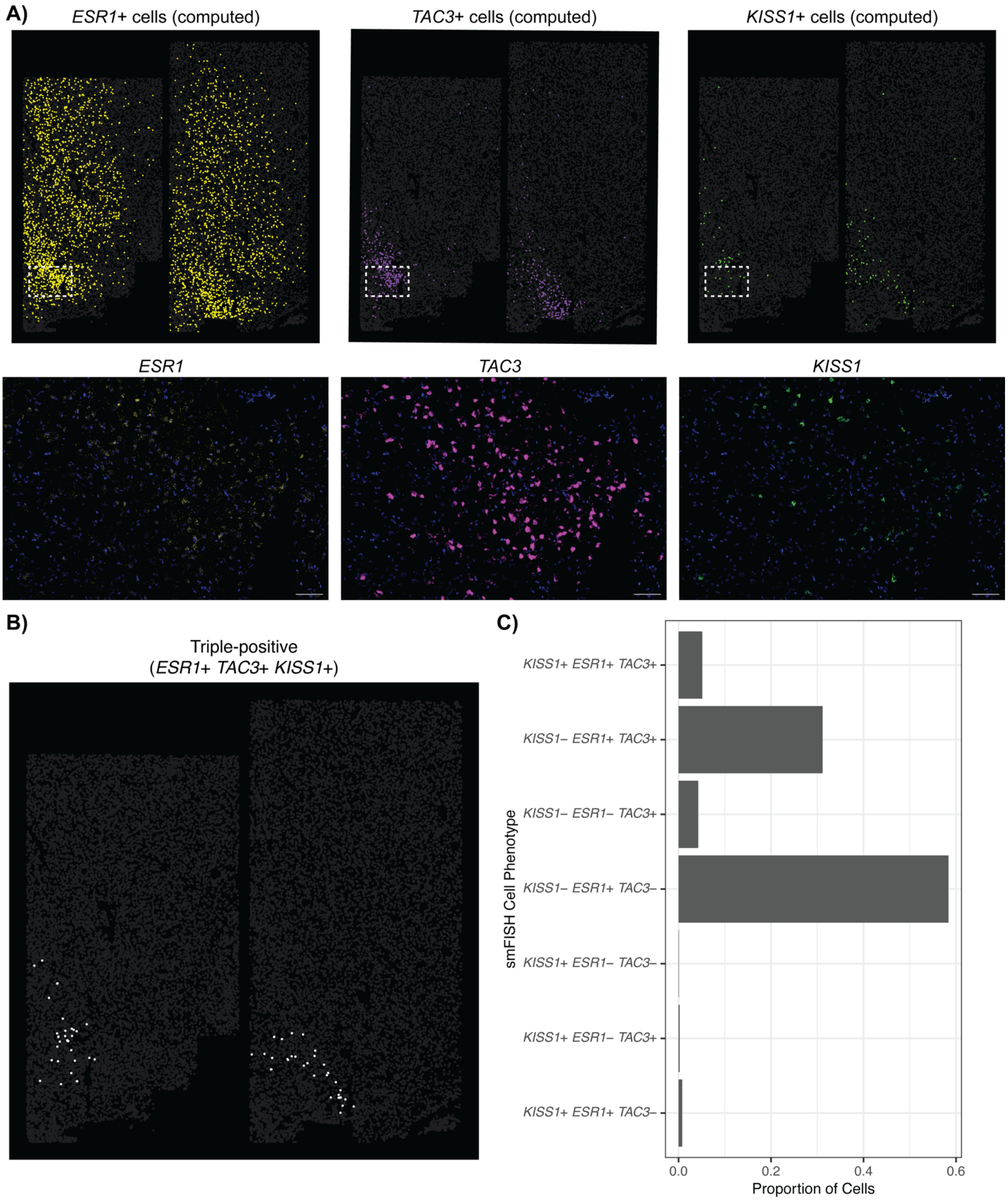


**Figure S26. VMH-ARC smFISH of *ESR1*, and *TAC3*, and *KISS1*. A**) Top row illustrates smFISH cells from two Xenium-adjacent serial tissue sections from donor Br1225 that were programmatically defined (*Methods*) as positive for the indicated gene (*ESR1*, *TAC3*, or *KISS1*). Boxes indicate the approximate tissue area shown below from raw fluorescence confocal images in the ARC region for the same gene (and DAPI). Scale bars 100µm. **B**) smFISH cells programmatically defined as triple-positive for *ESR1*, *TAC3*, and *KISS1*. **C)** Proportion of segmented cells in the ARC region of interest (*Methods*) across both tissue sections expressing ≥1 of *ESR1*, *TAC3*, or *KISS1*.

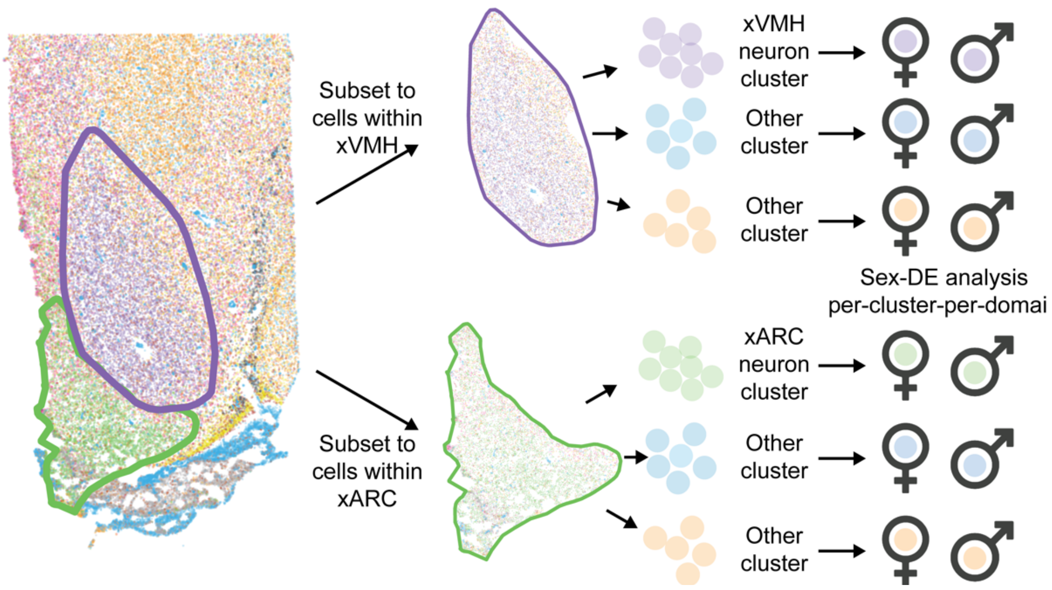


**Figure S27. Schematic overview of sex DE analysis procedure for Xenium clusters.** Sex-DE analysis was performed at the level of Xenium clusters by first subsetting to all cells within the boundaries of the xVMH domain or the xARC domain. Then, cells of each type are tested separately for sex-DE within that domain. Thus, broadly distributed cell clusters/types (e.g., glia) are tested for sex-DE in each domain. Meanwhile, sex-DE testing of domain-specific neuronal clusters (e.g., *TAC3-ESR1* in xARC) is filtered to the cells given that label *and* found in their domain (i.e. in the expected anatomic space). This simplified schematic only shows one domain-specific cluster each for xARC and xVMH.

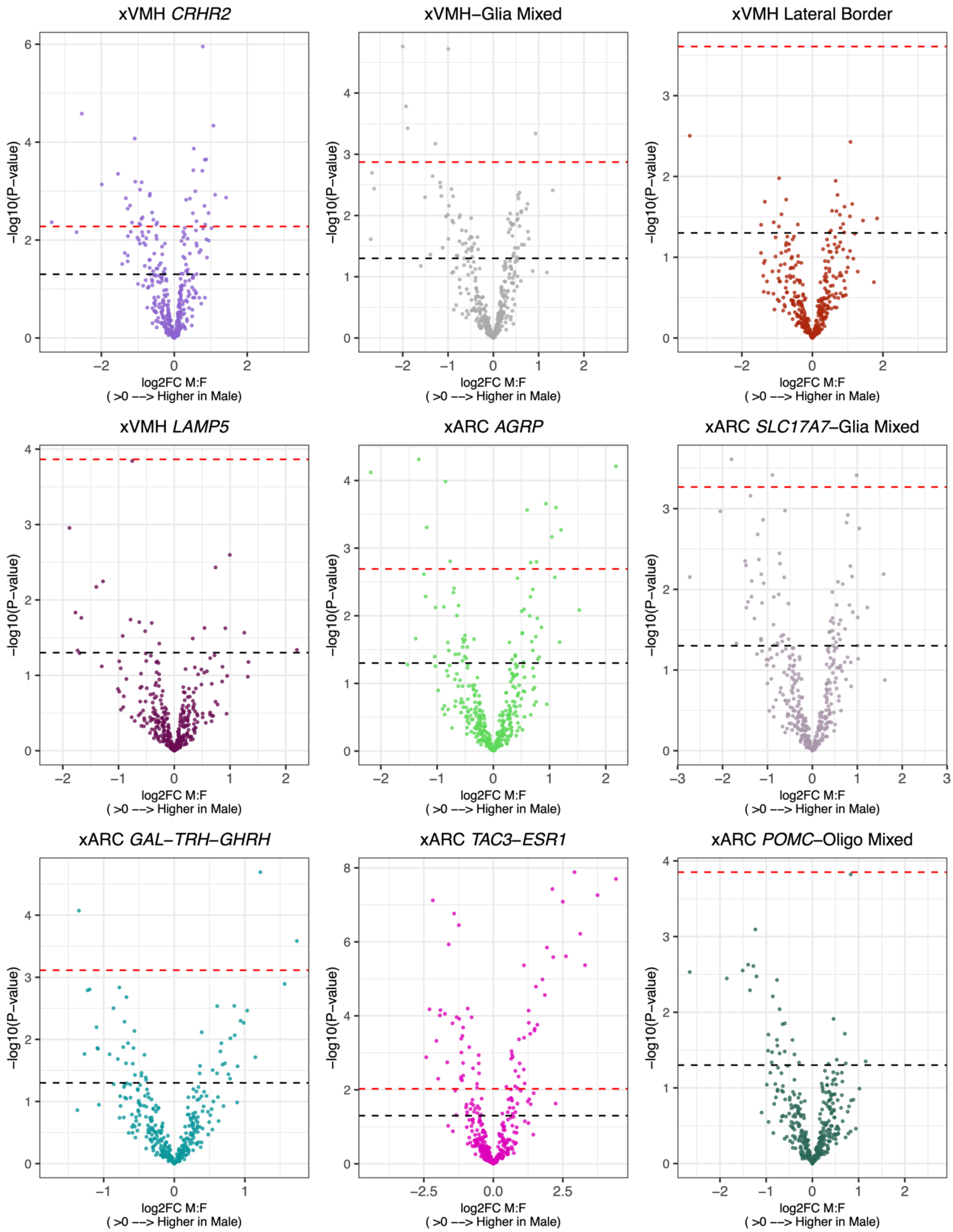


**Fig S28. Sex-DE volcano plots of the 5 xARC neuron clusters and the 4 xVMH neuron clusters.** Volcano plots are shown for each xARC/xVMH neuron cluster. Each cluster was analyzed only considering cells falling within their corresponding domain. Black and red dashed lines indicate cutoffs for $p<0.05$ and FDR<0.05, respectively.

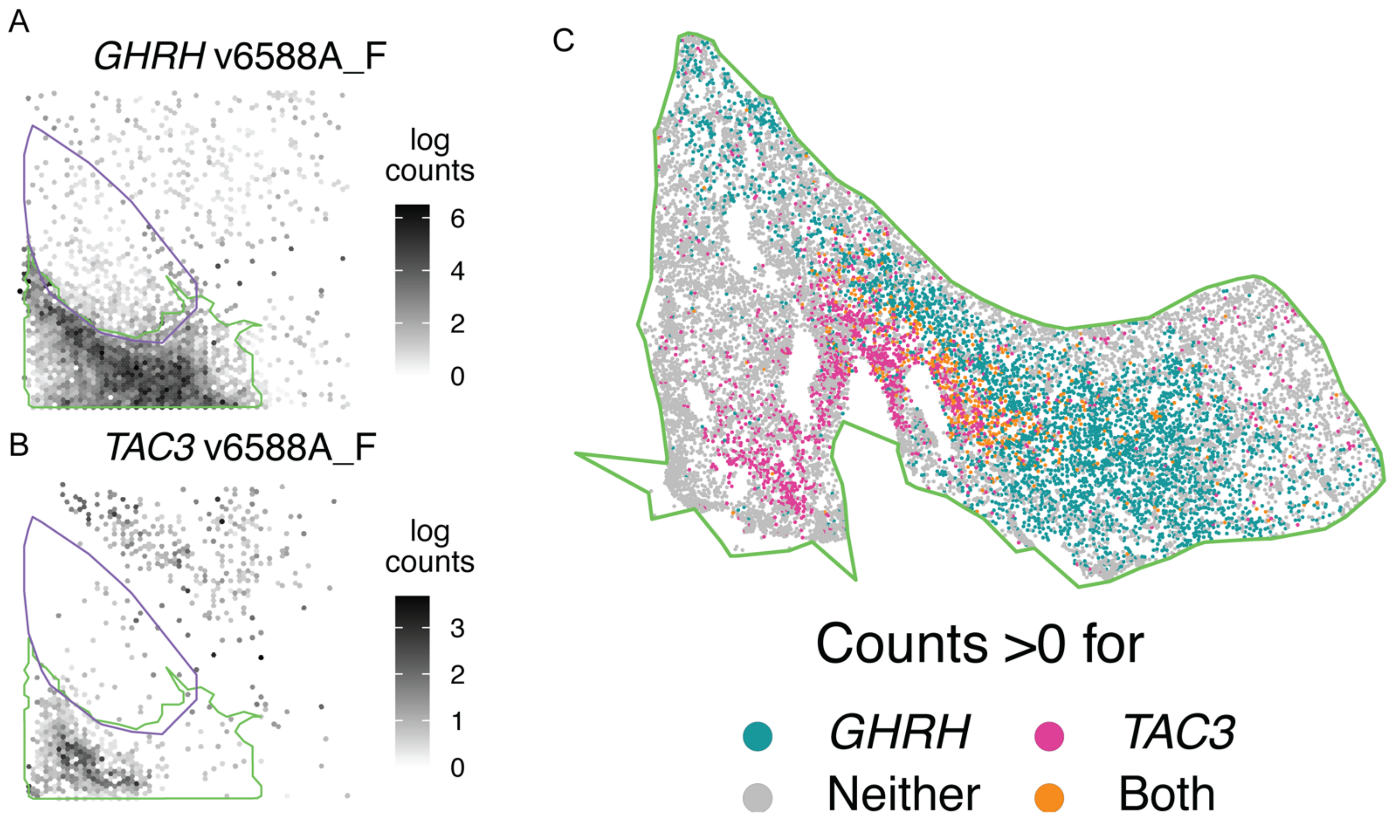


**Figure S29. Identification of spatially-restricted cell populations: selecting Xenium genes based on SVG analysis within vARC.** Analysis of spatially variable genes (SVGs) in Visium suggested spatial organization of cell types in ARC, exemplified by separate areas with peak expression of **A)** *TAC3* and **B)** *GHRH*. **C)** Expression of *TAC3* and *GHRH* in individual cells of xARC shown for x6588B_F (*i.e.*, from the same donor as panels A and B) measured by Xenium, confirming spatially-exclusive expression of these two genes.

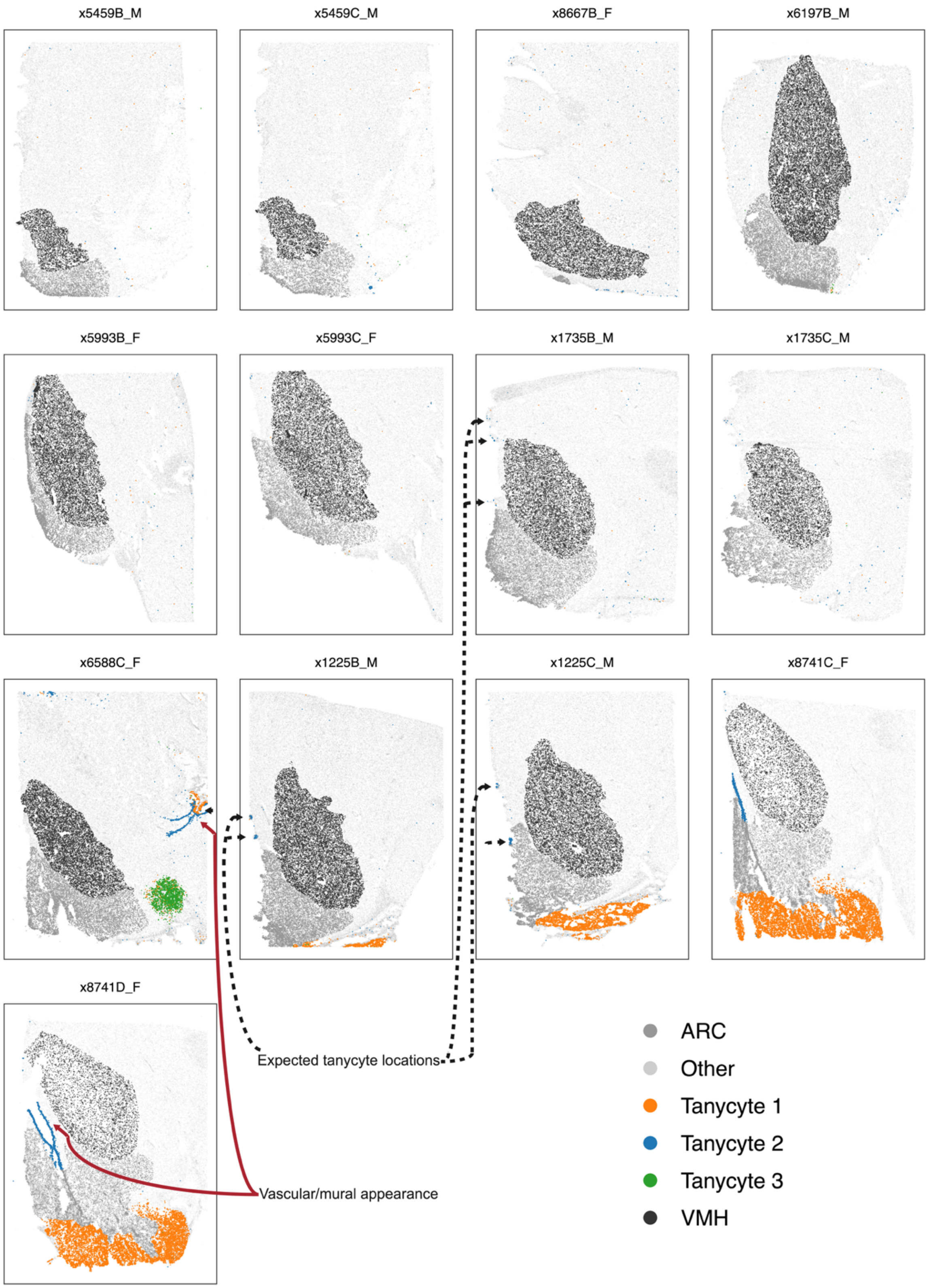


**Figure S30. Potential tanycyte clusters identified by Xenium variably map to anatomically-expected zones.** Each Xenium sample is shown, with cells in clusters including possible tanycytes colorized (legend, bottom right). Example areas with the cells in anatomically-expected locations and with the cells in arrangements resembling vascular/mural/connective tissue (*i.e.*, less likely to be bonafide tanycytes) are indicated with arrows.
